# A telomere to telomere assembly of *Oscheius tipulae* and the evolution of rhabditid nematode chromosomes

**DOI:** 10.1101/2020.09.04.283127

**Authors:** Pablo Manuel Gonzalez de la Rosa, Marian Thomson, Urmi Trivedi, Alan Tracey, Sophie Tandonnet, Mark Blaxter

## Abstract

Eukaryotic chromosomes have phylogenetic persistence. In many taxa, the number of chromosomes is related to the number of centromeres. However, in some groups, such as rhabditid nematodes, centromeric function is distributed across multiple sites on each chromosome. These holocentric chromosomes might, *a priori*, be expected to be permissive of large-scale chromosomal rearrangement, as chromosomal fragments could still partition correctly and fusions would not generate lethal conflict between multiple centromeres. Here, we explore the phylogenetic stability of nematode chromosomes using a new telomere-to-telomere assembly of the rhabditine nematode *Oscheius tipulae* generated from nanopore long reads. The 60 Mb *O. tipulae* genome is resolved into six chromosomal molecules. We find evidence of specific chromatin diminution at all telomeres. Comparing this chromosomal *O. tipulae* assembly with chromosomal assemblies of diverse rhabditid nematodes we identify seven ancestral chromosomal elements (Nigon elements), and present a model for the evolution of nematode chromosomes through rearrangement and fusion of these elements. We identify frequent fusion events involving NigonX, the element associated with the rhabditid X chromosome, and thus sex-chromosome associated gene sets differ markedly between species. Despite the karyotypic stability, gene order within chromosomes defined by Nigon elements is not conserved. Our model for nematode chromosome evolution provides a platform for investigation of the tensions between local genome rearrangement and karyotypic evolution in generating extant genome architectures.

## INTRODUCTION

Linear chromosomes are basic elements of the organisation of eukaryotic nuclear genomes. The number of chromosomes and the position of orthologous loci on them is generally conserved between closely-related species, and conserved karyotypic elements have been identified even between distantly related taxa (Nakatani et al., 2007; Putnam et al., 2008). The evolutionary trajectories of genes, in terms of rate of drift and efficiency of selection, are influenced by their chromosomal location. For example, genes on sex chromosomes will be exposed as haploid in the heterogametic sex (whether X0, XY or WZ), and their effective population size will be only 0.75 that of autosomal loci. More subtly, genes resident on longer chromosomes may be more affected by linked selection, as the number of recombination events is frequently limited to one per chromosome (Hammarlund et al., 2005) or chromosome arm, and the number of bases per centiMorgan will be larger in longer chromosomes. Gene evolution is also shaped by placement within chromosomes, with some regions, such as centromeres and subtelomeric regions experiencing higher rates of per-base and structural change (Rockman and Kruglyak, 2009). On longer timescales, genes that have travelled together on single chromosomes might evolve to share dependence on long-range regulatory landscapes, such as the three-dimensional topologically associated domains that characterise chromosomal organisation within interphase nuclei. For some sets of loci, such as HOX and paraHOX loci in most Metazoa, this constraint is evident between organisms that last shared common ancestors hundreds of millions of years ago (Krumlauf, 2018). Holocentric chromosomes offer a contrasting pattern of organisation to centromeric karyotypes, and this must affect the mode and tempo of genome and gene evolution.

Chromosome structural change is an important component of genome and species evolution (Sturtevant and Dobzhansky, 1936). Chromosomal elements, sets of loci that have been colocated on the same linkage group for long periods of evolutionary time, have been identified in many taxa, including mammals, Diptera (Bhutkar et al., 2008), Lepidoptera (d’Alençon et al., 2010) and Nematoda (Tandonnet et al., 2019). While many groups have deeply conserved karyotypes, species that have very different numbers of chromosomes or synteny relationships to closely related taxa will allow exploration of the constraints that act to retain chromosome number and gene content, and also of the mechanisms that are involved in karyotypic evolution.

Most animals (Metazoa) have chromosomes with a defined centromere. However several groups have holocentric chromosomes, where centromeric function is distributed across each chromosome and there is no single pairing centre during meiosis (Albertson and Thomson, 1982; Melters et al., 2012). *A priori*, holocentric organisation might be thought to predispose a genome to increased rates of rearrangement both within and between chromosomes, as any chromosome fragment remaining after fission could still carry centromeric function, and fusions of chromosomes would not result in competing centromeres on the same molecule. Lepidoptera have holocentric chromosomes and generally conserved karyotypes (d’Alençon et al., 2010). The ancestral lepidopteran chromosome number is estimated to be 31, and while the genomes of some species that have fewer chromosomes, such as the genus *Heliconius* (where n=21) can be modelled through a series of simple fusions (d’Alençon et al., 2010), others, such as *Pieris napi* (the green-veined white butterfly; n=25) exhibit extensive rearrangement, including presumed ancestral linkage group fragmentation and fusion (Hill et al., 2019). Thus, to distinguish conservation of chromosome number *per se* from conservation of linkage groups, and to define the patterns and processes involved in changes in karyotype, complete, telomere-to-telomere chromosomal assemblies are needed (Hill et al., 2019).

Nematode chromosomes are also holocentric. The model nematode *Caenorhabditis elegans* (Rhabditomorpha, Rhabditina, Rhabditida; see (De Ley and Blaxter, 2002)) has n=6 and an X0 sex determination mechanism. In the order Rhabditida, n=6 is the commonest karyotype (Table S1) (Walton, 1959),, but n varies between 1 (e.g. *Diploscapter coronatus*, closely related to *Caenorhabditis*) (Fradin et al., 2017) and >50 (e.g. *Meloidogyne* polyploids) (Triantaphyllou, 1963). Conservation of karyotype implies that sets of genes will be colocated on the same chromosome for long evolutionary periods. The retention of syntenic groups of loci can be uncoupled from retention of large-scale gene order. In *Caenorhabditis* species (all with n=6) orthologous genes are overwhelmingly located on orthologous chromosomes but the order of genes is very different between species due to rampant within-chromosome rearrangement (Stein et al., 2003; Stevens et al., 2020; Teterina et al., 2020). This pattern is also seen when comparing *Caenorhabditis* to other genera: linkage groups tend to be conserved but gene order is not (Doyle et al., 2019). In contrast, both linkage groups and gene order within each linkage group are conserved in the similarly holocentric Lepidoptera (Mongue et al., 2017).

Some nematodes have different karyotypes in their somatic cells compared to their germline. This process involves scission of germline chromosomes and loss of germline material, and is called chromatin diminution (Wang and Davis, 2014). Chromatin diminution has been observed in several metazoan taxa, including chordates (Kinsella et al., 2019) and arthropods, and is involved in the generation of the ciliate macronucleus (Rzeszutek et al., 2020). The process of diminution is best understood in *Ascaris suum* (Ascarididomorpha, Spirurina, Rhabditida), where the germline has n=24 chromosomes but somatic cells have n=36 (Wang et al., 2020). In *A. suum* the breakage events affect some but not all of the X chromosomes (*A. suum* has five X chromosomes) and autosomes, and breakage and neo-telomere addition happens in a defined area of the chromosome, but not at a precise base position. Related ascarididomorph nematodes also display diminution. Chromatin diminution has also been described in the tylenchomorph nematode *Strongyloides papillosus,* where loss of a specific internal fragment of one copy of the X chromosome generates a haploid region that is associated with males (i.e. sex determination in *S. papillosus* is effectively XX:X0, but the nullo-X is determined through specific deletion).

Previously we proposed the existence of seven ancestral chromosome elements in rhabditine nematodes, named Nigon elements, and used this model to understand chromosome evolution in a few genome-sequenced Rhabditina (Tandonnet et al., 2019). The lack of chromosomally-complete genomes limited the power of the model. Here we present an improved, chromosomal genome assembly of *Oscheius tipulae* (Rhabditomorpha, Rhabditina, Rhabditida). *O. tipulae* is a satellite genetic model organism that is used to understand the evolution of developmental systems such as the specification of the nematode vulva, and the genome sequence is required to underpin detailed genetic mapping (Besnard et al., 2017). The new telomere-to-telomere assembly allowed us to identify unexpected features of chromatin diminution at the telomeres of each chromosome. We used the *O. tipulae* genome and other chromosomally-complete nematode genomes to fully define sets of orthologous genes associated with ancestral Nigon elements. This analysis allowed us to map ancient chromosomal fusions and scissions and identify a set of genes that is always associated with the X chromosome in both X0 and XY taxa in the order Rhabditida.

## METHODS

### Nematode culture, DNA extraction and QC

*Oscheius tipulae* strain CEW1 (Evans et al., 1997) was obtained from Marie-Anne Félix (Institute of Biology of the Ecole Normale Supérieure, Paris), and cultivated at 20°C in 5 cm nematode growth medium lite plates seeded with *E. coli* HB101 (Stiernagle, 2006). Nematodes were washed from culture plates using an M9 buffer supplemented with 0.01% Tween 20. Nematodes were pelleted by low-speed centrifugation, and 100 µL samples transferred with minimal supernatant to 1.5 mL LoBind Eppendorf tubes. Nematodes were lysed by addition of 600 µL of Cell Lysis Solution (Qiagen) and 20 µL of proteinase K (20 µg/µL) and incubated at 56 °C with mixing at 300 rpm for 4 h. RNA was digested by adding 5 µL of RNAse Cocktail Enzyme Mix (Invitrogen) and incubating at 37°C for 1 h. Protein was precipitated by adding 200 µL of ice-cold Protein Precipitation Solution (Qiagen), gentle mixing and incubation on ice for 10 minutes. The precipitate was pelleted by centrifugation for 30 min at 4°C at 15,000 RPM. The supernatant was transferred to a LoBind tube and nucleic acids precipitated by addition of 600 µL of ice cold isopropanol, mixing by inversion, and incubation on ice for 10 min. Nucleic acids were pelleted by centrifugation at 4°C at 15,000 RPM. The supernatant was discarded and the pellet washed twice using 600 µL of 70% ethanol. The pellet was air dried for 5 min and resuspended in 20 µL of Elution Buffer. DNA recovery and quality was assessed by Qubit fluorimetry, Tapestation genomic Screentape (Agilent) and pulsed field gel electrophoresis using a Pippin Pulse instrument. The sample used for sequencing had a DNA concentration of 120 ng/µL, a DNA integrity number of 9 and an RNA concentration of 9 ng/µL.

### Genomic sequencing on Oxford Nanopore PromethION

To generate fragments of a suitable size range for Oxford Nanopore PromethION sequencing, high molecular weight DNA was diluted to a concentration of 25 ng/uL and fragmented to an average peak size of 25 kb using a Megaruptor-2 instrument (Diagenode). Small fragments <1 kb were removed and the DNA concentrated using bead purification (0.4 x volumes of Ampure-XP beads. Two aliquots of 1 µg of sheared *O. tipulae* DNA and control DNA (lambda 3,2 kb fragment) were subjected to DNA damage repair (NEBNext FFPE DNA Repair Mix; New England Biolabs) followed by DNA End Repair (NEBNext Ultra II End Repair/ dA-tailing Module; New England BioLabs). A second 0.4 x volume Ampure-XP bead clean up was carried out and DNA eluted in sterile distilled H_2_O. Oxford Nanopore sequencing adapters were ligated to 750 ng of the recovered, end repaired DNA using the Ligation Sequencing kit (SQK-LSK-109; Oxford Nanopore) and NEBNext Ligation Module (New England BioLabs). Following a further 0.4 x volume Ampure-XP purification, the recovered DNA (16.25 fmol) was loaded onto a R9.4.1 PromethION flow cell following the manufacturer’s instructions and a 60 hr sequencing run was initiated.

Raw reads were basecalled using Guppy (see Table S2 for software tools and settings used). The resulting dataset of 8.8 M reads spanned 108.4 Gb and had a read N50 of 19.1 kb (Figure S1 A; Table S3). To identify sequence contamination, we assembled a custom kraken2 database composed of bacteria, fungi, human, UniVec core and a selection of nematode genomes including the previous *O. tipulae* assembly (Table S4). The vast majority of reads (99.5%) were classified. One fifth (19.1% of the total bases) were classified as Nematoda, and the remainder as Proteobacteria (97.5% of these belonging to *Escherichia*; these likely derive from bacterial food). Minor human, fungal and other bacterial contamination was also present (Table S5). We removed reads classified as Bacteria, Chordata, Ascomycota, Basidiomycota or Microsporidia. The remaining data spanned 20.7 Gb in 2.8 million reads with a read N50 of 14.4 kb (an estimated ∼340 fold coverage).

### Genome assembly and polishing

Several different assembly strategies were explored (Table S6). Flye (Kolmogorov et al., 2019) in metagenome mode with the whole long read set yielded a chromosome level assembly of *O. tipulae* together with contaminant species (Figure S1 B). This assembly was polished using Racon (Vaser et al., 2017) and medaka using the decontaminated read set. For Pilon polishing (Walker et al., 2014), Illumina reads were trimmed with BBDuk (Bushnell, 2017) and aligned with BWA-MEM (Li and Durbin, 2009). We derived the chromosome assembly nOti 3.1 by stitching back the two sequences of chromosome I with RaGOO (Alonge et al., 2019) using the unpolished assembly as a reference. An alternate assembly was obtained using Flye in metagenome mode with only 40x Canu-corrected, decontaminated reads, followed by Racon and Pilon polishing. This assembly had higher BUSCO completeness than nOti 3.1, but was more fragmented. We derived a new consensus, nOti 3.2, from nOti 3.1 and the decontaminated-read Flye assembly using gap5 (Bonfield and Whitwham, 2010) giving the decontaminated read assembly a 100x relative weight. The resulting contigs were assigned chromosome names by the longest match to contigs in nOti 2.0, which were previously assigned to chromosomes (Besnard et al., 2017). Alignment of the raw reads against nOti 3.2 showed that the nuclear genome had an average per-base coverage of 334 fold (standard deviation of 198). The initial Oti_chrV sequence had the highest coverage and coverage heterogeneity (354 fold, SD 478) due to collapse of the ribosomal RNA cistron between positions 7413040 and 7440271 (see below; Table S7).

We curated the nOti 3.2 assembly by examining read coverage across the genome. A gap5 (Bonfield and Whitwham, 2010) database was built from a 200x sub-sample of the longest PromethION canu corrected reads. We noticed that all the chromosomes were characterised by a shorter majority sequence (80% of the average coverage depth) and a longer minority sequence (20% coverage depth). Both of these sequences terminated in telomeric repeat (long tandem repeats of TTAGGC). These alternate telomeric repeat addition sites appeared not to be artefacts because long reads supporting both versions were anchored in unique sequence. Previous Illumina short read data also identified similar major and minor components of the chromosome ends, and supported the same telomeric repeat addition sites. We manually extended all reads containing soft-clipped telomeric repeat sequence and then used the gap5 realign function to produce a new consensus from them. The left hand end of the Oti_chrIV sequence produced directly by the assembler was characterised by an artificial sequence as evidenced by a lack of reads that mapped to it and all reads being soft-clipped either side of it. This sequence was replaced with realigned, soft-clipped sequence from the adjacent mapped reads. Restoration of these soft-clipped data identified telomeric repeat at both ends of each chromosome. We estimated the size of the highly collapsed ribosomal RNA cistron repeat on Oti_chrV. The rRNA cistron repeat was estimated to be 6.8 kb long and to be present in 117 copies, based on its coverage by Illumina short reads. The left hand side of the rRNA cistron repeat terminated in an obviously mispredicted sequence which was dealt with as for the Oti_chrIV telomere sequence. We extended the reads at the junctions into and out of the rRNA cistron repeat sequence to a minimum depth of 2 reads. We joined these flanks together using 760128 “N” characters to match 111 additional copies of the 6.8 kb rRNA cistron repeat. We polished this assembly via three rounds using freebayes through snippy (Seemann, 2014) with Illumina reads. We identified spliced leader RNA (SL) and 5S rRNA loci using Rfam models (Nawrocki et al., 2015) (Figure S2).

### Genome annotation

We created a repeat library for our assembly following published protocols (Coghlan et al., 2018). Briefly, we identified repetitive sequences with RepeatModeler2 (Flynn et al., 2020), transposons with TransposonPSI (Haas, 2007) and Long Terminal Repeats (LTR) with LTRharvest (Ellinghaus et al., 2008). We discarded TransposonPSI predictions shorter than 50 bp. We filtered out LTRharvest predictions that lacked PFAM (Finn et al., 2016) and GyDB (Llorens et al., 2011) hidden Markov model domain hits using LTRdigest (Steinbiss et al., 2009). The three prediction sets were classified with RepeatClassifier and merged into a single library. Sequences were clustered if they had more than 80% identity using USEARCH (Edgar, 2010). This repeat library, together with the CONS-Dfam_3.1-rb20181026 database (Hubley et al., 2016), was used by RepeatMasker to annotate the repetitive regions in the assembly. We predicted protein coding genes using GeneMark-ES (Lomsadze et al., 2005). We explored helitron predictions using HelitronScanner (Xiong et al., 2014). For nucleotide and protein-coding gene comparisons in the telomere extensions we used BLAST+ (Camacho et al., 2009), ClustalW (Thompson et al., 2002) and JalView (Waterhouse et al., 2009). We identified genes and other sequence features of telomeric extensions using BLAST similarity searches and hidden Markov model searches using Dfam helitron models (Wheeler and Eddy, 2013). Images were generated using the circos toolkit (Krzywinski et al., 2009).

### Comparison to genomes of other Rhabditida and identification of loci supporting Nigon elements

We assessed chromosomal assemblies and annotations of fifteen rhabditid nematode species (Table S8, Table S9). We extracted the longest protein per gene from genome annotation GFF3 files using AGAT (Dainat, 2020). Orthogroups were identified by Orthofinder (Emms and Kelly, 2019, 2015) using these proteomes. Orthogroups were filtered using Kinfin (Laetsch and Blaxter, 2017) to identify fuzzy single copy orthologues in at least 12 of the 15 species compared (see Supplementary Text S1).

We identified loci that are found on the same chromosomal unit in multiple species using a subset of nine genome assemblies that are resolved into chromosomes: *Auanema rhodensis, Brugia malayi, Haemonchus contortus, Meloidogyne hapla, Onchocerca volvulus, Oscheius tipulae, Pristionchus pacificus, Steinernema carpocapsae* and *Strongyloides ratti*. For *Meloidogyne hapla*, contigs were grouped into chromosomes according to the genetic linkage map (Opperman et al., 2008). For *Steinernema carpocapsae*, only the X chromosome was assembled to completeness, while the four autosomes were present as twelve unassigned scaffolds. To reduce phylogenetic bias we used a single *Caenorhabditis* species (*C. elegans*).

The chromosomal location of each single copy orthologue in each species was extracted from genome GFF3 files and collated in an orthologue-chromosomal allocation matrix by species. Scaffolds “a” and “b” of *O. volvulus* chromosome 1 were treated as a single chromosome, as were the *S. ratti* X chromosome scaffolds. We calculated the Dice distance (represented as 1-Dice distance) between all orthologue pairs based on the pattern of their chromosomal allocation between species. A pair of genes found on the same chromosome in all the species would have a similarity of 1, while a pair found on different chromosomes in all the species would have a similarity of 0. This matrix was clustered with CLARA (Clustering Large Applications) using an expected number of clusters, k, from 1 to 10 (Figure S3). CLARA identified the medoids (center of the clusters) by applying PAM (partitioning around medoids) to 5 independent subsamples of 10% of the orthologous loci. To identify the best number of clusters we assessed average silhouette values which will be higher when the average loci is closer to members of the same clusters than to loci of other clusters. The highest silhouette value was found at k=7 when using CLARA or PAM clustering (Figure S3 A and D). CLARA was used because it reduced computation time. Similarly, 7 clusters were identified by the t-Distributed Stochastic Embedding (t-SNE) plot. These clusters were also found when using different perplexity values, which can be interpreted as the number of effective nearest neighbors, from 30 to 1000 (Figure S4). Orthologues were assigned to clusters as long as at least 7 taxa agreed on their colocation with other orthologues in the cluster. Cluster numbers were assigned to putative element groups when they contained more than 20 single copy orthologues, and allocated to Nigon element labels according to our previous classification (Tandonnet et al., 2019).

### KEGG pathway enrichment analysis

We assessed functional enrichment of KEGG pathways among each Nigon defining loci set using the *C. elegans* representatives through gProfiler (Raudvere et al., 2019). We downloaded the *C. elegans* genes KEGG annotations using the R package KEGGREST (Tenenbaum, 2019). We used a Fisher exact test controlling the false discovery rate by Benjamini-Hochberg p-value correction to assess pathway enrichment using the 2175 *C. elegans* Nigon defining loci as comparator.

### Data availability

The raw PromethION data are available in INSDC under accession SRR12179520 associated with the BioSample SAMN15480678. The genome assembly has been deposited in INSDC with accession number GCA_013425905.1. The data used to generate the Tables and Figures are available at https://docs.google.com/spreadsheets/d/1gO4j4jSgSYQ_Aofl59RdGbwHrz2L2loWfrDIyyRaohw/edit?usp=sharing. Scripts and intermediate files associated with this study are available in https://github.com/tolkit/otipu_chrom_assem (commit dc73c57) under a GPL-3.0 License.

## RESULTS

### A chromosomal assembly of *Oscheius tipulae* CEW1 from Oxford Nanopore PromethION data

Our previous assembly of *O. tipulae* CEW1 had a contig N50 less than 1 Mb, and was resolved to chromosomes using genetic map data, which necessarily excluded contigs with no mapped loci and could not orientate contigs mapped through a single genetic locus (Besnard et al., 2017). To generate a telomere-to-telomere, chromosomally complete *O. tipulae* genome we resequenced CEW1 using the Oxford Nanopore Technologies PromethION platform, which generates large numbers of long reads from single molecules. After removing reads from bacterial contamination, we retained approximately 340 fold coverage of the expected 60 Mb genome (Table S5). We explored assembly options using a range of tools, and polished the assemblies to correct remaining errors with previously obtained Illumina short read data. Twelve assemblies were generated, with nematode sequence contiguities ranging from 7 to over 900 contigs (Table S6). The most contiguous assemblies had seven contigs, six with multi-megabase lengths and one corresponding to the mitochondrial genome. Each of the assemblies contained about 90% of the BUSCO set of conserved nematode orthologues. An assembly generated using Flye (Kolmogorov et al., 2019) in metagenome mode, combining data from decontaminated and full read sets, scored best using BUSCO and contiguity measures. We curated the genome by circularising the mitochondrion, and identifying and correcting two remaining issues in the nuclear sequence. The ribosomal RNA repeat cistron was present as a collapsed and jumbled sequence, as was the case until recently with the *C. elegans* complete genome sequence. A gap representing the estimated span of the repeat was inserted. While the 5S rRNA and spliced leader 1 RNA loci are present as a tandem repeat in *C. elegans* and related nematodes, the 5S and SL1 RNA genes were not clustered in *O. tipulae* (Figure S2).

The second issue concerned the chromosomal termini. Only four of the twelve ends of contigs in the initial assembly ended in telomeric repeat sequence ([TTAGGC]_n_, the same repeat as found in *C. elegans*). We were able to extend each contig end into telomeric repeat using long reads that overlapped the unique sequence at each end of each contig. The assembly was thus judged complete, telomere-to-telomere. The six longest contigs corresponded to the six genetically-defined linkage groups of *O. tipulae*, and we named these Otip_I, Otip_II, Otip_III, Otip_IV, Otip_V and Otip_X, following the previously defined chromosome nomenclature (Besnard et al., 2017). However, at each telomere we identified two independent and specific sites where there was transition from unique sequence to tandem hexamer telomere repeat: an internal site supported by 80% of the PromethION reads, and an external site, supported by a minority (20% of reads). The extension sequences were confirmed by relatively even, 60-fold coverage of PromethION reads and by matching reads in Illumina short-read data, which also showed a majority-short and minority-long distribution (Figure 1). These telomere repeat addition sites define three components on each chromosome: a central portion, present in all copies, and two sub-telomeric extension sequences, from the regions at the left and right ends, present in a minority of copies. In turn this implies that *O. tipulae* chromosomes are each found as long-form (carrying the extensions) and short form (lacking the extensions) versions. We cannot exclude the possibility that absence of the extensions on the left and right ends of each chromosome is determined independently, but the very similar proportional coverage of all extensions strongly suggests that chromosomes are either all short form or all long form in each cell.

**Figure 1:**
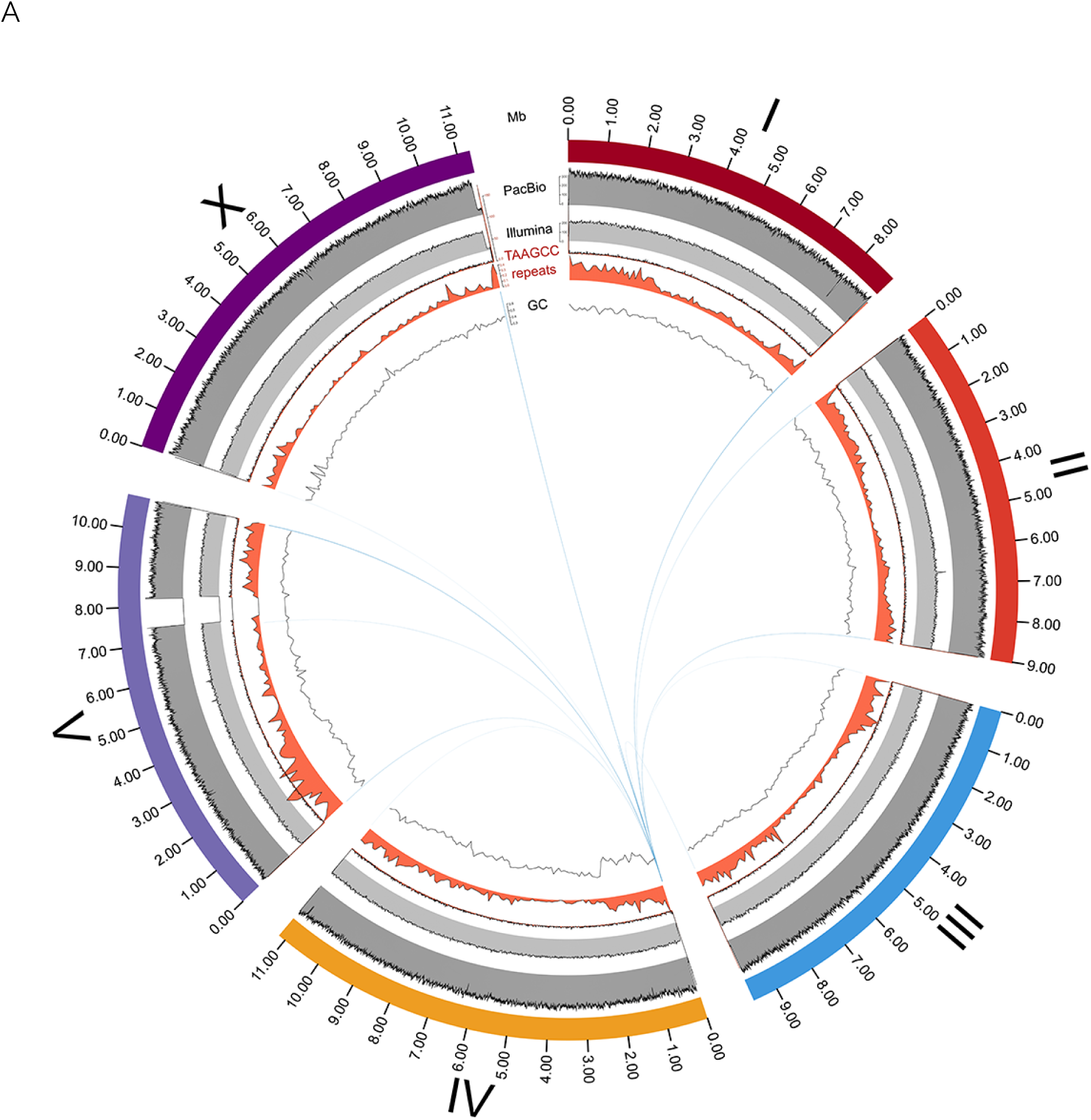
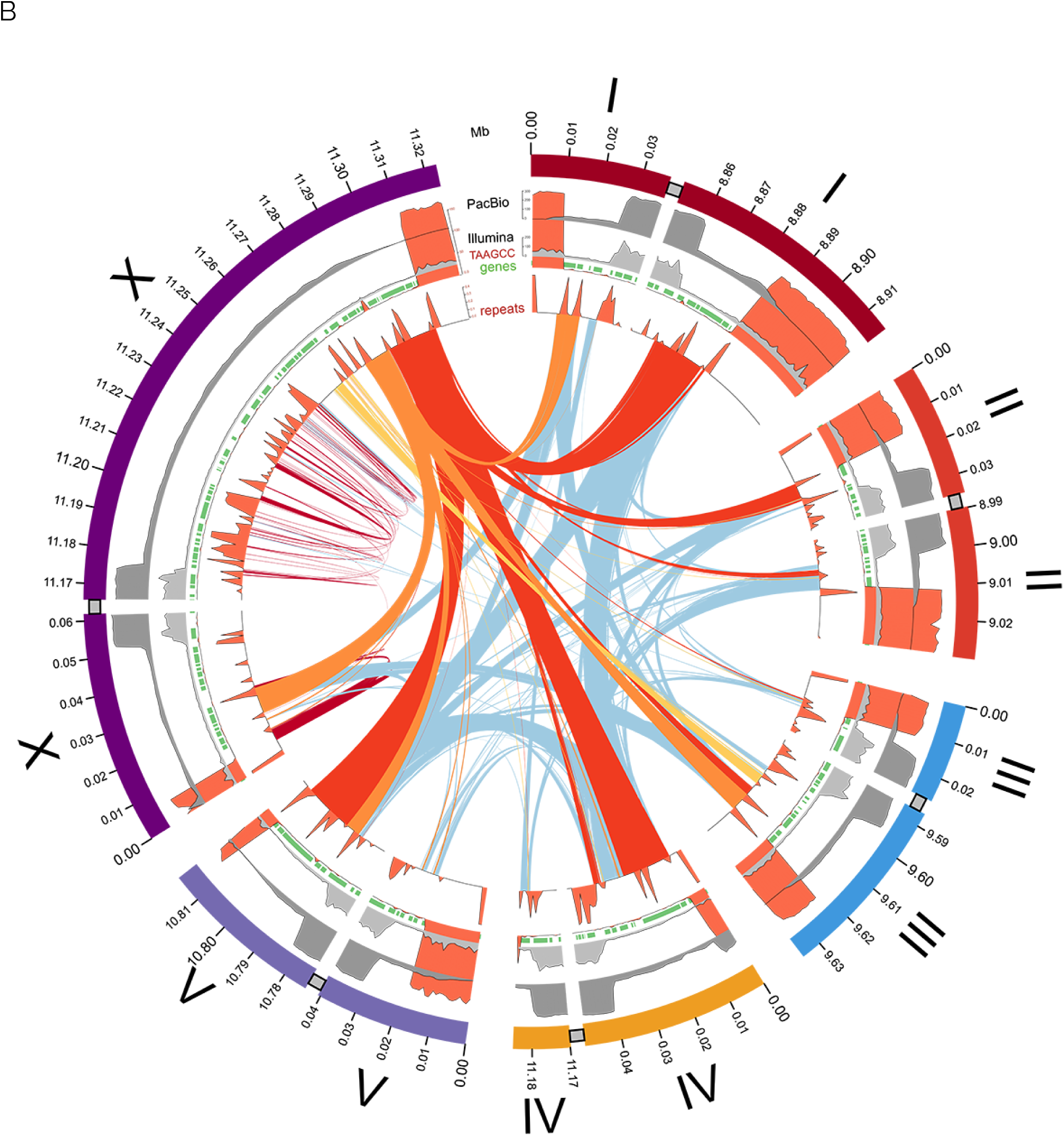
Dual telomeres on the chromosomes of *Oscheius tipulae* CEW1. A Circos plot of the *O. tipulae* CEW1 genome showing (from outside to inside) the chromosomes with a scale in Mb, coverage in PromethION reads (dark grey), coverage in Illumina short reads (light grey), count of telomeric repeats per 10 kb (red), density of repeats (red) and GC proportion in 10kb windows (line plot). The inner arcs link all significant nucleotide sequence (blastn) matches between the presumed near-complete copy of the helitron on the left end of Otip_IV (with an open reading frame of 3895 amino acids, spanning 26 kb of the telomere extension) and the rest of the genome. B Circos plot of the low-coverage telomeric extensions of *O. tipulae* CEW1, with 10 kb of normal coverage flanking chromosome, showing (from outside to inside) the chromosomes with a scale in 100 kb, coverage in PromethION reads (dark grey), coverage in illumina short reads (light grey), count of telomeric repeats per 10 kb (red), density of repeats (red) and predicted coding genes (green). Arcs are drawn linking significant blastn matches between three sequential helitron-like components from the right telomere extension of chromosome X (yellow, orange and red links), between repeats limited to within Otip_X R (dark red), and between all other telomeric extension sequences (light blue).

The sub-telomeric extensions ranged from 4 kb to 133 kb (excluding the telomeric hexamer repeats), and totaled 349 kb (Table 1; Figure 1 B). The assembly included an additional 150 kb of telomeric repeat. The sub-telomeric extension sequences had an unremarkable GC proportion (45% - 49%) and contained unique sequence, including predicted protein coding loci that had support from uniquely-mapping transcript evidence (Besnard et al., 2017). Notably, the telomere extensions contained repeat families that were largely limited to the extensions (Table 1; Figure 1 B). One predicted protein coding gene overlapped the internal telomere addition site (gene 632t, encoding a neprilysin M13 metallopeptidase homologue, on Otip_V right end (Otip_V R)). This gene prediction appears to represent a full-length locus, as it aligns well with *C. elegans* orthologues, and would therefore be predicted to be non-functional in the short-form chromosome. The repeat sequences unique to the sub-telomeric extensions were predicted to encode protein coding genes that had similarities to helitron transposon genes (Figure 1). The longest helitron-like predicted gene, 9690_t on Otip_IV left end (Otip_V L) (3985 amino acids) contains domain matches to Pif1-like, ATP-dependent DEAH box DNA helicases. Sequences similar to this putative helitron-like gene are present on eleven of the twelve telomere extensions (it was absent from the Otip_I L extension; Figure 1 A, Table 1). A single additional match was found on Otip_V, near the rRNA cistron repeat. Scanning of the genome for helitron-like sequences using RepeatFinder (Tarailo-Graovac and Chen, 2009) and HelitronScanner (Xiong et al., 2014) identified many additional, distinct helitron-like sequences, scattered across all the chromosomes. However, the telomere-associated helitron-like elements scored poorly in these searches, suggesting they are a new, perhaps distinct family.

**Table 1:** Telomeric extension sequences in Oscheius tipulae.

| *Chromosome* | *Arm* | *Extension span*<br>*(bp)* | *Sub-telomeric*<br>*extension start*<br>*site \** | *External*<br>*TTAGGC*<br>*telomere*<br>*repeat*<br>*addition site \** | *External*<br>*TTAGGC*<br>*telomere*<br>*repeat span*<br>*(bp)* |
| --- | --- | --- | --- | --- | --- |
| **Otip_I** | L | **16822** | 27184 | 10363 | 10362 |
|  | R | **24306** | 8885185 | 8860879 | 31405 |
| **Otip_II** | L | **10122** | 25180 | 15059 | 15058 |
|  | R | **10413** | 9010174 | 8999761 | 19801 |
| **Otip_III** | L | **4060** | 16298 | 12239 | 12238 |
|  | R | **25188** | 9621518 | 9596330 | 13932 |
| **Otip_IV** | L | **34114** | 39782 | 5669 | 5668 |
|  | R | **4711** | 11184555 | 11179844 | 463 |
| **Otip_V** | L | **23599** | 30663 | 7065 | 7064 |
|  | R | **29106** | 10812563 | 10783457 | 4916 |
| **Otip_X** | L | **32697** | 51845 | 19149 | 19148 |
|  | R | **133498** | 11309184 | 11175686 | 14155 |
| **Total** |  | **348636** |  |  | **154210** |

\* Base position with reference to the full chromosome sequence length (including telomere hexamer repeat and telomere extensions). This position is the site of addition of telomeric hexamer repeat in the short-form chromosomes.

The final nuclear genome assembly was a significant improvement over the previous reference in contiguity and completeness (Besnard et al., 2017) (Table 2). The 253 contigs of the previous assembly were super-scaffolded using genetic map information. All but eight of these previous assignments were replicated in the new assembly, and the data underlying the new assembly affirmed the eight new assignments. The longest contigs from the previous assembly tended to be found in the centres of the chromosomal contigs, and the ends of the new chromosomal contigs were represented by multiple shorter contigs in the previous assembly (Figure S5). The proportion of complete nematode BUSCO loci (nematoda_odb10) identified in both assemblies was 90%. Close analysis of the assemblies identified candidates for some of the 254 apparently missing orthologues. We note that similar low BUSCO completeness scores were recorded for the closely-related nematode *Auanema rhodensis* (Table S9).

**Table 2:**
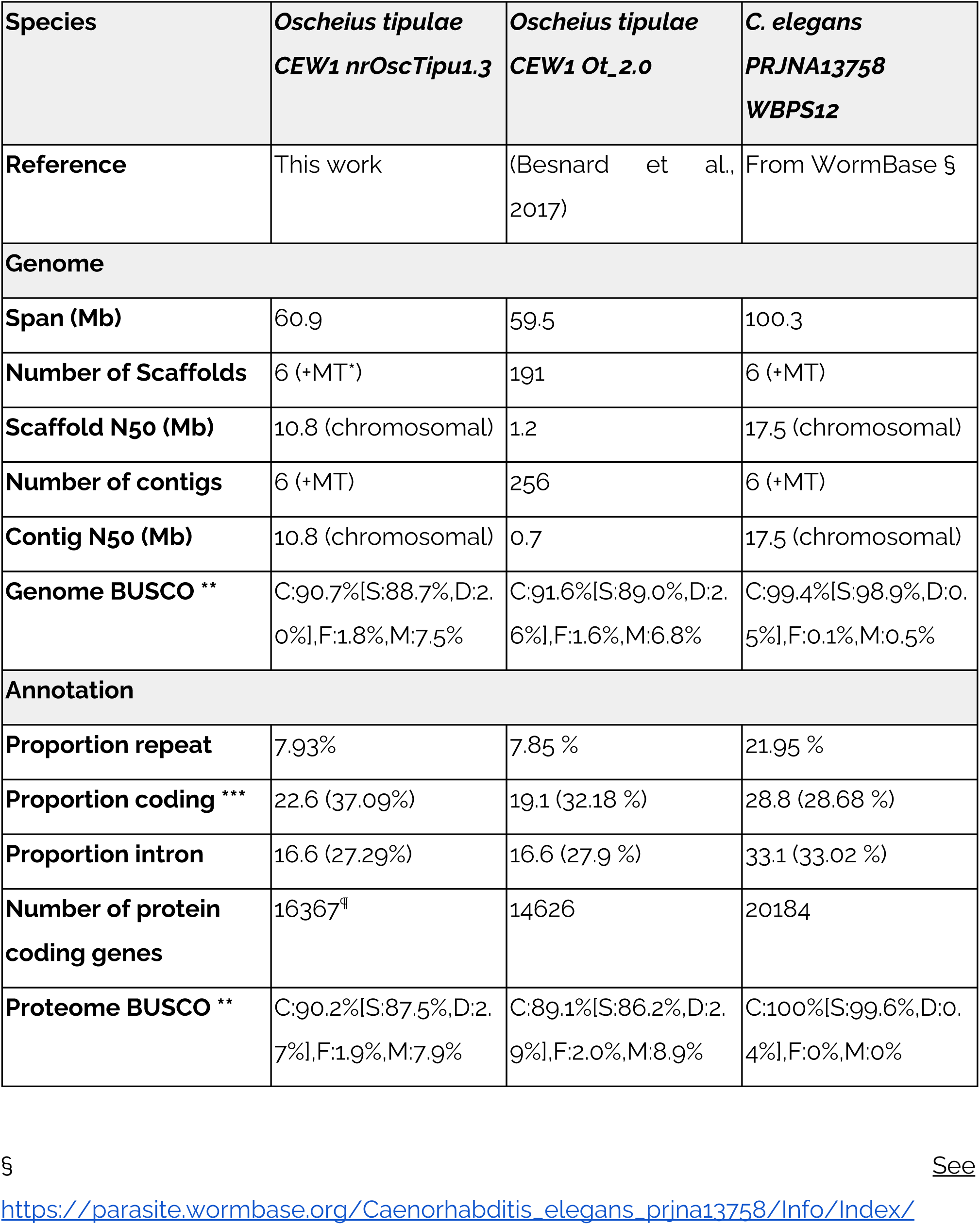
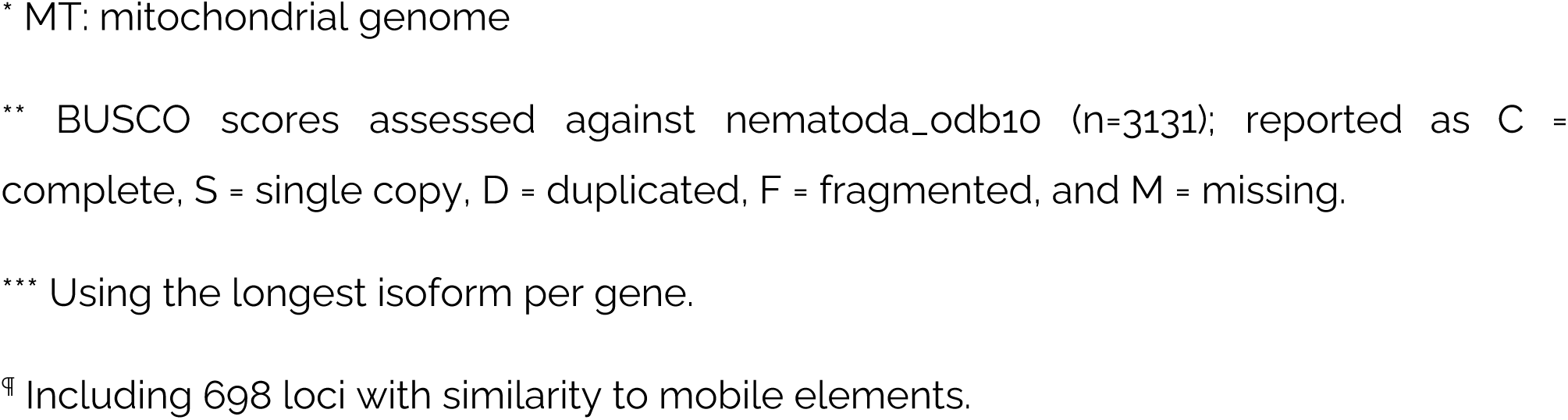
Metrics of the Oscheius tipulae CEW1 genome assembly.

| Species | *Oscheius tipulae*<br>CEW1 nrOscTipu1.3 | *Oscheius tipulae*<br>CEW1 Ot_2.0 | *C. elegans*<br>PRJNA13758<br>WBPS12 |
| --- | --- | --- | --- |
| Reference | This work | (Besnard et al., 2017) | From WormBase § |
| **Genome** |  |  |  |
| Span (Mb) | 60.9 | 59.5 | 100.3 |
| Number of Scaffolds | 6 (+MT*) | 191 | 6 (+MT) |
| Scaffold N50 (Mb) | 10.8 (chromosomal) | 1.2 | 17.5 (chromosomal) |
| Number of contigs | 6 (+MT) | 256 | 6 (+MT) |
| Contig N50 (Mb) | 10.8 (chromosomal) | 0.7 | 17.5 (chromosomal) |
| Genome BUSCO ** | C:90.7%[S:88.7%,D:2.0%],F:1.8%,M:7.5% | C:91.6%[S:89.0%,D:2.6%],F:1.6%,M:6.8% | C:99.4%[S:98.9%,D:0.5%],F:0.1%,M:0.5% |
| **Annotation** |  |  |  |
| Proportion repeat | 7.93% | 7.85 % | 21.95 % |
| Proportion coding *** | 22.6 (37.09%) | 19.1 (32.18 %) | 28.8 (28.68 %) |
| Proportion intron | 16.6 (27.29%) | 16.6 (27.9 %) | 33.1 (33.02 %) |
| Number of protein coding genes | 16367<sup>¶</sup> | 14626 | 20184 |
| Proteome BUSCO ** | C:90.2%[S:87.5%,D:2.7%],F:1.9%,M:7.9% | C:89.1%[S:86.2%,D:2.9%],F:2.0%,M:8.9% | C:100%[S:99.6%,D:0.4%],F:0%,M:0% |

§
See
https://parasite.wormbase.org/Caenorhabditis_elegans_prjna13758/Info/Index/
\* MT: mitochondrial genome
\*\* BUSCO scores assessed against nematoda\_odb10 (n=3131); reported as C = complete, S = single copy, D = duplicated, F = fragmented, and M = missing.
\*\*\* Using the longest isoform per gene.
¶ Including 698 loci with similarity to mobile elements.

### Comparing chromosome structure in *O. tipulae* and other nematodes

A striking feature of the *C. elegans* genome is the strong patterning of genic and non-genic features along each chromosome (C. elegans Sequencing Consortium, 1998). Repeats are more abundant on the autosome arms, and are largely excluded from the chromosome centres, while GC proportion has higher variance in the arms. The *C. elegans* X chromosome has a similar pattern albeit less pronounced. These patterns are likely to be generated by the local recombination rate, which is high on autosome arms and lower in the autosome centres and on the X chromosome (Rockman and Kruglyak, 2009). We explored the chromosomal *O. tipulae* genome for similar patterns. The arms of *O. tipulae* chromosomes show a higher repeat and intronic sequence fraction and a lower exonic fraction compared to the centres (Figure 2). The pattern of GC proportion along *O. tipulae* chromosomes also matched expectations from *C. elegans* (and other species, Figure S6), except for two regions. Both Otip_IV and Otip_V have regions where there is a step change in GC proportion (Figure 1 A, inner circle). The boundaries of these step changes were examined and were supported by a normal (320 fold) coverage of PromethION reads. We interpret these as recent inversions where background processes have not yet restored the local GC proportion pattern.

**Figure 2:**
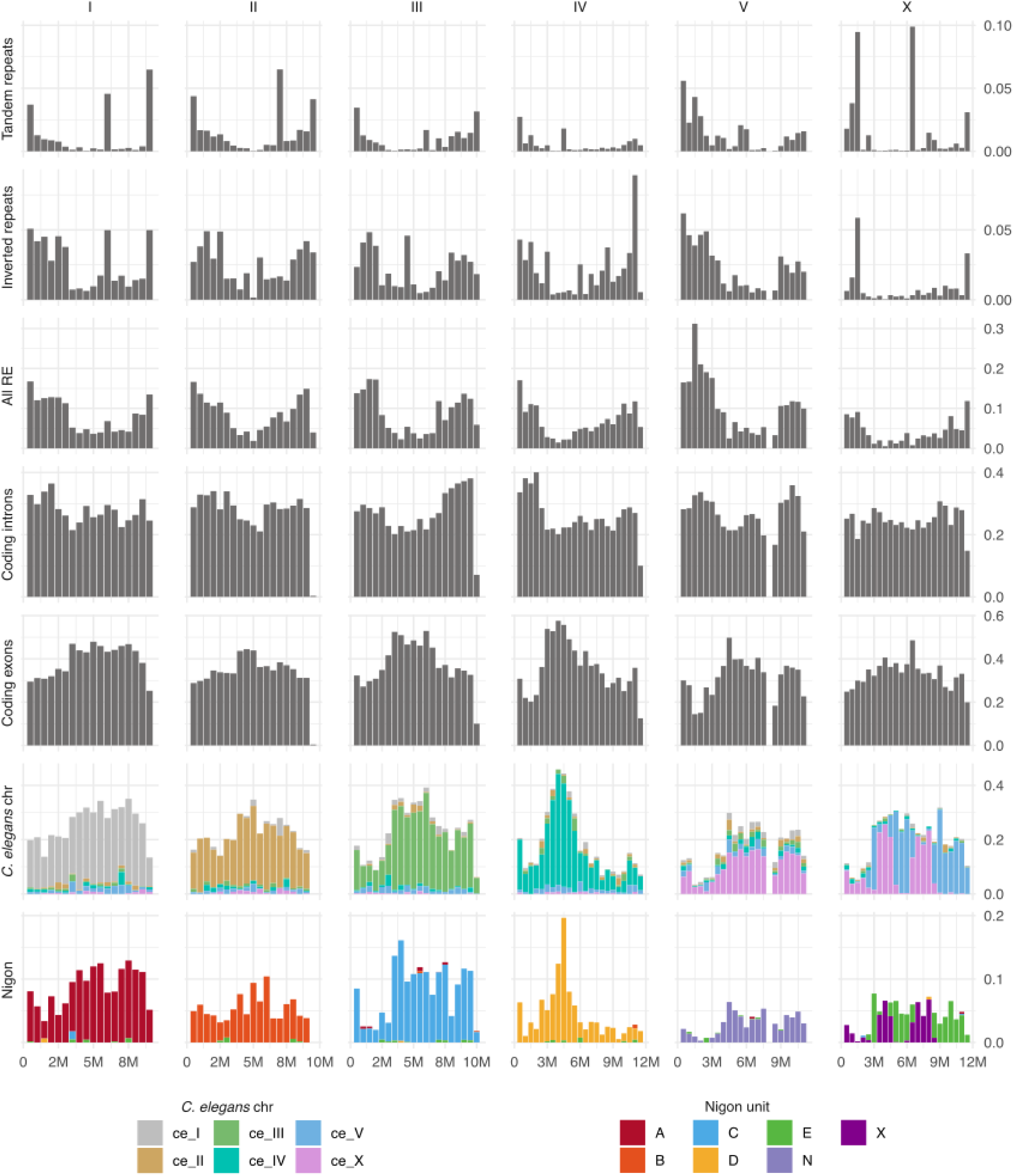
The *Oscheius tipulae* CEW1 genome. Repeat and gene densities along *O. tipulae* chromosomes, and mapping of placement of orthologues of *C. elegans* and Nigon element assigned loci. Feature densities (repeat and gene features and orthologue mapping) were calculated in 500 kb non-overlapping windows along each chromosome. For the mapping of orthologues (rightmost two chart sets), the plots are stacked bars coloured by either *C. elegans* chromosome (Ce chr) or Nigon element group (Nigon).

As expected from comparisons within the genus *Caenorhabditis* (Slos et al., 2017) (Stevens et al., 2019) (Stevens et al., 2020) and between *Caenorhabditis* and the strongylomorph *H. contortus (Doyle et al., 2020)*, while neighbouring orthologous genes tended to be found on the same linkage groups in *O*. *tipulae* and in *C. elegans*, local gene neighbourhoods were not highly conserved (Figure 3). Interestingly, genes in the centres of the *C. elegans* autosomes were not more likely to be retained in the centres of *O. tipulae* chromosomes, and gene neighbourhood conservation was largely absent apart from conserved operonic gene sets.

**Figure 3:**
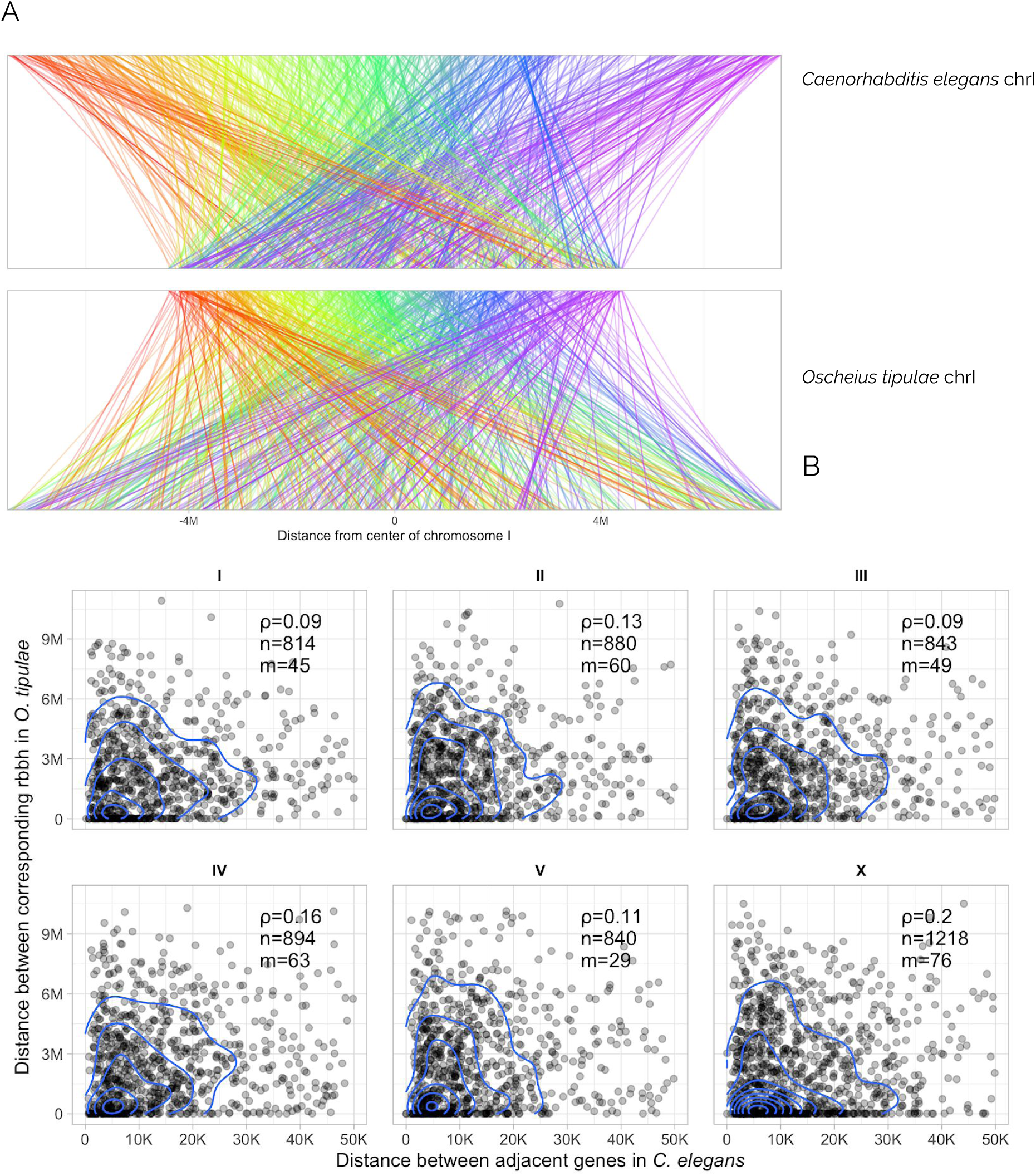
Local gene order comparison between *Oscheius tipulae* and *Caenorhabditis elegans*. A Whole chromosome comparison of *Oscheius tipulae* chromosome I (Otip_I) and its homologue, *Caenorhabditis elegans* chromosome I (Cele_I). Lines link loci that are reciprocal best BLAST hits. In the top panel the lines are coloured by their placement on *C. elegans* Cele_I (a spectrum from red on the left arm through to purple on the right arm). In the lower panel the lines are coloured by their placement on *O. tipulae* Otip_I (a spectrum from red on the left arm through to purple on the right arm). B Gene neighbourhoods are not conserved between *O. tipulae* and *C, elegans*. We measured the separation distance between each pair of neighbouring *C. elegans* loci for which we could identify single-copy reciprocal best-BLAST relationships between *O. tipulae* and *C. elegans*, and plotted the separation between these orthologue pairs in *C. elegans* (x axis) and *O. tipulae* (y axis) for each chromosome. Ortholog pairs more than 50 kb apart in *C. elegans* were excluded. R: Spearman correlation; n: number of orthologous pairs including pairs excluded from the plot; m: the number of pairs excluded from each plot.

### Chromosomal elements are conserved across the Rhabditida

Previously we defined conserved nematode chromosomal elements, called Nigon elements, through manual comparison of five genomes from species in Rhabditina (Clade V) (Tandonnet et al., 2019). Manual generation of chromosome assignments is not sustainable, and piecewise addition will fossilise initial taxonomic and data biases. We therefore developed an objective, algorithmic method to identify and group loci that define conserved elements based on shared chromosomal co-location (Figure 4 A). Using this method we were able to include all available rhabditid nematode genomes that have been reported to be chromosomally-complete or near-complete. We identified 2191 loci that had a one-to-one orthologous relationship between most species. These formed seven clusters of loci co-located on chromosomes in most species. These clusters of loci were used to paint the nematode chromosomal assemblies, and replicate our previous, manual definition of Nigon elements. Then clusters had between 534 loci (defining Nigon element A) and 119 (defining NigonX) (Figure 4 B and Table S10). The *Meloidogyne hapla* genome had low BUSCO scores and low representation of orthologues in all the Nigon element sets.

**Figure 4:**
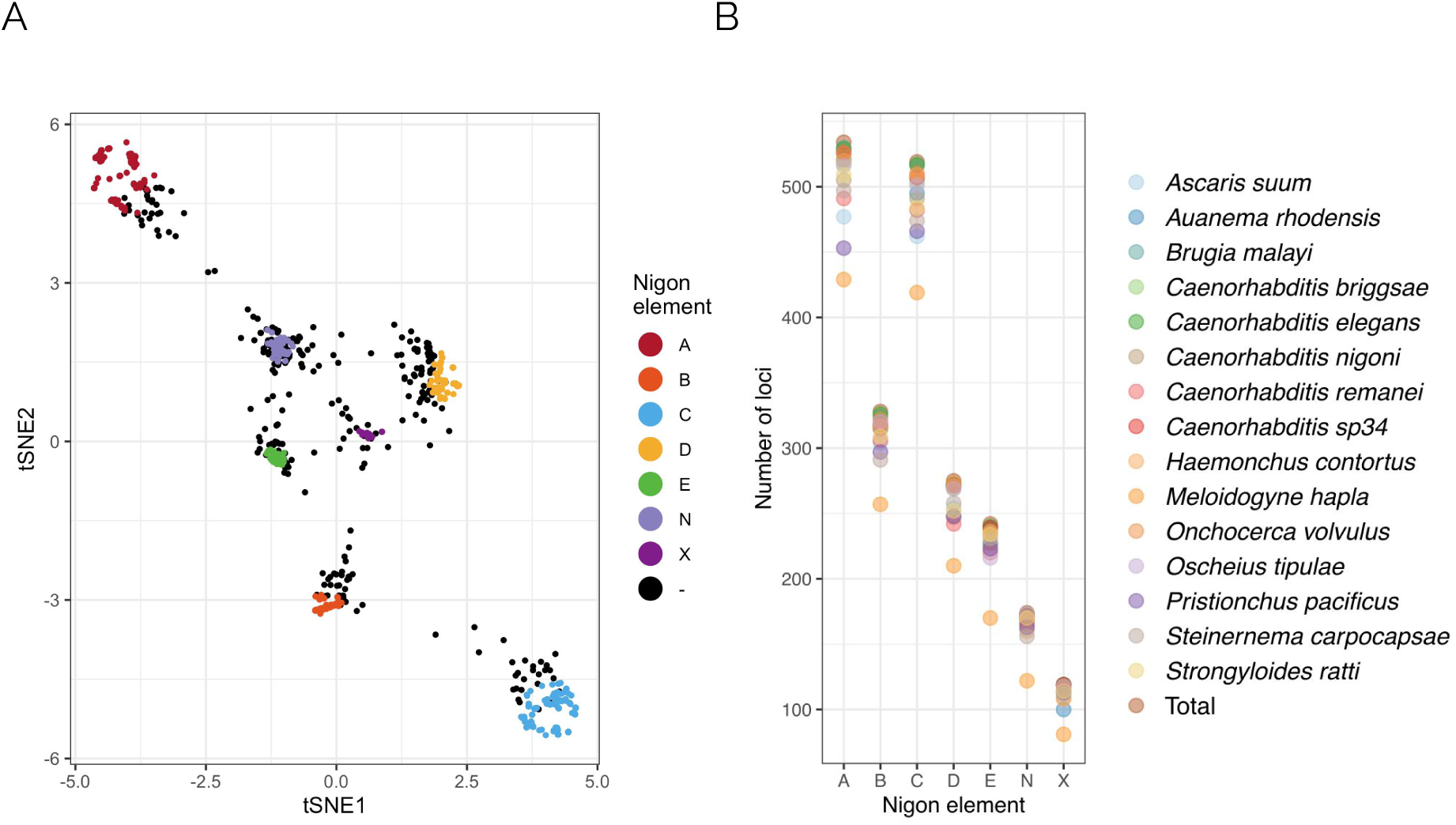
Loci that define Nigon elements in rhabditid nematodes. A t-distributed Stochastic Neighbor Embedding (t-SNE) plot of the Gower distance between 3412 orthologous gene families mapped to the chromosomal assemblies of 9 species. The 2191 loci that are included in the Nigon element sets are coloured. The black dots represent loci not assigned to a Nigon unit. B For each Nigon-defining set of loci, we counted the number found in each species. The total number of loci per set is indicated by a star. The assembly of *Meloidogyne hapla* has the fewest loci and lowest proportion of loci in all Nigon sets. C The proportion of Nigon-defining loci found in each species’ genome correlates with the BUSCO completeness of the genome.

### Independent chromosomes in iB. Malayi *and* O. volvulus*, and a set of* Chromosome evolution and homology in the Rhabditida

The general conservation of chromosome number (n=6) in rhabditid nematodes might suggest that these karyotypes reflect a static pattern of locus co-location and Nigon element structure. Previous analyses suggest that this is not the case (Rödelsperger et al., 2017; Tandonnet et al., 2019), and we went on to use our chromosome painting to define Nigon element structure in each species (Figure 2, Figure 5) and to build a model of rhabditid karyotype evolution (Figure 6). We recapitulated previous findings made on a limited set of species (Tandonnet et al., 2019) and extended the Nigon element schema to species across Rhabditida. In general, Nigon elements were found as independent chromosomes across Rhabditida, but many species showed patterns of Nigon element defining locus mixing that evidenced past fusions and breakages.

**Figure 5:**
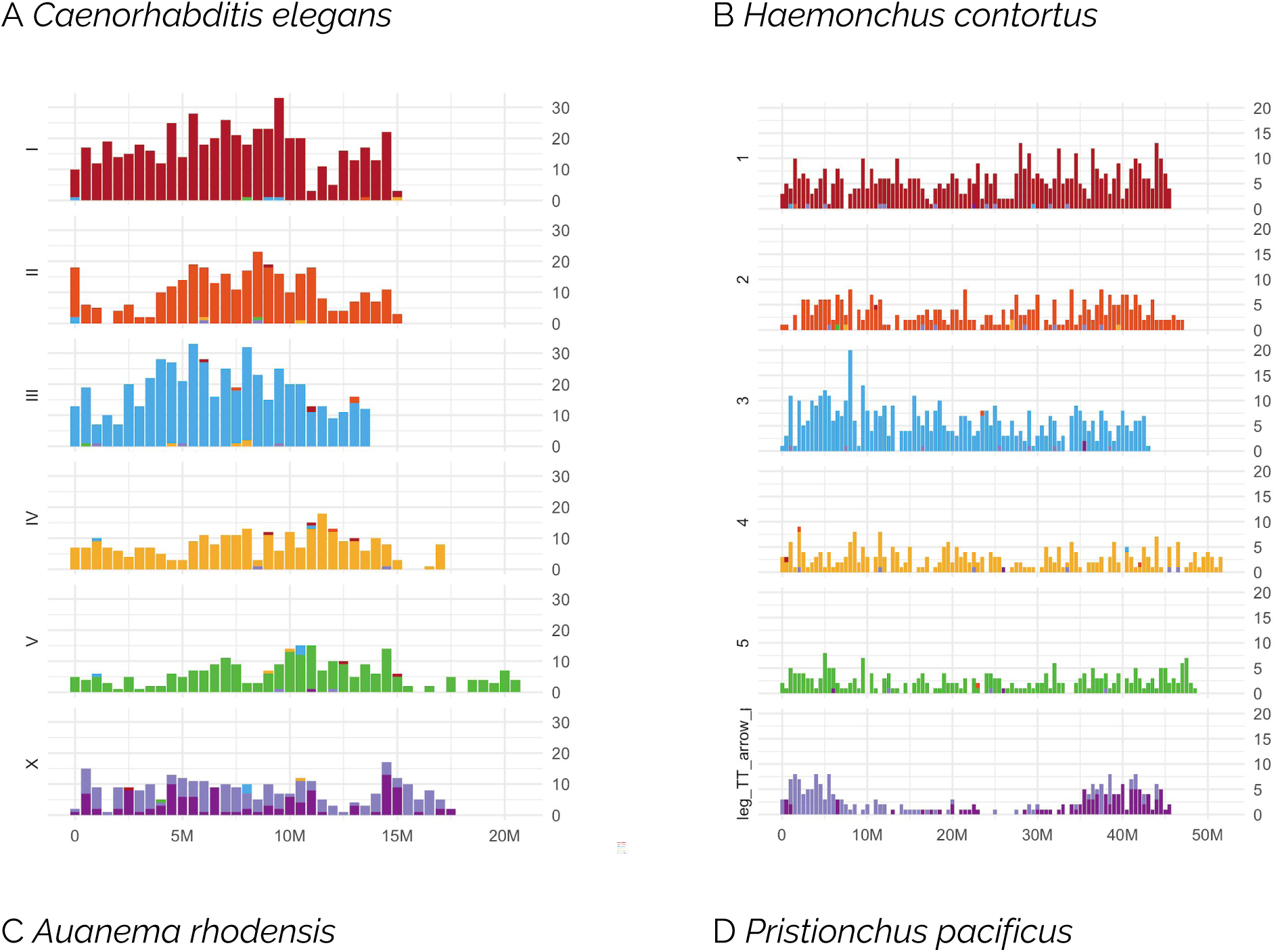
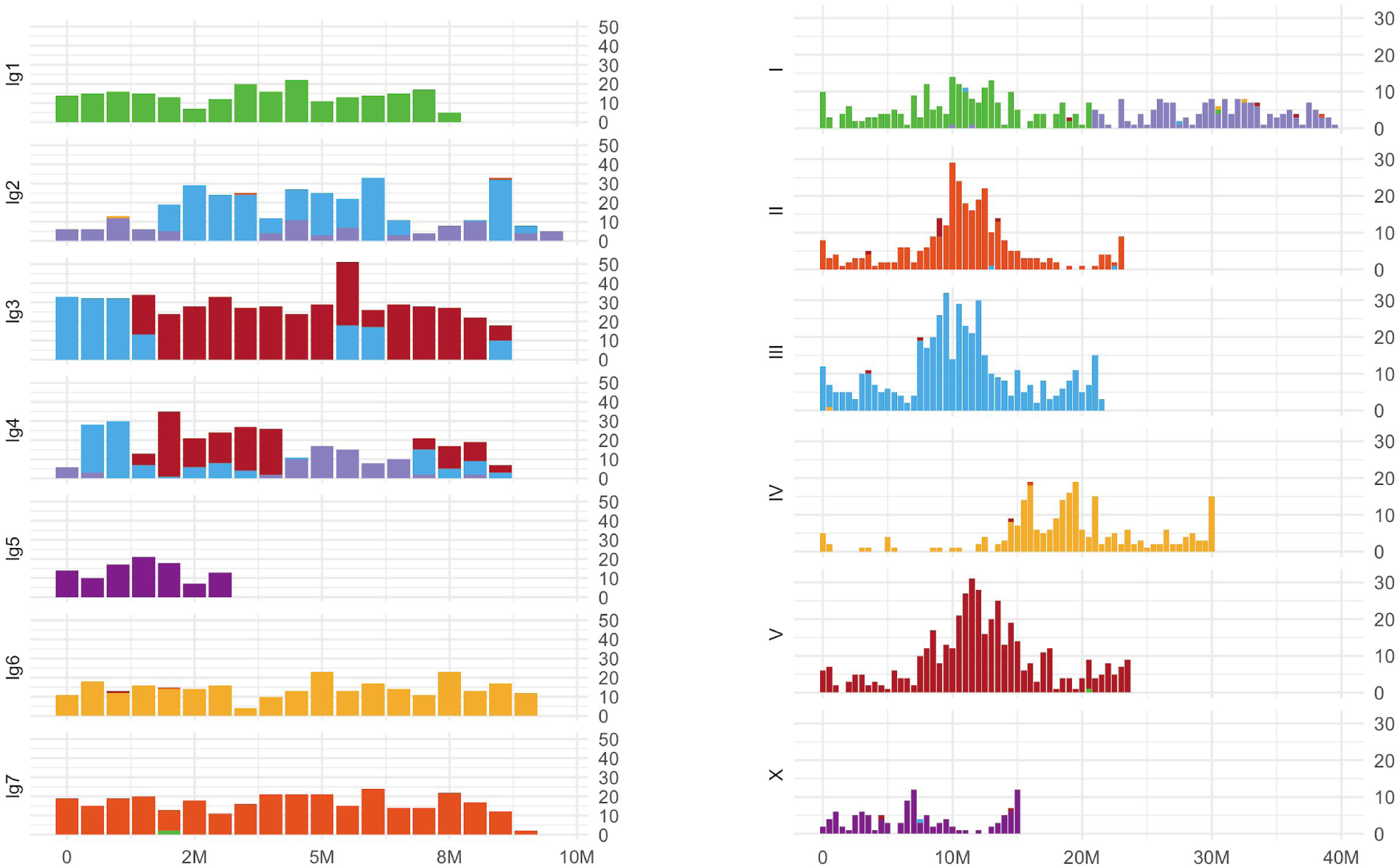
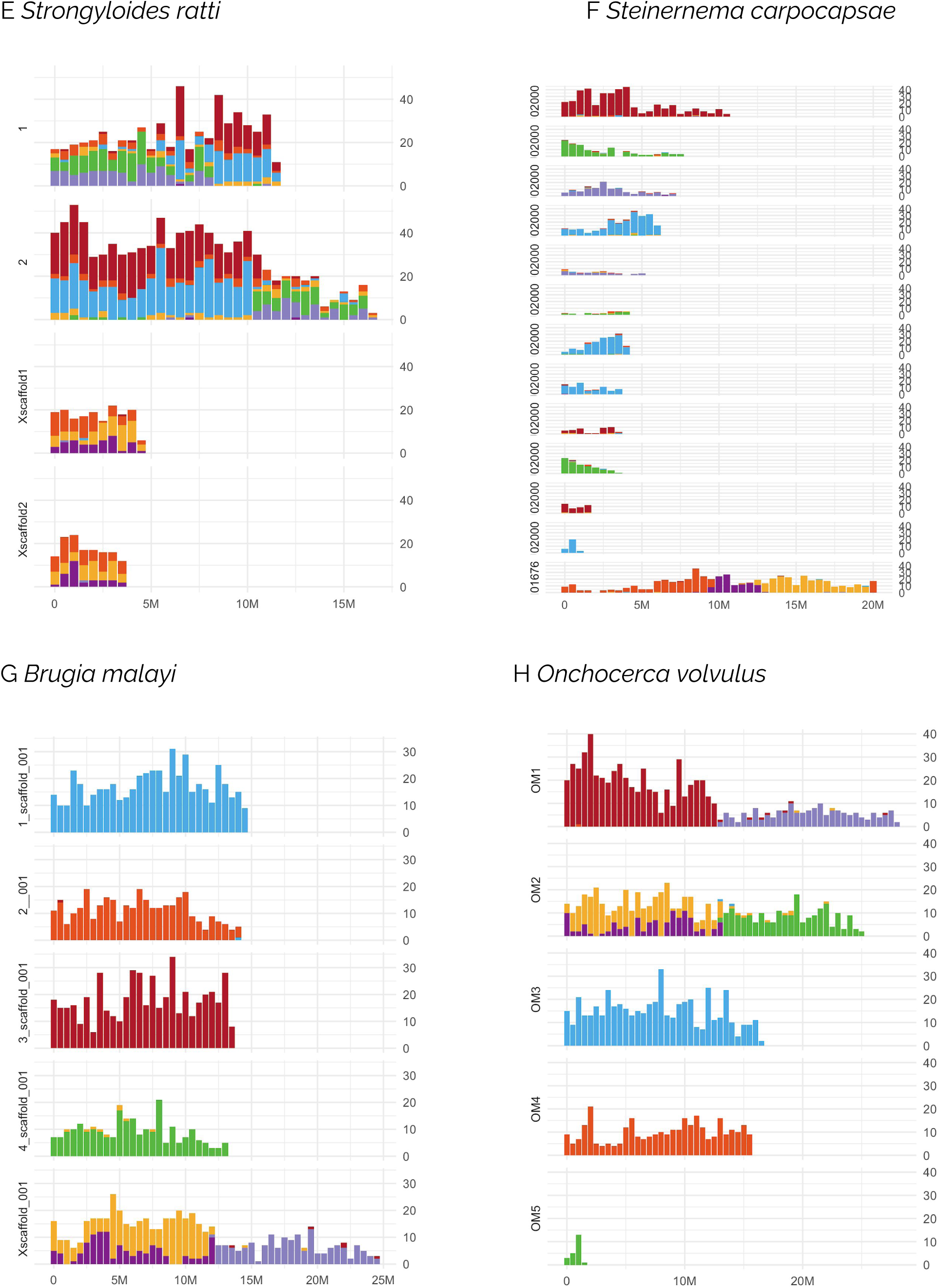
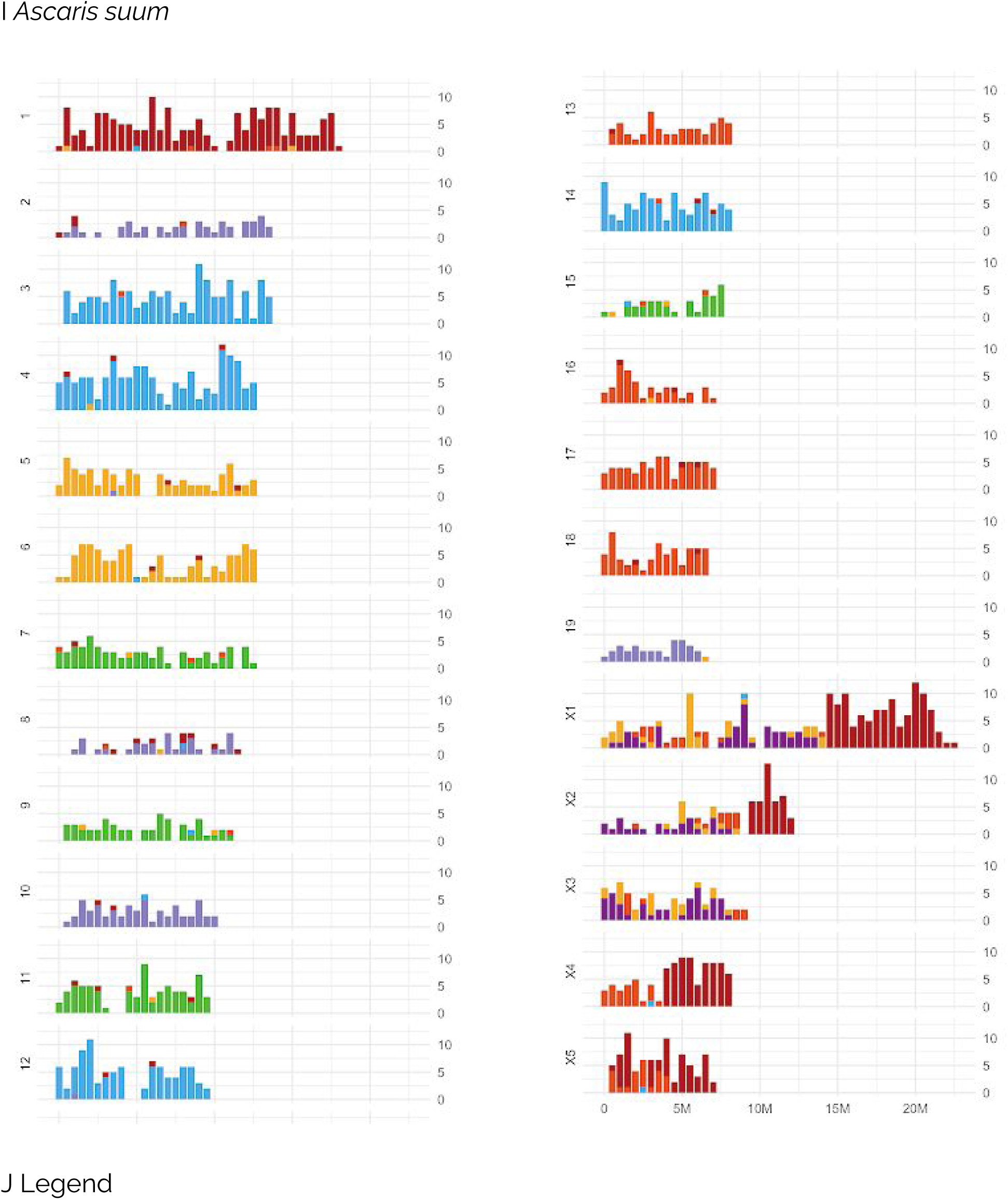
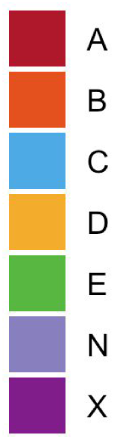
Nigon painting of Rhabditid nematode chromosomes. Examples of nematode chromosomal assemblies painted by their content of Nigon-defining loci. Each subgraph shows the count of loci mapped in non-overlapping 0.5 Mb windows along a chromosome as a stacked histogram coloured by Nigon origin. The X- and Y-axes are scaled to the maxima within a species in each panel (X: chromosome length, Y: Nigon-defining loci per interval) within each species. The legend in panel J applies to all nine chromosome panels. A *Caenorhabditis elegans* (Rhabditina, Rhabditomorpha), B *Haemonchus contortus* (Rhabditina, Rhabditomorpha), C *Auanema rhodensis* (Rhabditina, Rhabditomorpha), D *Pristionchus pacificus* (Rhabditina, Diplogasteromorpha), E *Strongyloides ratti* (Tylenchina, Panagrolaimomorpha), F *Steinernema carpocapsae* (not a fully chromosomal assembly; Tylenchina, Panagrolaimomorpha), G *Brugia malayi* (Spirurina, Spiruromorpha), H *Onchocerca volvulus* (Spirurina, Spiruromorpha), I *Ascaris suum* (Spirurina, Ascaridomorpha). Panel J shows the colour key for the other panels.

**Figure 6:**
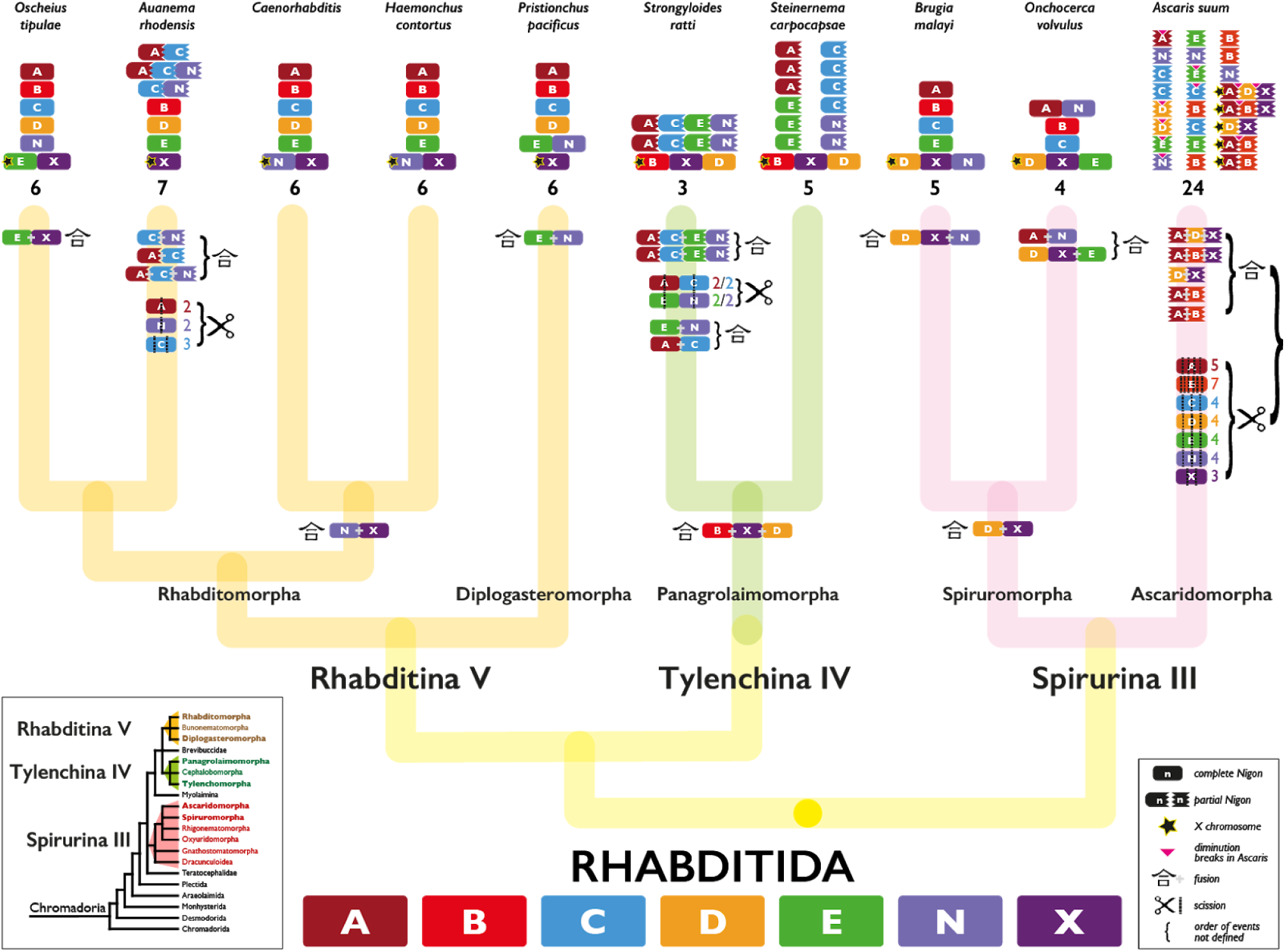
A model of chromosome evolution in Rhabditida. For each species the classification of chromosomes to Nigon elements is shown (see Figure 2, Figure 5 and Supplemental Figure S7), including rearranged chromosomes. X chromosomes are indicated by a star. For each lineage we have inferred the patterns of chromosome scission (the ✂ symbol; the number indicates the number of fragments resulting) and fusion (the Kanji symbol for “fusion point” 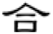) on the tree. Where the order in time of events is not resolved, we have bracketed them. In *A. suum* a pink triangle indicates positions of internal cleavage of germline chromosomes during diminution. Note that the “partial Nigons” in *S. carpocapsae* are assembly scaffolds rather than chromosomes. The cladogram representing the phylogeny of Chromadoria nematodes (inset, lower left; (Blaxter and Koutsovoulos, 2015)) is derived from phylogenetic analysis of shared protein coding genes. In the main figure we have resolved a previous uncertain polytomy in Rhabditomorpha to favour a closer relationship between *Haemonchus contortus* and the genus *Caenorhabditis* based on the exact conservation of linkage groups.

In Rhabditina, all seven of the Nigon elements were present as distinct chromosomes in at least one species. A relatively limited number of fusions and scissions were necessary to explain the observed patterns of Nigon assignment. For example, in *P. pacificus* Ppa-chrI was identified as a recent fusion of NigonA and NigonN (Figure 5 D). As noted previously (Rödelsperger et al., 2017; Tandonnet et al., 2019), Ppa-chrI retains structural signal of two chromosomal elements, with a dual pattern of repeat density and gene density peaks, and these features are consistent with the NigonA and NigonN portions of this chromosome. The *P. pacificus* X chromosome contained only loci from NigonX. Nigon painting of the *Caenorhabditis* species and *H. contortus* was very similar, and showed that the n=6 karyotype of these species is derived from the seven Nigon elements through fusion of NigonN with NigonX to form the X chromosome (Table 3). In contrast to the NigonA-NigonN fusion in *P. pacificus*, the NigonN and NigonX loci in the X chromosomes of *Caenorhabditis* and *H. contortus* are intermixed, suggesting that the fusion was ancestral to the split between these nematodes, and that processes of intrachromosomal rearrangement have removed evidence of distinct NigonN or NigonX domains. This NigonX-NigonN fusion was not observed in other Rhabditina species. In *O. tipulae* NigonX was found to have fused with NigonE to for the X chromosome, and the blocky pattern of locus distribution suggested that the fusion was more recent than the NigonN-NigonX fusion in *Caenorhabditis* and *H. contortus* (Figure 5 A and B). In *A. rhodensis* the loci defining NigonX were all found on Arh-lg5, which is the X chromosome (Figure 5 C). NigonA, NigonC and NigonN loci were distributed, with a blocky pattern, across three *A. rhodensis* autosomes (Arh-chr2, Arh-chr3, Arh-chr4), suggesting a relatively recent set of scission and fusion events (Figure 5 C). Thus, at the base of Rhabditina, we predict there were seven distinct linkage groups, corresponding to Nigon elements A through E, N and X (Figure 6).

**Table 3:**
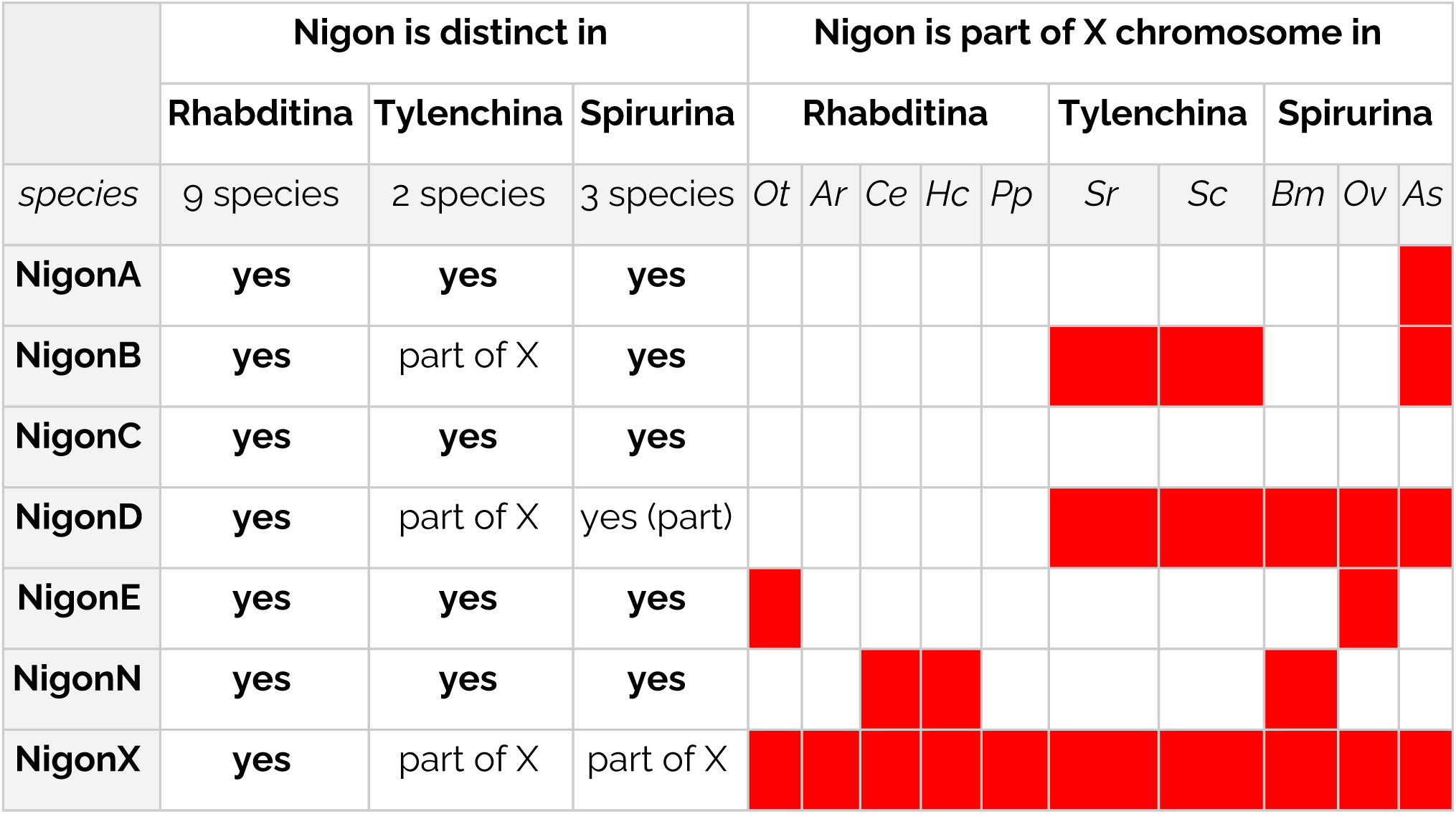
Nigon elements and the X chromosomes of rhabditid nematodes.

In Spirurina, NigonA was an independent autosome in *B. malayi* (*Bma-chr3*) and a NigonA element was identified as recently fused with a NigonN element in *O. volvulus* to form *Ovo-OM1* (Figure 5 G and H; Supplemental Figure S7). Chromosomes consisting nearly completely of NigonA loci were found in *A. suum* (*Asu-chr1*) (Figure 5 I). NigonB is an independent autosome in both *B. malayi* and *O. volvulus*, and four *A. suum* autosomes are painted only by NigonB loci (Figure 5 G, H and I). In *A. suum*, some NigonB loci are also found on the X chromosomes. Similarly, NigonC loci paint independent chromosomes in *B. malayi* and *O. volvulus* and four autosomes in *A. suum.* NigonN appeared to have fused recently and independently in *B. malayi* (with NigonD to form the X chromosome; Figure 5 G) and *O. volvulus* (with NigonA to form *Ovo-OM1*; Figure 5 H). That these fusions are recent is supported by patterns of repeat and GC content across the chromosomes. In *A. suum* NigonN loci are largely restricted to four NigonN autosomes (Figure 5 I, Figure 6). The X chromosomes of *A. suum*, *B. malayi* and *O. volvulus* each carried evidence of a fusion between NigonD and NigonX. It was not easy to discern whether this fusion was ancestral to Spirurina or arose independently in the Ascarididomorpha and Spiruromorpha (Figure 6). Independent fusion was supported by the finding that there are distinct *A. suum* autosomes only painted by NigonD, that NigonX loci while wholly limited to the five *A. suum* X chromosomes are not always associated with NigonD loci, and that NigonD and NigonX loci form distinct blocks in the *A. suum* X chromosomes. On *A. suum* Asu_chrX1, the NigonD and NigonX domains are resolved as distinct somatic chromosomes by chromatin diminution. In *B. malayi* and *O. volvulus* the NigonD and NigonX loci were intermixed, suggesting long-term association and mixing by intrachromosomal rearrangement.

It is notable that while all the autosomes of *A. suum* were painted by loci from a single Nigon element set, all five X chromosomes had mixed origins, involving blocks of NigonA, NigonB, NigonD and NigonX loci. The chromosomes of *A. suum* are subject to chromatin diminution in somatic cells of the embryo (Wang et al., 2020). This process generates remodelled telomeres for all chromosomes, and specific cleavage at internal sites in some chromosomes such that somatic cells have more chromosomes (but less genetic material overall) than do germline cells. It is striking that all but one of the intrachromosomal cleavage points were between blocks of chromosome with distinct Nigon identity. The one cleavage not between Nigon blocks (in *Asu-chr1*) separated two NigonA components. Not all blocks of Nigon identity were separated by cleavage during diminution (Figure 6).

In Tylenchina, retention of ancestral units, breakages and fusions also describe the present day chromosome structures observed (Figure 6). In *S. carpocapsae*, which is not fully chromosomally assembled, the twelve autosomal scaffolds, which correspond to four autosomes, each had a single Nigon identity. We predict that these scaffolds will assemble to yield four autosomes corresponding to NigonA, NigonC, NigonE and NigonN. The *S. carpocapsae* X chromosome comprised three domains corresponding to NigonB, NigonX and NigonD (Figure 5 F). *S. ratti* has two autosomes and an X chromosome. The autosomes were modeled as being a complex product of a two stage fusions-scissions-fusions process. The first fusions were between NigonA and NigonC, and between NigonE and NigonN. After some time (as evidenced by the intermixing of loci) these fusion chromosomes were split, and refused to form Sr-chr1 and Sr-chr2. The second set of scissions and fusions was relatively recent, as each autosome had distinct domains corresponding to the presumed ancestral fusions. The *S. ratti* X chromosome was painted by loci corresponding to NigonB, NigonD and NigonX (Figure 5 E). The NigonB, NigonN and NigonX loci were fully intermixed. Again it was difficult to disentangle ancient fusion versus parallel fusion of NigonD and NigonX in Tylenchina. The distinctness of the NigonX and NigonD domains in the *S. carpocapsae* argues for independent events. Nonetheless it is striking that NigonD and NigonX are associated with the X in both Tylenchina and Spirurina. *Meloidogyne hapla* has seventeen chromosomes, but Nigon painting of these did not yield definitive Nigon assignments (Supplementary Text 2). We noted that there was no association between NgonD and NigonX loci on the *M. hapla* chromosomes.

The independent existence in a common ancestor of Rhabditina, Tylenchina and Spirurina of elements corresponding to Nigons A, B, C, E and N was evident from the identification of chromosomes, or distinct chromosome domains, in all three suborders corresponding to these elements. NigonD and NigonX were not associated in Rhabditina, but were variably associated in Tylenchina and Spirurina. It was unclear whether this should be modeled as an ancient fusion with differential resolution in extant species, or repeated fusion events. The existence of chromosomal domains corresponding to NigonD and NigonX in *S. carpocapsae* (Tylenchina) and *A. suum* (Spirurina) and of NigonD-only autosomes in *A. suum* argued against a single, ancestral event (Figure 6).

### Dynamic evolution of rhabditid sex chromosomes

NigonX was the only element consistently associated with the sex chromosome all the nematode species analysed (Table 3). Additionally the NigonX element was much more likely to be involved in fusions with other Nigon elementsHowever, the histories of loci defining NigonD and NigonX were intertwined in Spirurina and Tylenchina. In Tylenchina, the X chromosomes were ancestral fusions of NigonB, NigonD and NigonX, and this fusion was fully intermixed in *S. ratti* (Figure 5 E) but unmixed in *S. carpocapsae* (Figure 5 F). NigonX loci were found on three of the five *A. suum* X chromosomes (As-chrX1, As-chrX2 and As-chrX3), mixed or fused with other Nigon element loci (Figure 5 I). The other two *A. suum* X chromosomes (As-chrX4 and As-chrX5) are fusions of NigonA and NigonB loci. The NigonA loci tended to form distinct domains in all the *A.suum* X chromosomes, which suggests relatively recent fusion, but the NigonD and NigonX loci were intermixed. *A. suum* also had two autosomes that contained only NigonD loci (As-chr5 and As-chr6), suggesting that this element had an independent existence and fissioned into at least two parts before fusion with NigonX. In both spiruromorph nematodes, a NigonD plus NigonX intermixed domain was found in the X chromosome, but this had fused with different, autonomous Nigon elements in *B. malayi* (with NigonN) and *O. volvulus* (with NigonE) (Figure 5 G, H).

The complete admixture of NigonD and NigonX loci in these spiruromorph X chromosome domains suggests ancient fusion, especially since the *B. malayi* and *O. volvulus* genomes are largely collinear both within non-fused chromosomes and within fused chromosome blocks (Foster et al., 2020). Because NigonX and NigonD are present, intermixed, in the X chromosomes of both Ascaridomorpha and Spiruromorpha, the NigonD-NigonX fusion could be ancestral to Spirurina. However the presence of NigonD-only autosomes in *A. suum* suggested that NigonD was present as an independent element in this lineage, and mapping of the relationships between NigonD loci in *A. suum* and spiruromorph genomes does not show any particular grouping. Thus we modeled these NigonD-NigonX fusions as independent events (Figure 6).

NigonX and NigonD were also present in the X chromosomes of Tylenchina, as part of a B-X-D fusion. In *S. carpocapsae*, this fusion appeared to be recent, as the three sets of Nigon element-derived loci occupied distinct domains, with NigonX central (Figure 5 F). In *S. ratti* the loci from NigonB, NigonD and NigonX were intermixed on *Sra-chrX*, but some NigonD loci were found on the two autosomes (Figure 5 E). These autosomal NigonD loci were found in association with some NigonB loci, perhaps as a result of translocation from an ancestral intermixed NigonB-NigonD chromosome. The distinctness of the NigonD domain within *S. carpocapsae* Sc_X suggested that NigonD was an independent element in Tylenchina also. The B-D-X fusion is, we suggest, an association of NigonD and NigonX independent of that in Spirurina.

### Functional coherence of Nigon element loci

The distinct histories of the gene sets associated with each Nigon element means that these sets of loci have been linked for a significant period of time, and may have been selected to stay together or evolved to collaborate. We explored whether such association might reflect the shared biological function of these genes. We interrogated functional enrichment of the Nigon-defining loci through the KEGG pathway annotation of *C. elegans* orthologues. We first compared KEGG pathway annotations of Nigon-defining loci to those of the full gene set of *C. elegans* (Table S11). Terms relevant to RNA transport were enriched in NigonA loci and NigonC loci, ribosome biogenesis was enriched in NigonA loci and spliceosome pathway was enriched in NigonC loci. ErbB signaling and calcium signaling pathways were enriched in the Nigon D locus set. In NigonN loci Hippo signaling was enriched. In NigonX loci, annotations relevant to axon regeneration, calcium signaling pathway and neuroactive ligand-receptor interaction were significantly enriched. No KEGG pathways were enriched among loci of NigonB or NigonE. The Nigon-defining loci were drawn from a specific, conserved subset of all *C. elegans* loci, and this conservation will have *a priori* biased the annotations being assessed. A also compared the annotations associated with each Nigon locus set to those of the set of all 2175 Nigon loci and only detected enrichment in NigonX loci, in the KEGG axon regeneration pathway (enrichment significance 4.40 e10-4).

## DISCUSSION

### New technologies and complete genome sequencing of *O. tipulae*

New sequencing technologies and the development of improved assembly toolkits are generating more highly-contiguous reference genome assemblies. Here we use the Oxford Nanopore long reads to generate chromosomally-complete contigs representing all the nuclear chromosomes and the mitochondrion of the free-living nematode *Oscheius tipulae*. While the data are sufficient to generate a chromosomally contiguous assembly, not all assembly tools were able to generate this from the data, and there is evidently still development work to be done. Alternate methods of generating single molecule, long read data such as the Pacific Biosciences SEQUEL II CLR (single pass) and HiFi (circular consensus, multiple pass) have similar properties to PromethION data, with the HiFi standing out as having higher per-base accuracy. This higher per-base accuracy likely simplifies the assembly process, in particular in the resolution of repeat structures that are close to but not 100% identical. However the PromethION data, which can include very long reads, may be better at traversing recent segmental duplications and homogenised multicopy loci (Nurk et al., 2020). In our assembly of *O. tipulae* the only identified remaining collapsed repeat was the 6.8 kb repeat of the ribosomal RNA cistron (nSSU, 5.8S and nLSU loci), which we estimate is repeated about 117 times, summing to 801 kb. This repeat is homogenised and thus only very long range technologies, such as ultralong nanopore reads or BioNano mapping, could resolve it fully.

One unexpected and striking feature of the *O. tipulae* genome is the structure of the telomeres. The PromethION data robustly predicts two telomere repeat addition sites at each end of each chromosome, generating a core, high coverage chromosome with lower coverage subtelomeric extensions at each end. The extensions are supported by unique mapping of independently generated short Illumina data. Nearly 350 kb (or 0.6% of the genome) is in these extensions, which carry additional, expressed protein coding genes. We currently interpret the extensions as segments of chromosomes that have been specifically removed from the genomes of a proportion of cells in each nematode, possibly through developmentally regulated chromatin diminution. The presence of helitron mobile elements specifically in the subtelomeric elements is intriguing. The helitrons contain nuclease and Pif1 DEAH-box helicase domains. In yeast and other taxa, Pif1 helicases are intimately involved in DNA metabolism, and in particular in DNA replication and telomerase function and regulation. It is possible that the *O. tipulae* telomeric helitrons are parasitic elements that have generated extended subtelomeric regions in which they reside, and that they also control the excision of these telomeric extensions and the specific addition of neo-telomeres. Alternatively the nematode may have co-opted the helitrons to regulate a chromatin diminution process that regulates expression of germline-restricted genes by eliminating them from the soma, as in *A. suum* (Wang et al., 2012). In nematodes, chromatin diminution distinguishing soma from germline has been described in Ascaridomorpha and in XX and X0 sperm made by parthenogenetic female *S. papillosus (Wang and Davis, 2014)*. Given that the diminution signal we observe is present on all chromosome ends, we currently favour an ascaridomorph-like process that distinguishes a germ-line genome from a somatic one. It must be distinct from the ascaridomorph process, as the internal breakage and addition sites in *A. suum* are associated with multi-kilobase regions while the *O. tipulae* sites are precise. It may be that similar processes are present in other nematodes, and other species, but have been overlooked because of the lack of contiguity of the previously available short-read sequence data.

### Evolution of rhabditid nematode karyotypes

We have refined an approach to defining loci that define conserved linkage groups. In many taxa, it is possible to use gene neighbourhoods (gene order and synteny) to drive inference of ancestral karyotypic organisation (Kim et al., 2017). However we and others have noted that gene order is poorly conserved in rhabditid nematodes (Stein et al., 2003; Teterina et al., 2020). This is interesting, not only because the position of protein coding genes in chromosomes of *C. elegans* correlates with gene conservation, such that deeply evolutionary conserved genes tend to be found in chromosomal centres, and novel loci on the arms (C. elegans Sequencing Consortium, 1998). Despite this we observed other chromosomal features that were similar to *C. elegans*, such as the differential abundance of repeats on the presumed arms of *O. tipulae* autosomes. This suggests that distinct evolutionary processes may drive these patterns.

We were able to derive sets of loci that traveled together on linkage groups through rhabditid genome evolution by clustering orthologues based on a numerical representation of their chromosomal location in each species. This process is robust, and extendable to incorporate additional genomes. It is also applicable to other taxa where chromosomally-complete genomes are available. In Rhabditida, we identified seven clusters of loci that define seven chromosomal units, named Nigon elements. These elements are fully congruent with a previous manual estimate (Tandonnet et al., 2019). Painting the chromosomal genome assemblies of fourteen rhabditid species revealed that in no species were all of these elements present as distinct chromosomes, but each Nigon element was found as a distinct element in several species. In Tylenchina and Spirurina, we found that NigonX was fused with other Nigon elements to form the X chromosome. These fusions include NigonD and NigonA in Tylenchina and NigonD in Spirurina. We have modeled NigonD as a distinct unit despite this frequent association with NigonX because some NigonD loci uniquely painted two chromosomes in *A. suum* and the NigonD painting of the X chromosome in *S. carpocapsae* identified a single contiguous block. We were not able to apply the Nigon element model to *M. hapla*, where mapping to the 17 chromosomal scaffolds yielded only a few with majority assignment to one element. It will be informative to explore chromosomal evolution in the plant parasitic Heteroderidae further.

*Brugia* and *Onchocerca* are unusual in Spiruromorpha in having an apparent XX:XY sex determination system (Post, 2005). Within the filarial nematodes an XY system has evolved twice from an ancestral XX:X0 system, once in the ancestor of *Onchocerca* and *Dirofilaria* species and once in the ancestor of *Wuchereria* and *Brugia* species (Figure 7) (Post, 2005). It was proposed from karyotypic analyses that the neo-X chromosome in *Onchocerca* and *Brugia* arose from the fusion of an autosome with the ancestral X, and that the neo-Y chromosome in these species was just this autosomal chromosomal component (Post, 2005). Our analysis of Nigon element conservation supports this model, but additionally suggests that the enlarged X chromosomes in the two species are the results of two distinct fusions with an ancestral X: in *B. malayi* with NigonN and in *O. volvulus* with NigonE.

**Figure 7:**
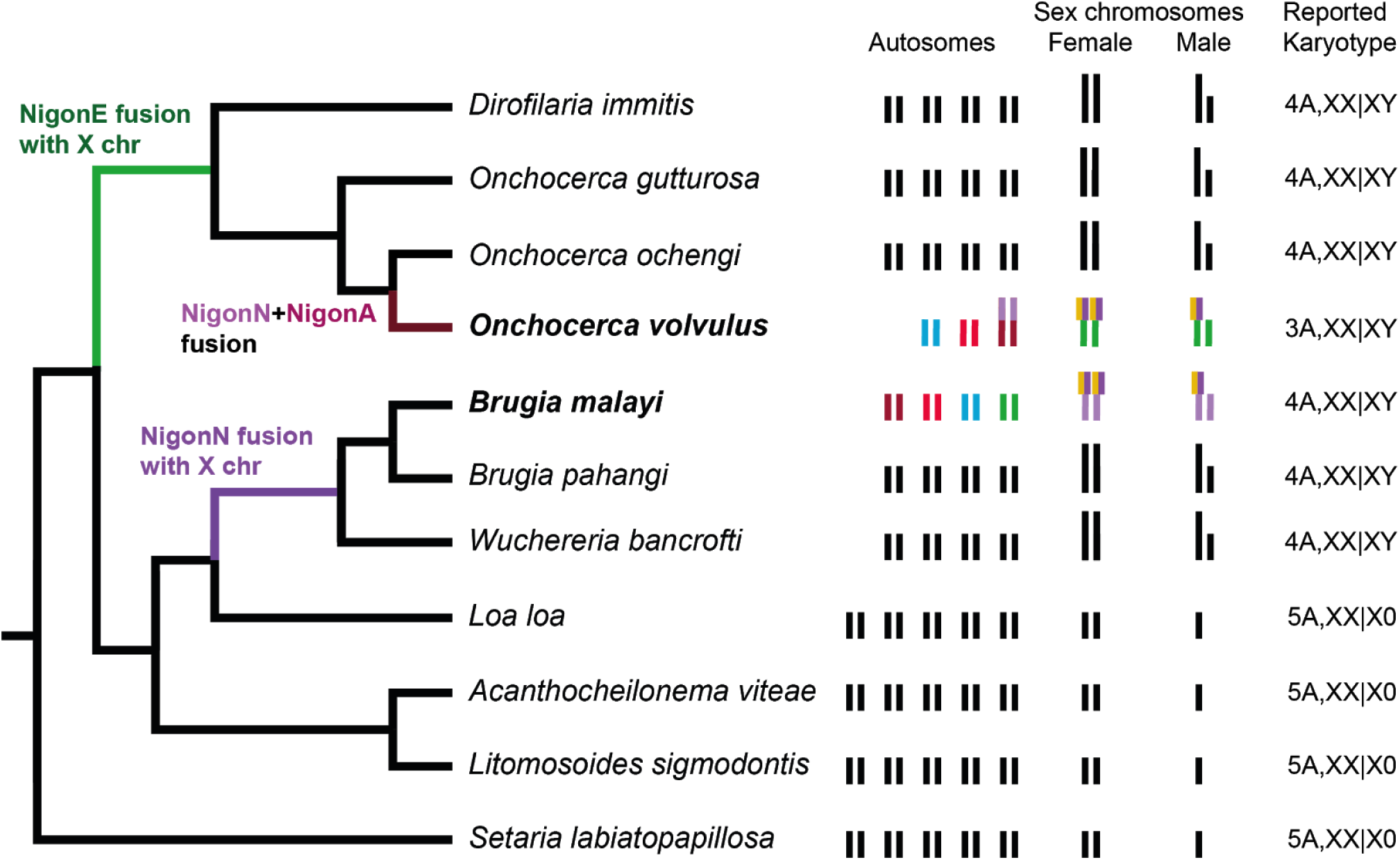
Sex chromosome evolution in the filarial nematodes (Spiruromorpha) Within the filarial nematodes, karyotypes have been determined for a number of species. The phylogeny (cladogram to the left) is derived from multilocus phylogenomic analysis, and is in agreement with marker gene-based phylogeny (Lefoulon et al., 2015). The karyograms and karyotype data are from the work of Rory Post and colleagues (Post, 2005). The inferred position of fusion events within the filarial phylogeny are indicated, and the karyotypes of *O. volvulus* and *B. malayi* are coloured by their Nigon assignment (see Figure 5 G,H and Figure 6).

While males in *B. malayi* and *O. volvulus* have a pair of sexually dimorphic chromosomes it is not clear whether the “Y” chromosome actually carries a male determining locus, or whether the system is a modified Xo system, as is found in the tylenchine *Strongyloides papillosus* (Albertson et al., 1979). In *Strongyloides ratti*, sex determination is XX:Xo, and the X chromosome is an intermixed fusion of NigonB, NigonD and NigonX. In the related *Strongyloides papillosus*, sex determination is also XX:Xo, but in this species haploidy of X is generated by intrachromosomal chromatin diminution. While the genome assembly for *S. papillosus* is not chromosomal, mapping of genetic marker loci indicated that the part that is lost is likely the NigonB-NigonD-NigonX component, while the remainder of the *S. papillosus* X appears to be homologous to *S. ratti* chromosome I (Nemetschke et al., 2010). Thus *S. papillosus* male-determining sperm have a “reduced-X” chromosome that contains only the autosomal part, and males are still diploid for the autosomal part. So are the *B. malayi* and *O. volvulus* sex determination systems XX:XY or apparent XY systems that biologically behave as XX:Xo?

Foster *et al*. (Foster et al., 2020) have argued, based on the presence on spiruromorph X chromosomes of NigonD loci (i.e loci mapping to *C. elegans* chromosome IV) that NigonD was the ancestral sex determination element of all Rhabditida, and that sex determination function transitioned to NigonX only later in rhabditid evolution. Foster *et al*. were unable to identify NigonN. Read data from males and females identified only a very small segment of genome (2.7 Mb in many short contigs) that was unique to male *B. malayi*, and a previously identified, male-linked locus (transposon on Y, TOY) was part of this male-limited genome. Karyotypic analyses identified the *B. malayi* X chromosome as being similar in size to the X chromosomes of XX:X0 species, and similar in size to the autosomes (Post, 2005). We interpret this sequence (and TOY) as being repeat accumulation in the subtelomeric regions of the short-form X chromosome that was lost consequent to the fusion event between the ancestral X and NigonN chromosomes, and doubt it has roles in sex determination. The NigonN component of the *B. malayi* X is diploid in males and females while the 12 Mb NigonD - NigonX portion is haploid in males.

Other spiruromorphs, including close relatives of *Onchocerca* and *Brugia* have an XX:X0 sex determination system (Post, 2005), suggesting this mode is ancestral. This pattern leads us to question whether sex determination in *B. malayi* and *O. volvulus* is in fact XX:XY, where, by analogy to XX:XY systems in other taxa, a sex (male) determining locus is present on the Y. In *B. malayi* and *O. volvulus*, where the non-NigonD-NigonX component of the X chromosome derives from fusion with different chromosomes, we propose that the same haploid-X mechanism operates, and that the apparent XX:XY sex determination system is in fact an XX:X0 system, where the additional component (NigonN in *B. malayi* and NigonE in *O. volvulus*) is always diploid. This must mean that the fusion partner, diploid in males and females, must be under distinct dosage compensation control compared to its sex-determining partner, as is likely the case in *S. papillosus*. Similar sex chromosome fusions where the newly fused parts have distinct dosage compensation mechanisms have been identified in another species with holocentric chromosomes, the butterfly *Danaus plexippus*. In *D. plexippus* the neo-Z chromosome has two distinct modes of compensation, spatially distributed along the fusion based on the origin of the segment (i.e. expression of the Z component, is halved in ZZ males, while expression of the autosomally derived fragment is doubled in WZ females (Gu et al., 2019).

### Functional coherence of loci that define Nigon elements

As these Nigon-defining loci have been colocated on the same karyotypic unit for much of rhabditid nematode evolution, we wondered whether each set had a functional coherence such that the colocated genes functioned together in specific pathways. We identified functional enrichment in five of the seven gene sets in *C. elegans* when the whole *C. elegans* gene set was used as comparator. As we selected these genes based on their largely one-to-one conservation across Rhabditida, it would be expected that they would be enriched in conserved function, and so this result is perhaps not surprising. When using only the set of Nigon-defining loci as a reference, however, we found functional enrichment only of NigonX-linked loci. Based on our model, the loci that define NigonX have been on the sex determination chromosome of rhabditid nematodes since the last common ancestor of Tylenchina, Spirurina and Rhabditina. These loci will thus have been exposed as haploid in males and will have had an effective population size of approximately 0.75 of the size of any autosomal locus for the entirety of rhabditid evolution. Genes with essential functions are more rarely found on the X chromosome than in the autosomes of *C. elegans*, while genes with non-lethal, post-embryonic knock-down phenotypes are enriched in the X chromosome (Kamath et al., 2003).

### Outlook

The telomere-to-telomere chromosomal assembly of *Oscheius tipulae* can now stand as a platform for future work on the developmental and population genetics of this important model species. Investigation of the biological importance of the telomeric extensions, and especially of their presence or absence in other species is of particular importance. Why do some genes stay on the same chromosome, while others appear to move freely? What mechanisms drive the processes of chromosomal structure, and why do some species have distinct patterns of intra- and inter-chromosome rearrangement? In Lepidoptera there is a general conservation of karyotype (with n=31) and genes tend to be situated on homologous chromosomes in the same order (i.e. there is strong conservation of micro- and macro-synteny), but some taxa diverge strongly from this pattern and have very different chromosome numbers (from n=4 to >200) (de Vos et al., 2020) that may not be simply described by fusion of whole chromosomes, or scission of chromosomes into multiple parts (Hill et al., 2019). In the Lepidoptera, karyotypic change is associated with speciation (de Vos et al., 2020), but whether this is also true in nematoda is not clear, though we note the relatively rapid karyotypic evolution in filarial nematodes (Figure 7) and the existence of genera and orders such as *Diploscapter* in Rhabditina (n=1 to 7) (Fradin et al., 2017) and the Ascaridomorpha (n=1 to 24), where chromosome counts vary greatly (Walton, 1959). We look forward to an increase of chromosomal assemblies from rhabditid and other nematodes in the near future to further explore patterns and processes in nematode chromosome evolution.

## ACKNOWLEDGEMENTS

PG is funded by a PhD Scholarship from the Darwin Trust of Edinburgh. PromethION sequencing at Edinburgh Genomics was funded by a UK Natural Environment Research Council Biomolecular Analysis Facility contract. Sophie Tandonnet was funded by CAPES/CNPq (201116/2014-6) and FAPESP (2019/07285-7). We acknowledge the assistance of our colleagues at Edinburgh Genomics and of members of the University of Edinburgh Institute of Evolutionary Biology evolutionary genomics community, especially Lewis Stevens and Andrea Martinez Martinez. The *O. tipulae* CEW1 strain was supplied by Marie-Anne Félix, and was originally isolated by Carlos Winter. We also thank Tom Freeman for discussions about clustering, and Jianbin Wang and Richard Davis for early access to the chromosomal genome assembly of *Ascaris suum*.

## Supplementary Information

### Supplementary Text S1: Defining sets of orthologous loci in linkage through Rhabditid evolution

We identified sets of loci that potentially define ancestral linkage groups - Nigon elements - by analysing the chromosomal colocation of sets of orthologues across nine chromosomally assembled nematode genomes. The nine genomes analysed were *Auanema rhodensis, Brugia malayi, Caenorhabditis elegans, Haemonchus contortus, Onchocerca volvulus, Oscheius tipulae, Pristionchus pacificus, Steinernema carpocapsae* and *Strongyloides ratti* (see Table S4). Proteomes of these nine species were clustered into orthogroups using orthofinder (Emms and Kelly, 2019). The orthofinder output was imported into KinFin (Dominik R Laetsch and Blaxter, 2017), and built in KinFin routines used to select sets of orthologues that conformed to particular presence-absence patterns. One to one orthologues, or single copy orthologues are orthologue sets where all the species in the analysis have exactly one copy. We defined fuzzy one to one orthologues, where an orthologue set was selected if it contained at least a given number of species with one copy in each, and the remaining species were permitted to be missing the locus (count of 0) or to have more than one copy. This allows fuzzy one to one orthologues to be defined even when one or a few genomes have experienced gene loss, where there are idiosyncratic gene duplications (or uncollapsed retained haploid duplications) or where the gene set or genome is incomplete. We explored the landscape of fuzzy one to one orthologue definition and chose a conservative cutoff of requiring seven of the nine input species to have single members, and allowing two species to have 0 or >1.

For each of these orthologues we recorded which chromosome it was placed on in each species, yielding a matrix where each orthologue was described by an array of chromosomal assignments. Missing and duplicate orthologues were assigned null values. Gower distances between orthologues were calculated from these categorical data, yielding a normalised distance matrix relating all orthologues across all species. This distance matrix was examined using t-SNE with different perplexity values and Graphia Pro using different correlation cutoff values We identified seven clusters of orthologues (Figure S4). CLARA clustering was used to cluster the orthologues by Dice distance, exploring a range of values of K from 1 to 10. Clustering with K=7 had the best scores in terms of mean silhouette width (Figure S3), and these seven clusters were used to define sets of loci that are associated with seven ancestral Nigon elements.

We explored the robustness of definition of these seven Nigon-defining orthologue sets by exploring the effect of the 1-to-1 orthologous relationship in different number of species used in identifying them (Figure S8). When low numbers of species were analysed, identification of seven elements was not achieved, and elements N and X tended to be merged. However for all analyses using over three species in the analysis, the numbers of orthology groups assigned to each putative Nigon-defining set was relatively constant. When the genomes with many chromosomes (*A. suum*, *M. hapla*) or the less-well assembled genomes were used, the number of Nigon-defining loci was impacted.

### Supplementary Text S2: Nigon element analysis of *Meloidogyne hapla*

#### Introduction

The genus *Meloidogyne* includes both diploid species and major clades of polyploid hybrids, with karyotypes of n=17 in diploids and n of up to 70 in polyploids. *Meloidogyne hapla* is diploid and has 17 chromosomes, defined by genetic mapping and karyotyping (Thomas et al., 2012). The genome assembly has been superscaffolded into 19 scaffolds using genetic cross mapping, with chromosomes 1 (mh_1A, mh_1B) and 2 (mh_2A, mh_2B) split in two. Each single superscaffold (e.g. “mh_10.12.8”) corresponds to a single genetic linkage group. There are 275 short, unplaced scaffolds. The superscaffolds were analysed in detail, with the split chromosomes 1 and 2 kept as two fragments.

#### Methods

Nigon-unit defining loci were identified in *Meloidogyne hapla* and their mapping to superscaffolds recorded.

#### Results

Compared to the other nematode species analysed, the published genome sequence for *M. hapla* had a noticeably lower proportion of matches to the set of shared orthologues (Figure 6 B,C). Of 2191 Nigon-defining loci, 1688 (or 77%) had matches in the *M. hapla* genome. The mapping rate was lowest for Nigon set X (68%) and highest for Nigon set B (81%) (Table S12). Three quarters (75%) of the mapped loci from the Nigon-defining sets were present in the chromosome-sized contigs, with the highest proportion of Nigon set matches in these unplaced contigs from the X set (38%). Most of the unplaced scaffolds had a single Nigon locus mapped (range 1-17, mean 1.54).

The number of Nigon-defining loci mapped to each chromosomal molecule ranged from 8 to 157. The two chromosome 1 contigs had a total of 201 loci mapped and the two chromosome 2 contigs had a total of 144 loci mapped. Four of the shorter chromosome-mapped scaffolds had <40 Nigon-defining loci mapped, and these were not analysed further. Assessment of possible assignment to ancestral Nigon elements using simple counts or proportions of mapped Nigon-defining loci (Table S12) biased assignment to those Nigon elements that contained more loci (e.g. the number of loci used to define Nigon X is less than one sixth the number used to define Nigons A and C). Thus we normalised Nigon-defining locus counts per chromosomal scaffold by expressing them as the proportion of Nigon-defining loci from each Nigon set.

Using normalised proportions, a few chromosomal superscaffolds could be assigned to ancestral Nigon elements with some confidence. Thus chromosome mh_16 contained 50% of all the NigonE set loci mapped to the chromosomes, and chromosome mh_8 contained 38% of the mapped NigonX set. Other chromosomes showed a bias towards one or a few Nigon-defining sets. Chromosome mh_5 had overrepresentation of NigonB, mh11 of NigonN, mh4 of NigonC, mh5 of NigonB and mh_5 of NigonX loci. Notably, the assignments of the two halves of chromosome 1 (mh-1A and mh_1B) were congruent (highest proportion to NigonA in both), while the mappings of mh_2A and mh_2B from chromosome 2 were discordant (highest proportions to NigonB and NigonD respectively).

## Discussion

Overall, the assignment of *M. hapla* chromosomal superscaffolds to Nigon origin is complex. No scaffold was simply assigned to one Nigon, but only seven had content where one Nigon set constituted a majority of mapped loci, and these counts were biased by the count of loci in each set. In *Strongyloides ratti* and *Steinernema carpocapsae*, we identified a shared fusion of NigonB, NigonD and NigonX to form the X chromosome of each species, with additional complex rearrangements in *S. ratti*. In the *M. hapla* assignments there is no signal of this predicted fusion as NigonX-derived loci are not particularly associated with either NigonB or NigonD loci. The two scaffolds on which a majority of NigonX loci were mapped (mh6 and mh8) had very few NigonD loci (zero and 1 loci respectively). The additional fusions predicted in *S. ratti* (fusions of C+E+N and of A+C+D+E+N) were also not evident.

Thus we conclude that, if the chromosomes of *M. hapla* contain a remnant signature of their derivation from ancestral Nigon elements, the fragmentation implied in going from seven ancestral to seventeen extant chromosomes also involved a large number of translocations. It will be very informative to examine additional high-quality genomes from additional *Meloidogyne* species, and other chromosomally-contiguous genomes from the Heteroderidae. Many *Meloidogyne* species are complex triploid and tetraploid hybrids, making assembly and interpretation difficult, but recent progress with long read data generation shows promise in generation of high quality genome estimates (Susič et al., 2020; Szitenberg et al., 2017).

---

**Figure S1:**
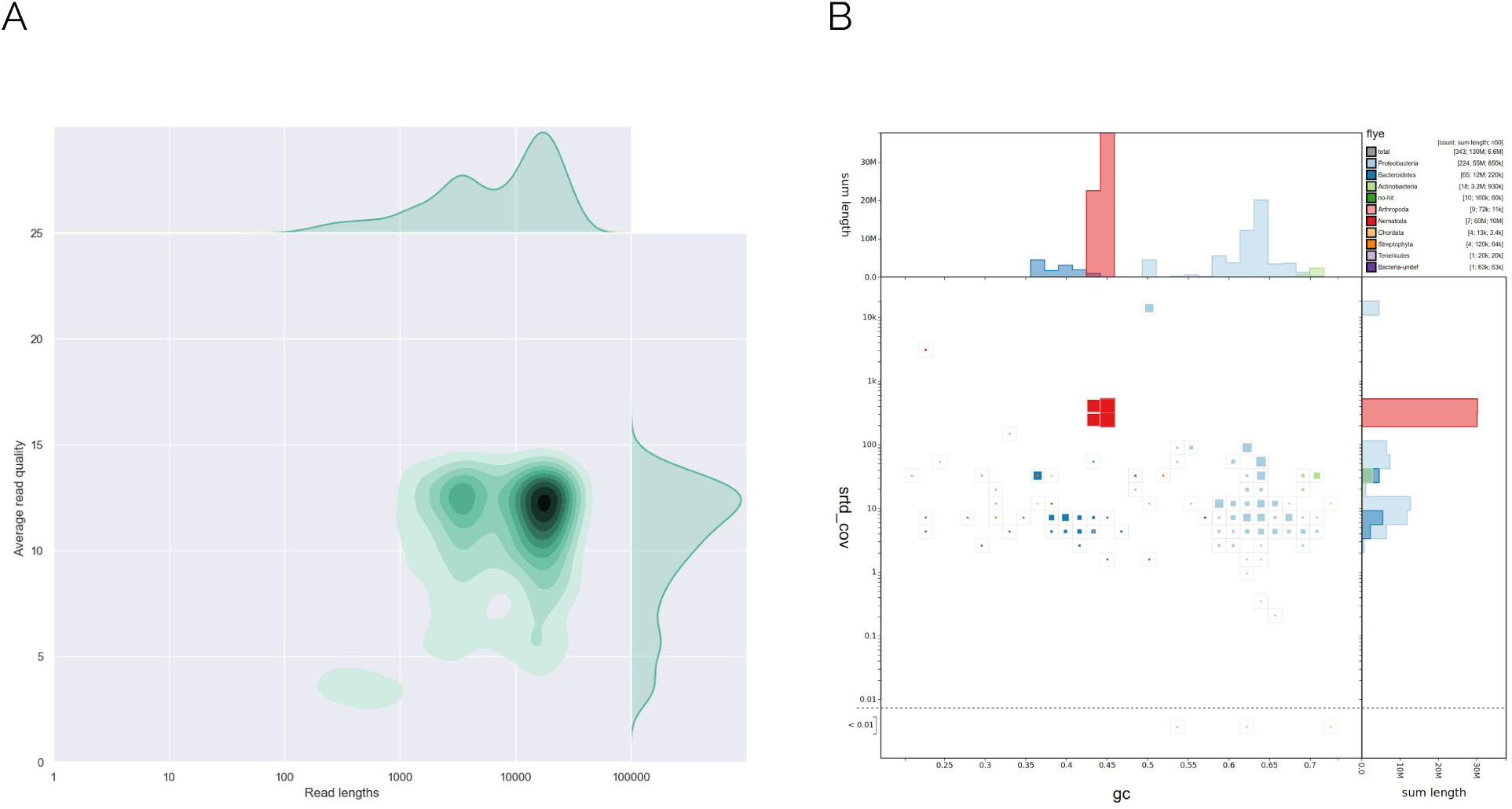
Raw data QC and read filtering. A. Density plot of raw PromethION nanopore read length (x axis) against average read quality (y axis) (generated by Nanoplot). B. BlobToolKit (Challis et al., 2020) plot (GC proportion [x axis] vestus read coverage [y axis]) of the flye assembly of all the sequenced reads. Taxonomic annotation was achieved using blastx and Diamond tblastn source classification.

**Figure S2:**
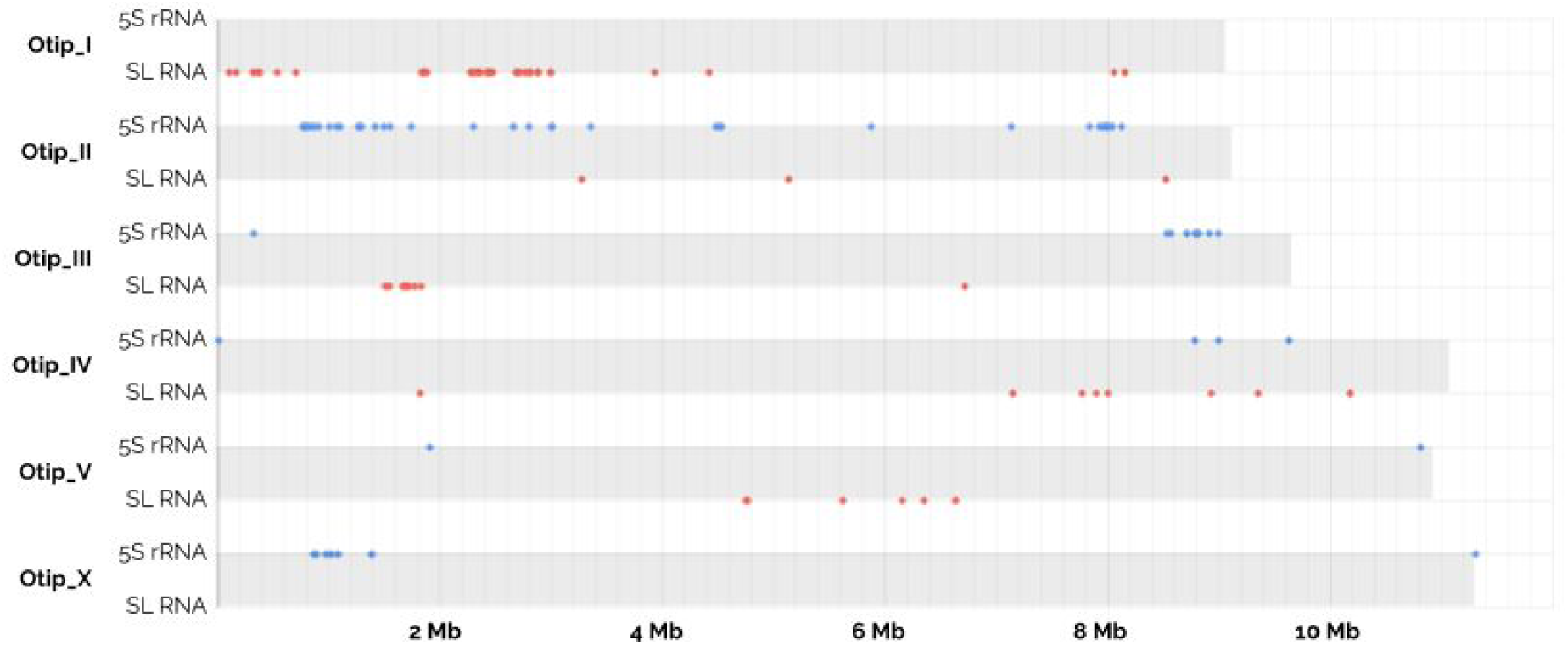
Distribution of SL and 5S RNA loci in the Oscheius tipulae genome. Spliced leader RNA and 5S ribosomal RNA loci were identified using Rfam (Kalvari et al., 2018) models and Rnammer (Lagesen et al., 2007), and are plotted by position along each of the *O. tipulae* chromosomes.

**Figure S3:**
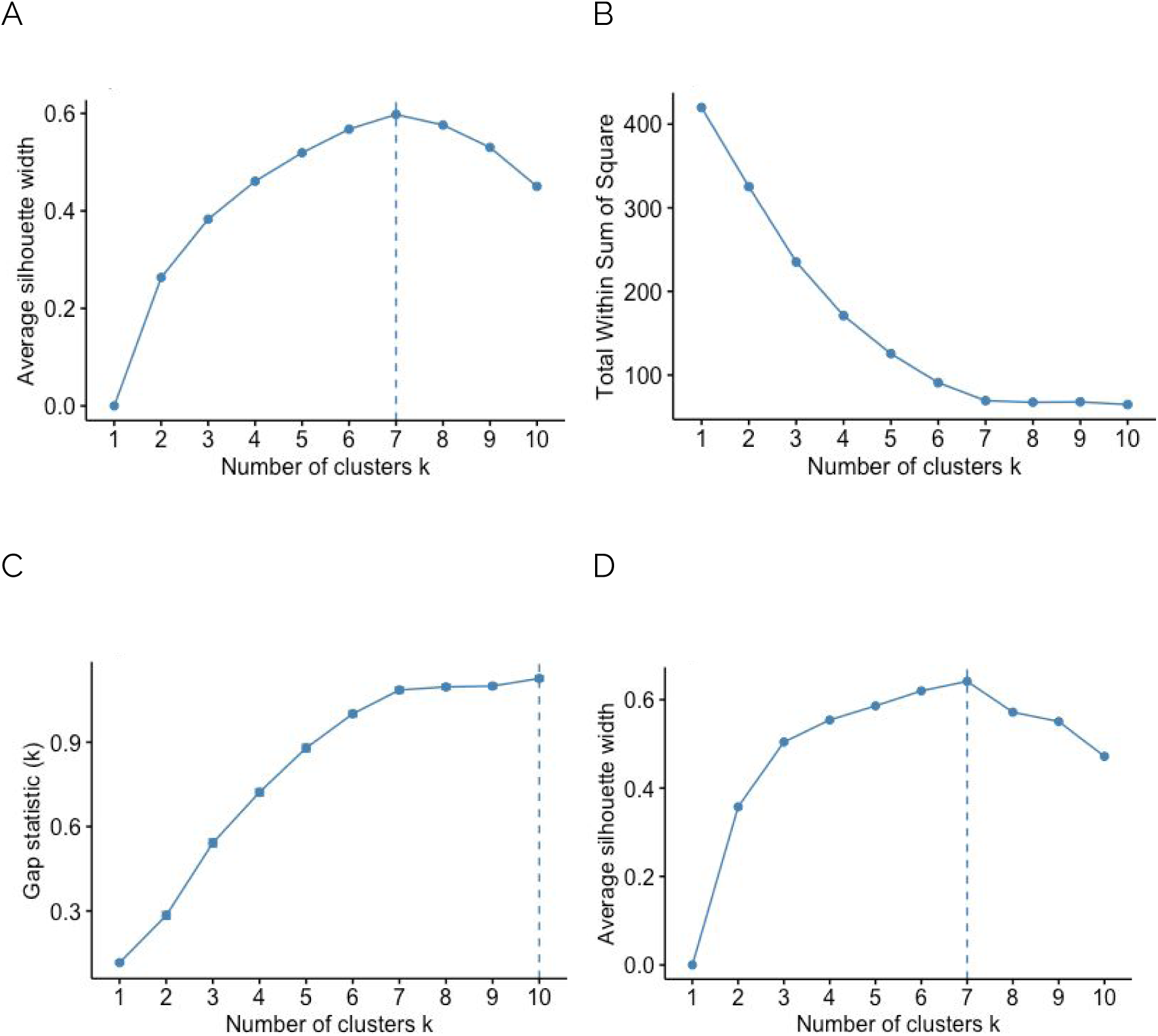
Clustering with different k values. Optimal number of clusters between 1 and 10 using CLARA (A, B and C) or PAM (D) on the gower distance matrix of 1172 single copy orthologous genes. Silhouette (A and D), within sum of square (B) and gap statistic (C) were used to estimate the optimal number of clusters. One hundred Monte Carlo samples were performed for the gap statistic.

**Figure S4:**
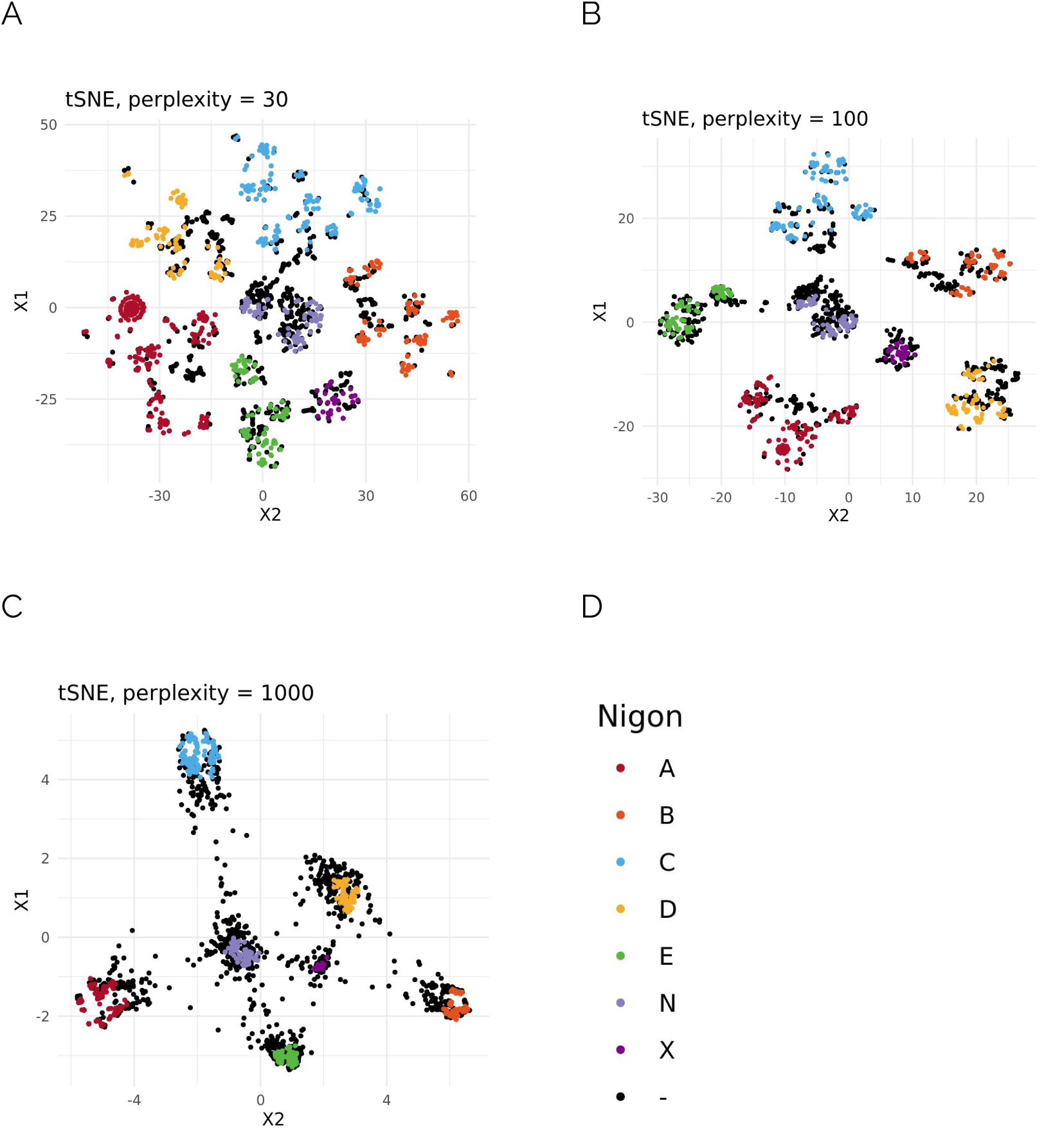
t-SNE of orthologous loci co-locations in 9 nematode species. Alternative t-SNE plots of 3412 orthologous gene families with different perplexity parameter values. A maximum of 1000 iterations were performed in all cases. Black dots denote orthologs not assigned to a Nigon unit. The legend in panel D applies to all tSNE panels. Parameters used for t-SNE, apart from the specified perplexity, are max_iter=1000, initial_dims=50 and theta=0.5.

**Figure S5:**
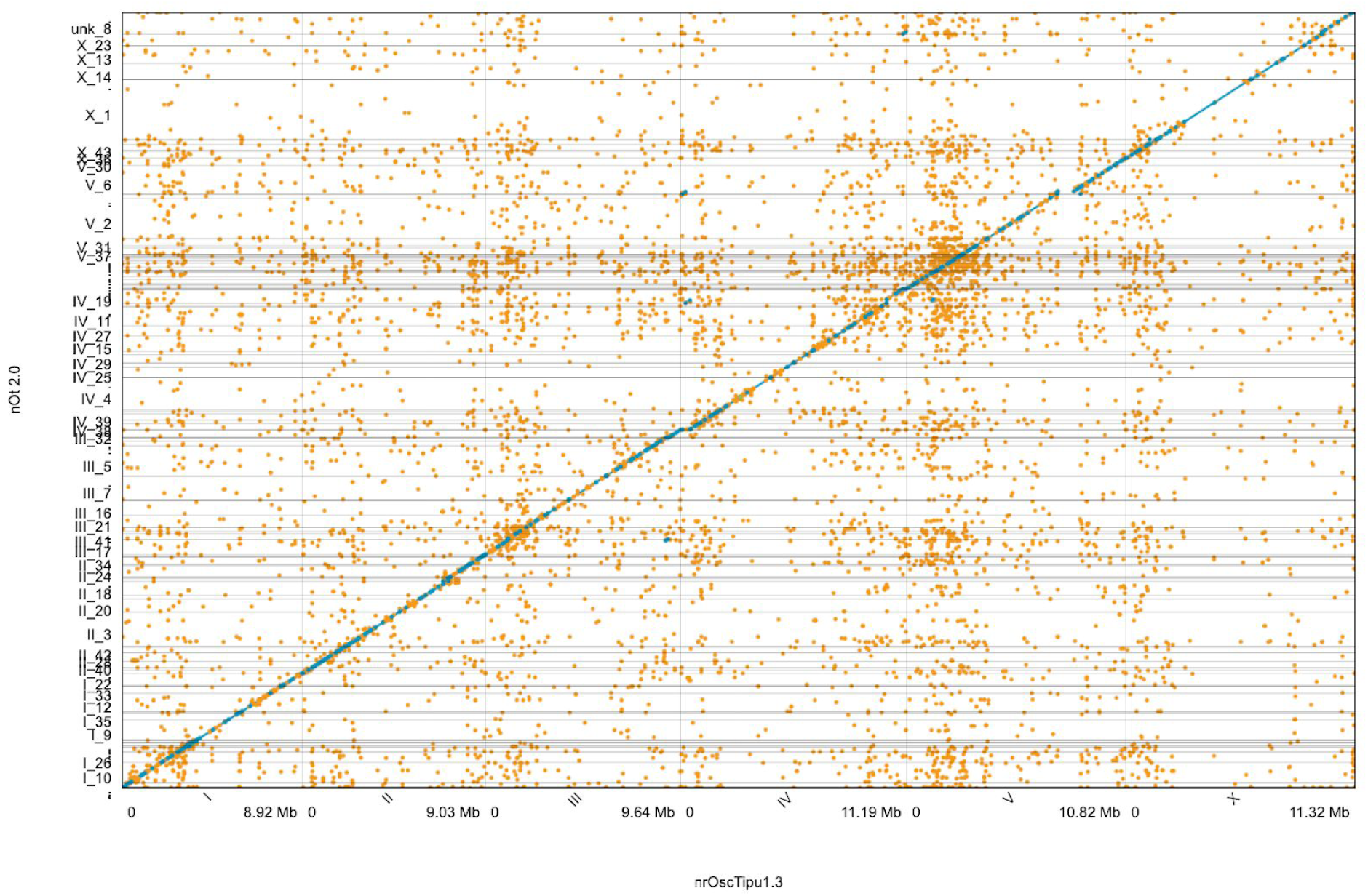
nOti 2.0 assembly aligned to nrOscTipu1.3. The telomere-to-telomere genome assembly (x axis) aligned to the previous assembly (Besnard et al., 2017) (y axis) in nucmer (Kurtz et al., 2004). Blue dots indicate co-oriented nucleotide matches, while orange dots indicate matches in opposite orientations.

**Figure S6:**
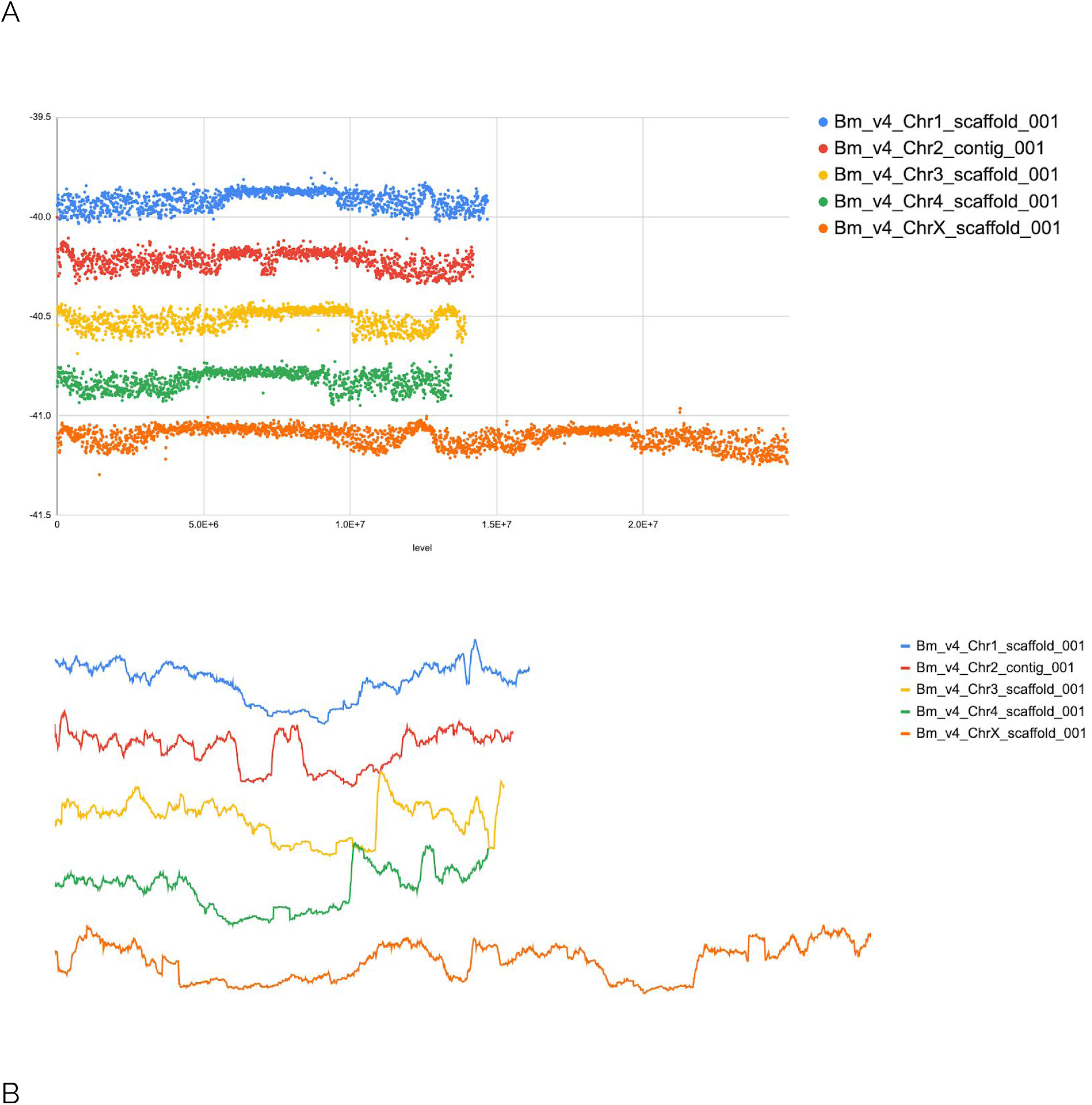
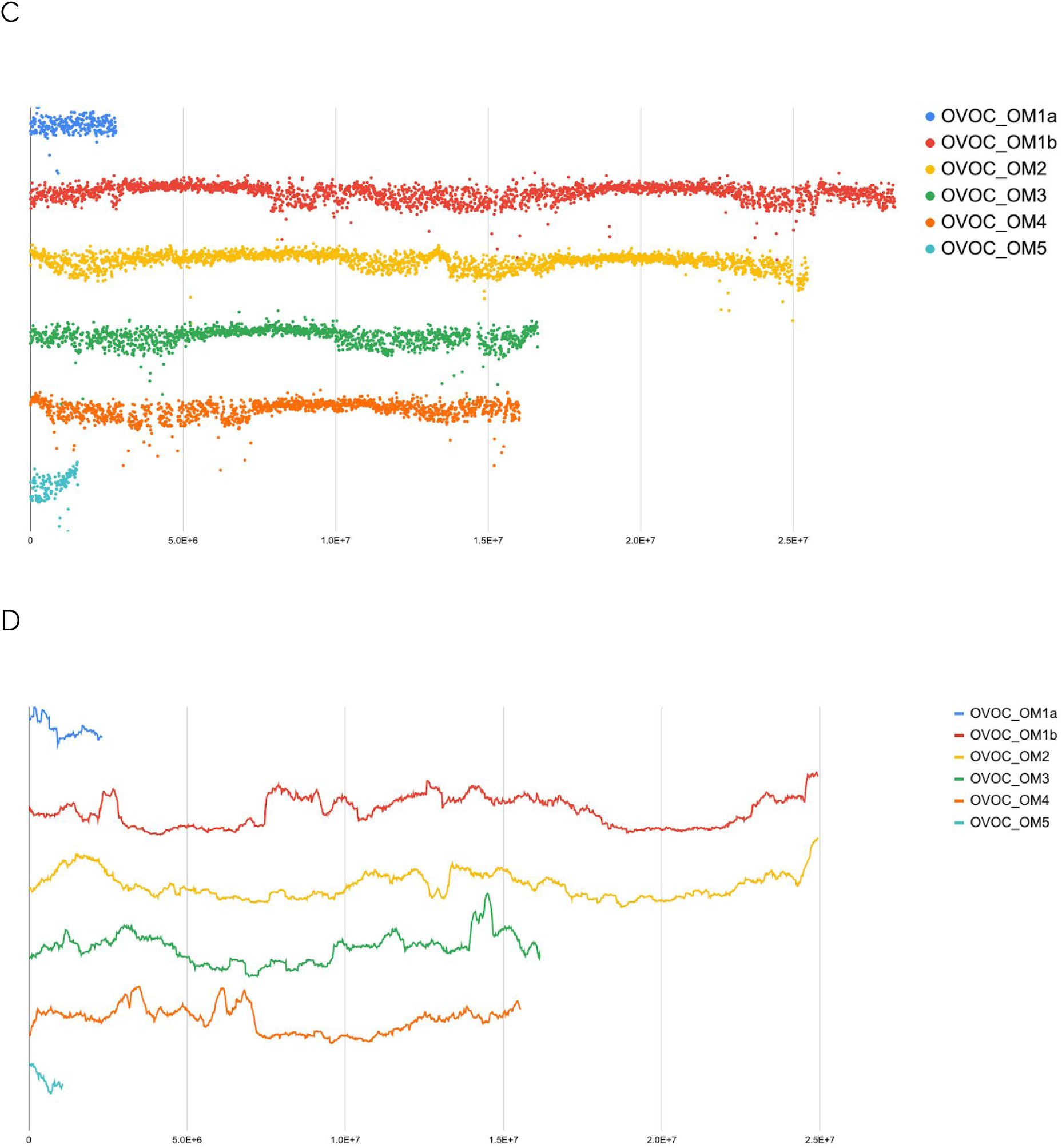
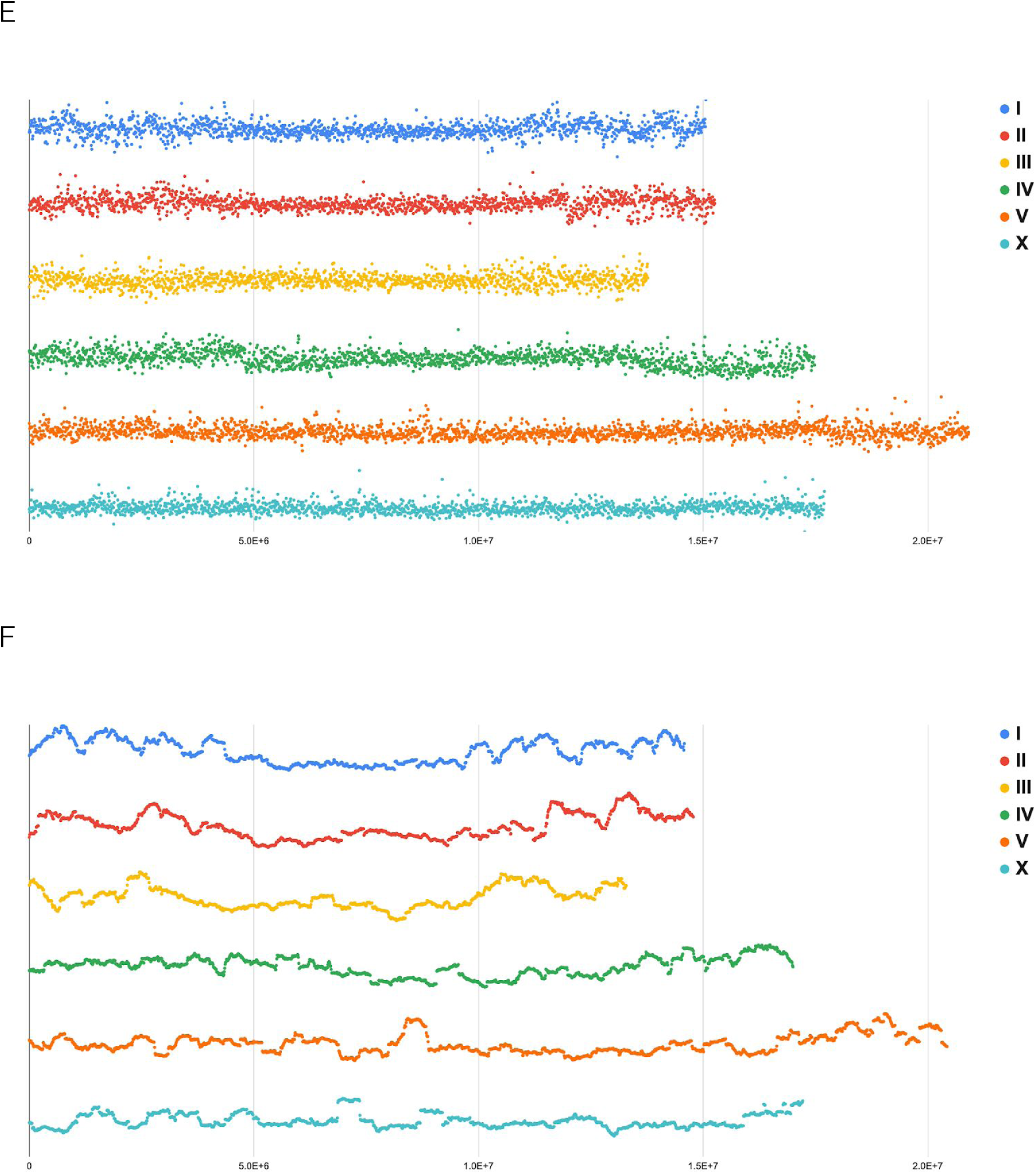
GC proportion plots of nematode chromosomes. A *B. malayi* GC proportion in 10kb windows across the chromosomes. B *B. malayi* GC proportion standard deviation in 0.5 Mb sliding windows across the chromosomes. C *O. volvulus* GC proportion in 10kb windows across the chromosomes. D *O. volvulus* GC proportion standard deviation in 0.5 Mb sliding windows across the chromosomes. E *C. elegans* GC proportion in 10kb windows across the chromosomes. F *C. elegans* GC proportion standard deviation in 0.5 Mb sliding windows across the chromosomes.

**Figure S7:**
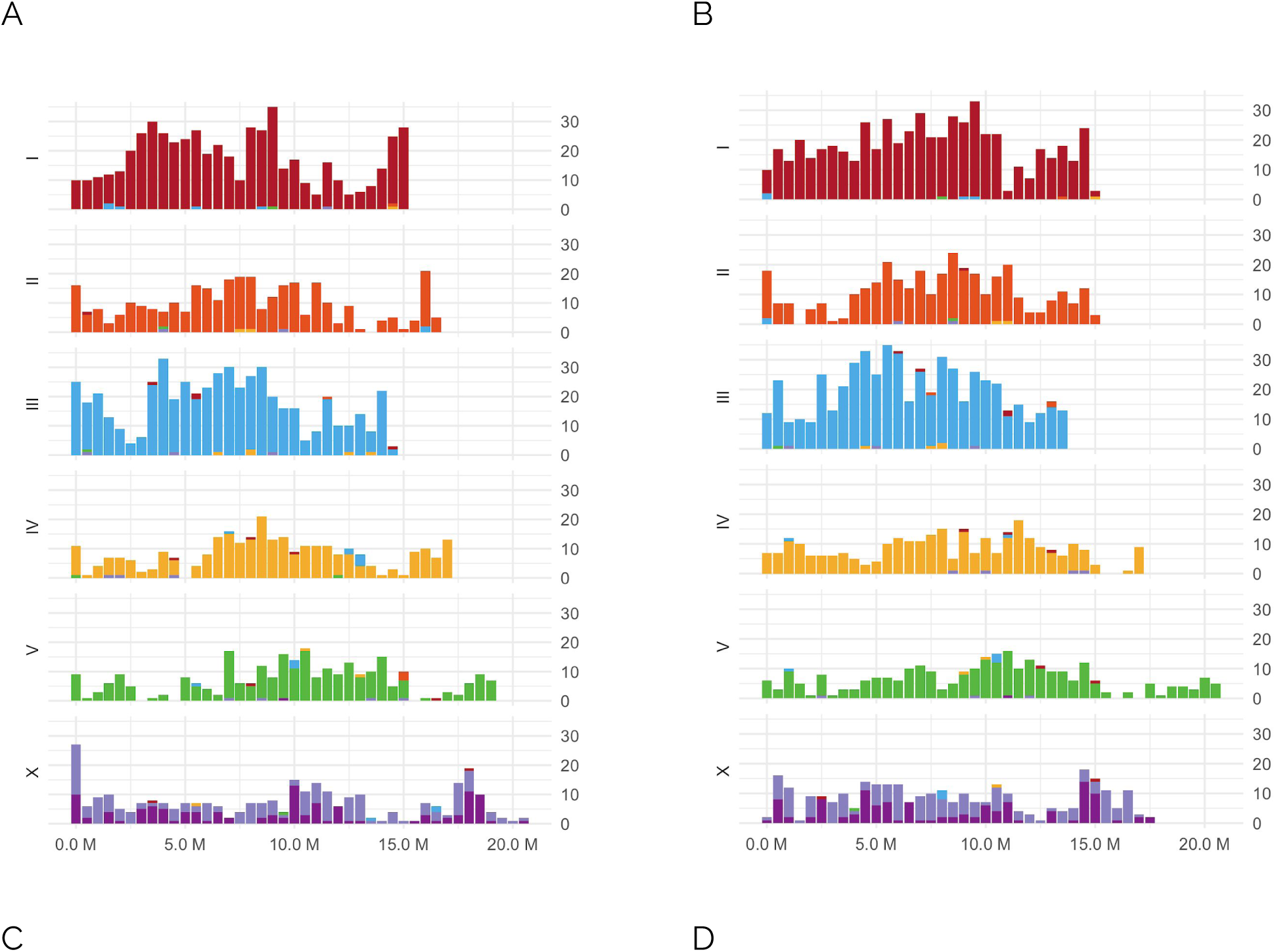
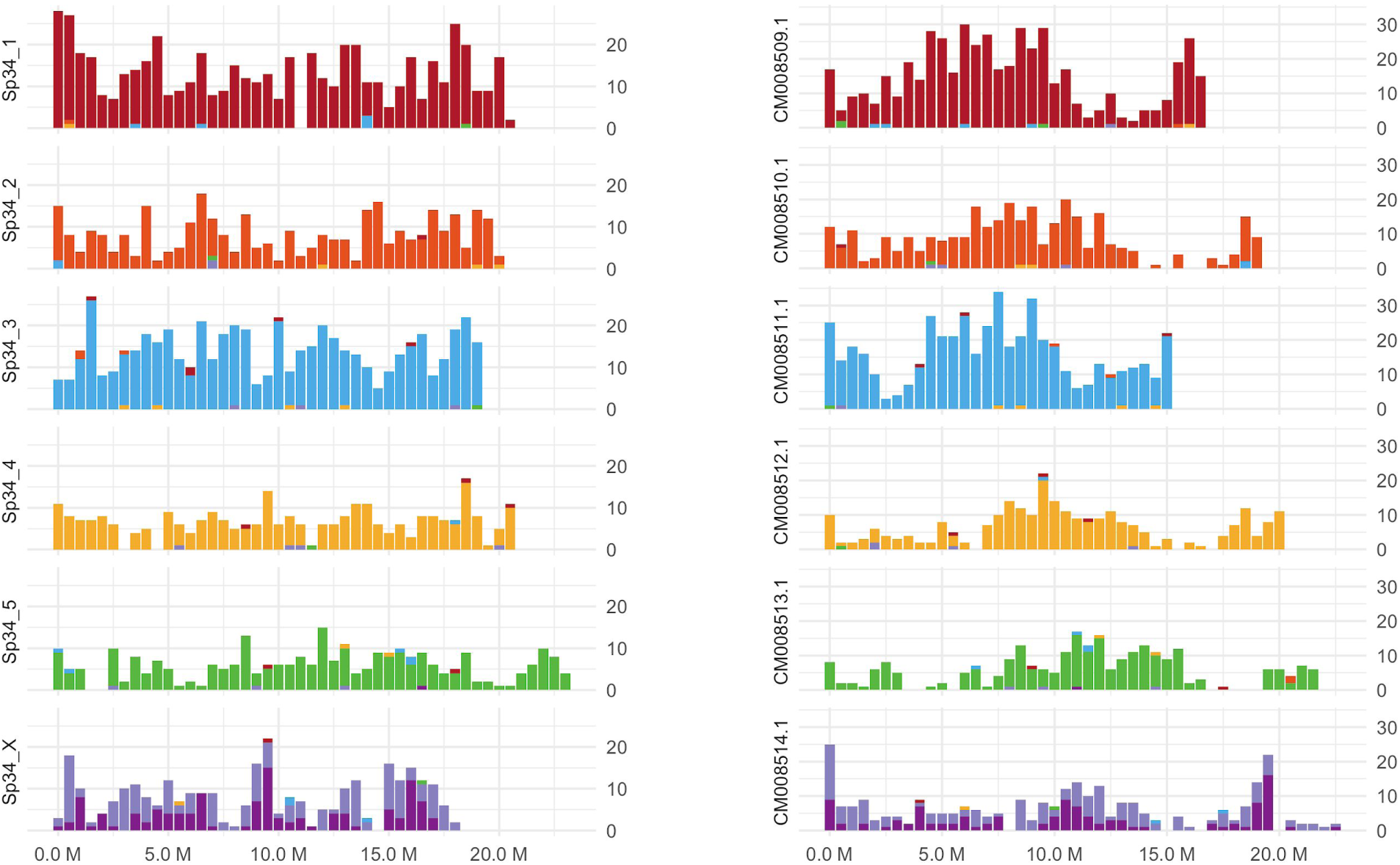
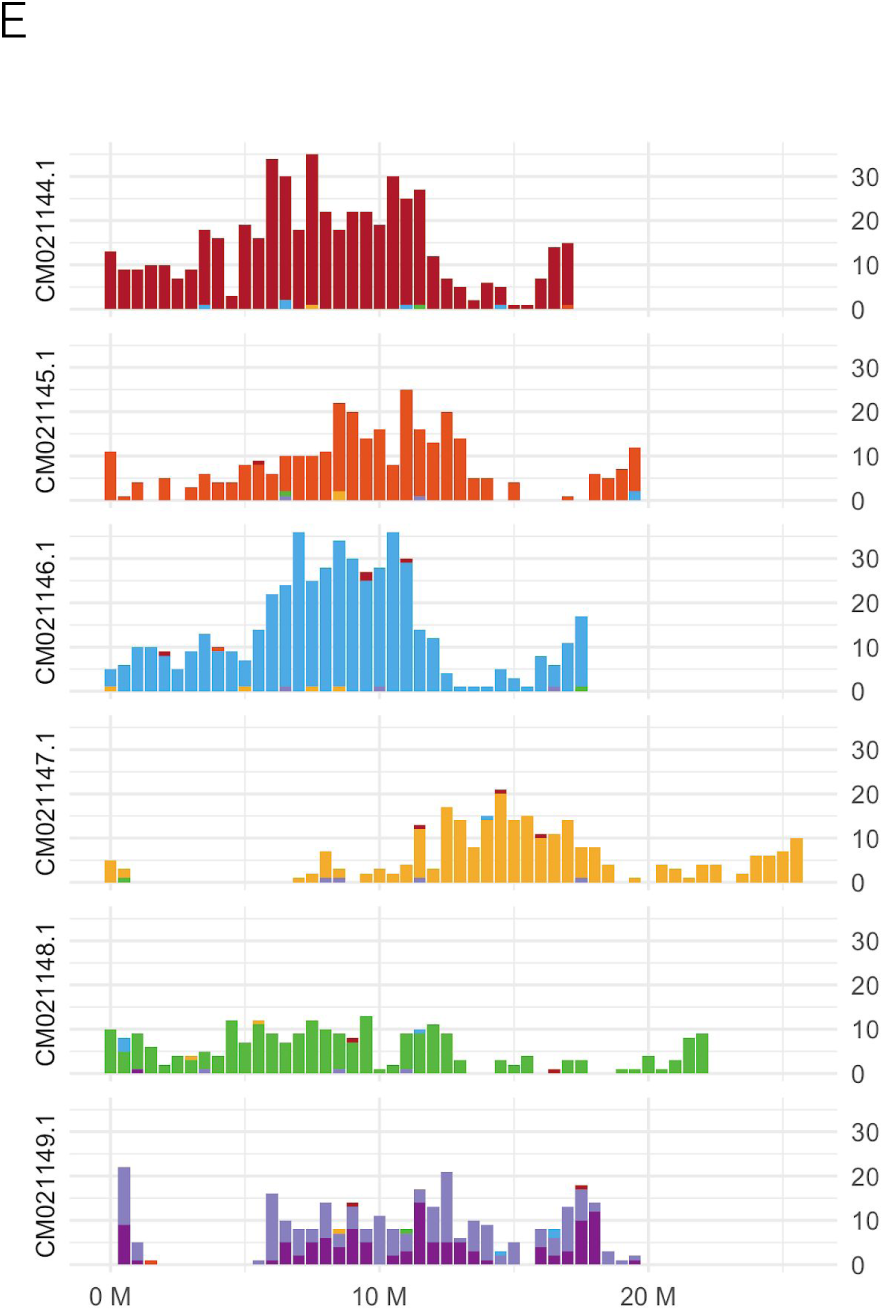
Nigon painting of *Caenorhabditis* species. Chromosomal assemblies of *Caenorhabditis* species painted using the Nigon-defining locus sets. See main text Figure 5 for other species. A *Caenorhabditis elegans,* B *Caenorhabditis briggsae,* C *Caenorhabditis inopinata,* D *Caenorhabditis nigoni*, E *Caenorhabditis remanei*.

**Figure S8:**
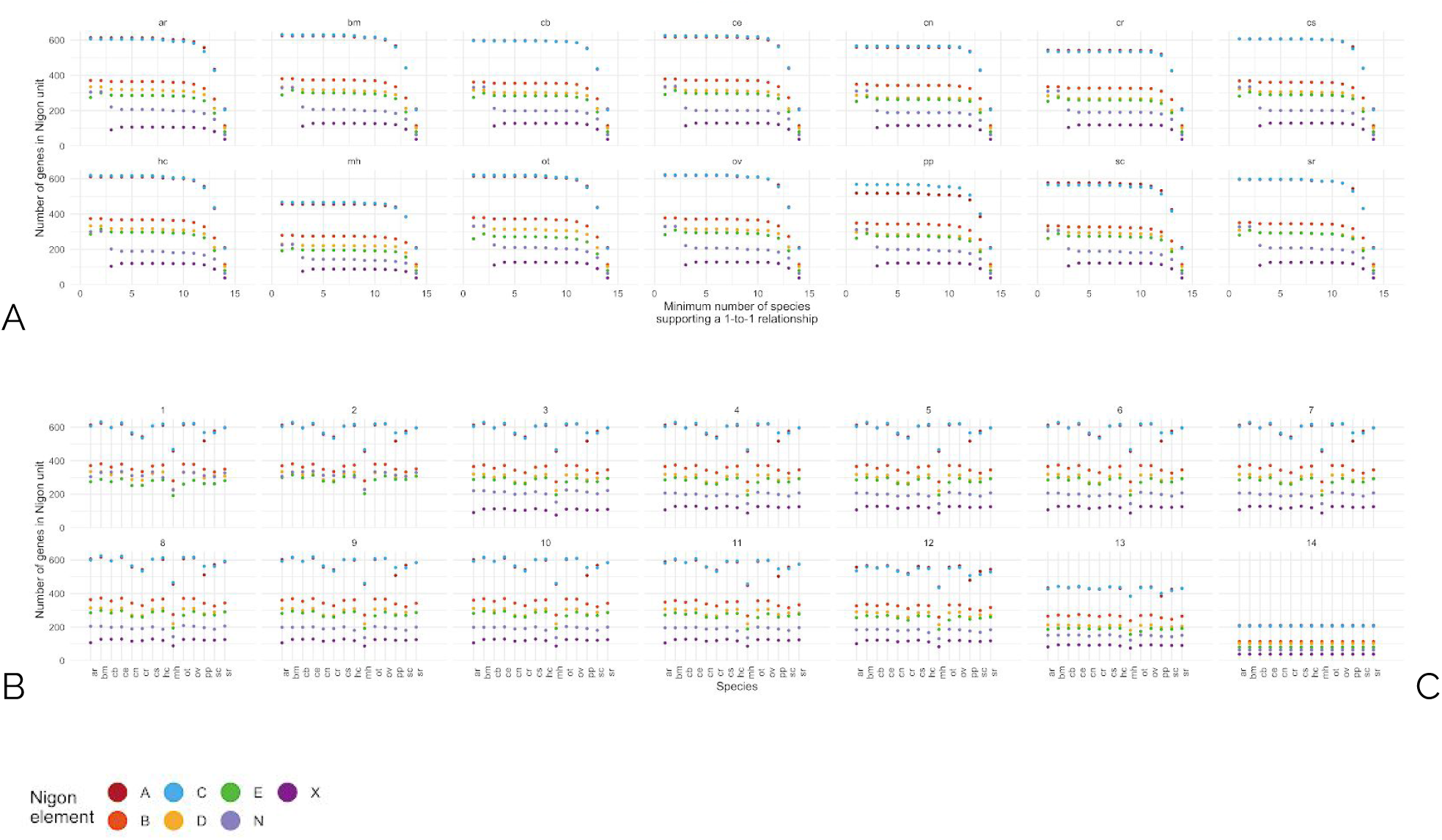
Exploring the robustness of definition of Nigon element-defining sets of orthologues. A Numbers of orthologous gene sets assigned to each putative Nigon-defining set in analyses using different numbers of species (from 2-14) as the minimum number of species having a 1-to-1 orthologous relationships input to the K-means clustering. Species abbreviations are ar: *Auanema rhodensis,* bm: *Brugia malayi,* cb: *Caenorhabditis briggsae*, ce: *C. elegans*, cn: *C. nigoni*, cr: *C. remanei*, cs: *C. sp34*, hm: *Haemonchus contortus,* mh: *Meloidogyne hapla,* ot: *Oscheius tipulae,* ov: *Onchocerca volvulus,* pp: *Pristionchus pacificus,* sc: *Steinernema carpocapsae* and sr: *Strongyloides ratti* B Same data as A, grouped by species rather than by minimum number of species having a 1 to 1 orthologous relationship. The number of orthologues assigned is stable in each species between K=3 and K=10. C Legend corresponding to panels A and B.

**Table S1:** Chromosome numbers in Nematoda*.

| **Clade** | **Subgroup** | ***n* in majority** | ***others*** |
| --- | --- | --- | --- |
| Clade I | Trichocephalida | 3, 4 |  |
| Clade II |  | ? |  |
| Clade C | Desmodoridae | 7 |  |
| Clade III | Oxyuromorpha | 4, 8 |  |
| | Ascaridomorpha | variable | highly variable: $n = 1-24$ |
| | Spiruromorpha | 5, 6 | rarely $n = 4, 7$ |
| Clade IV | Strongyloides | 3 |  |
| | Meloidogyninae | 17 | parthenogens variable to $n > 50$ |
|  | Pratylenchus | 5 - 7 |  |
| | Panagrolaimus | 4 | triploid parthenogens $n = 12 = 3 \times 4$ |
| | Steinernema | 6 | also $n = 8, 10$ |
| Clade V | Strongylomorpha | 6 |  |
| | Rhabditomorpha | 6 | varies $n = 1 - 9$ (especially variable in *Diploscapter*) |
| | Diplogasteromorpha | 6 | one with $n = 5$ |

\* See (Walton, 1959) for a summary of chromosome numbers in nematode parasites.

**Table S2:**
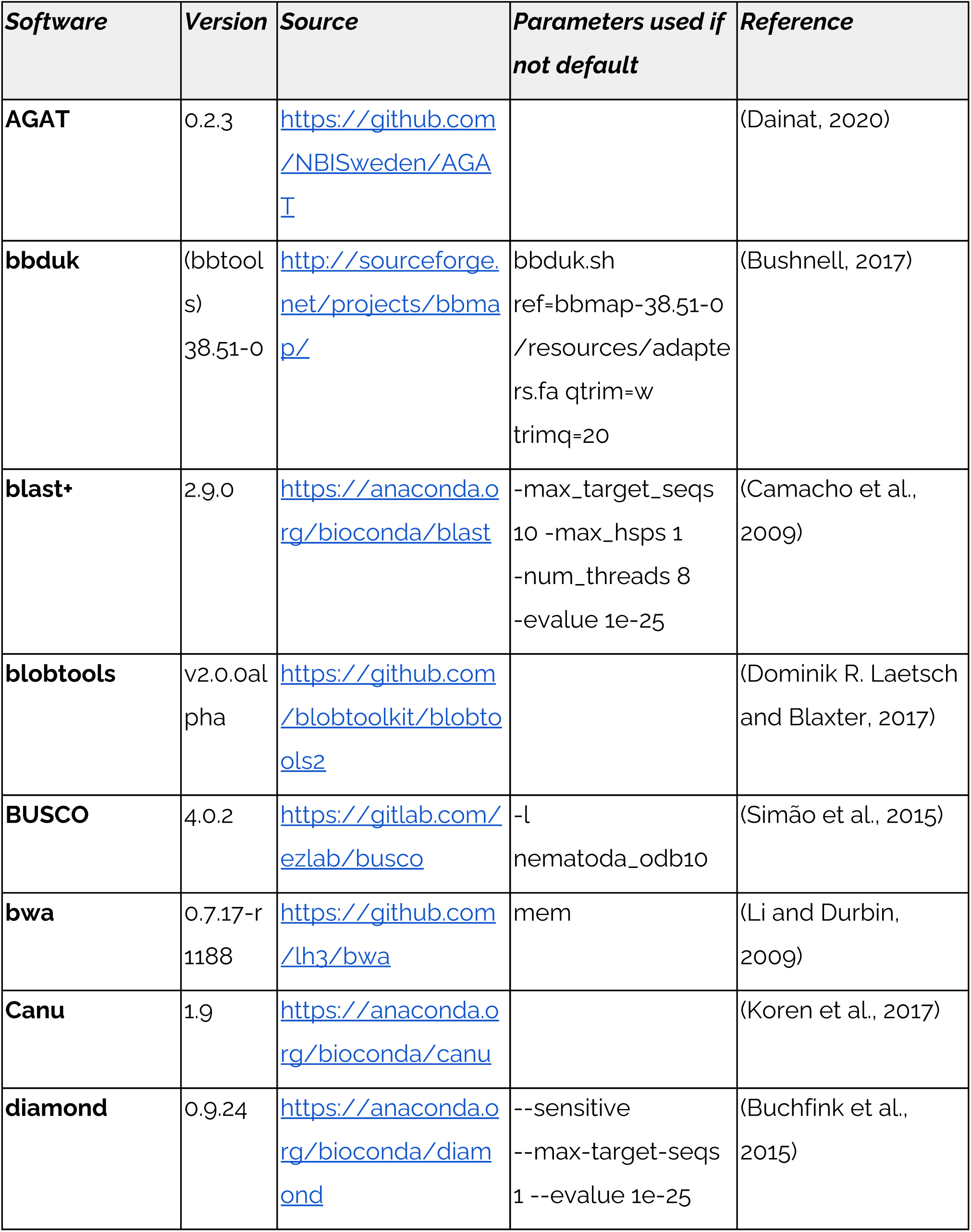
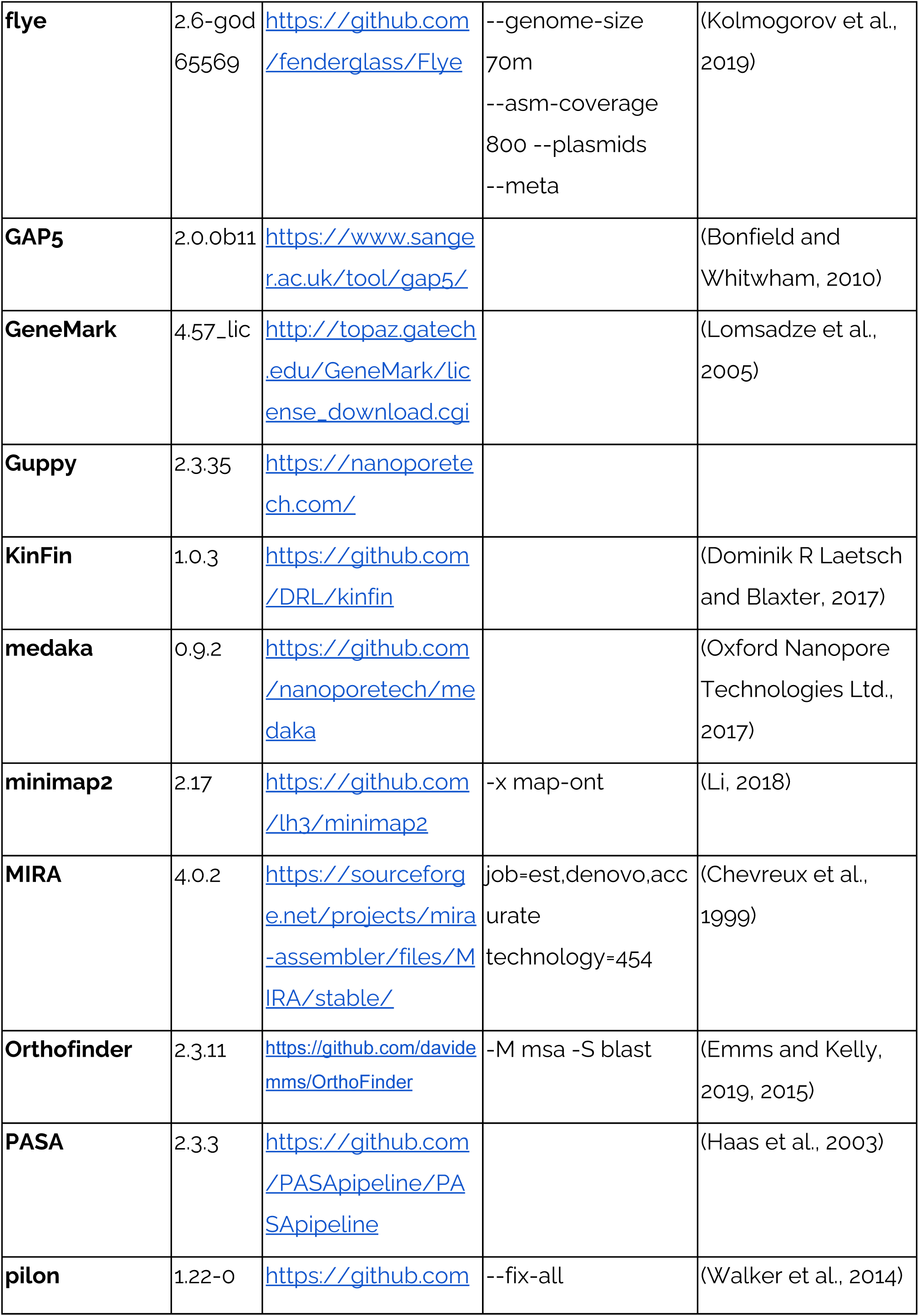
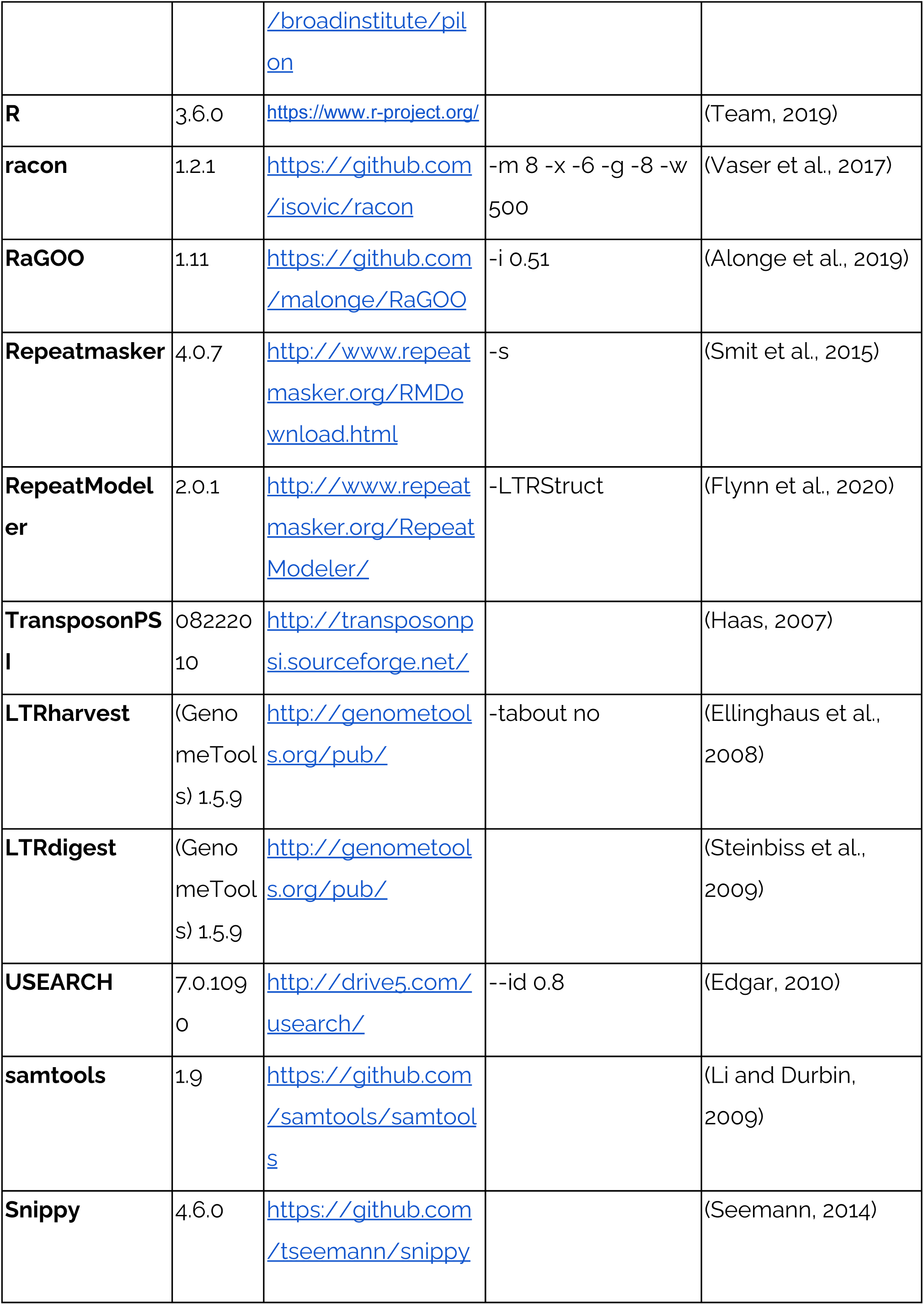
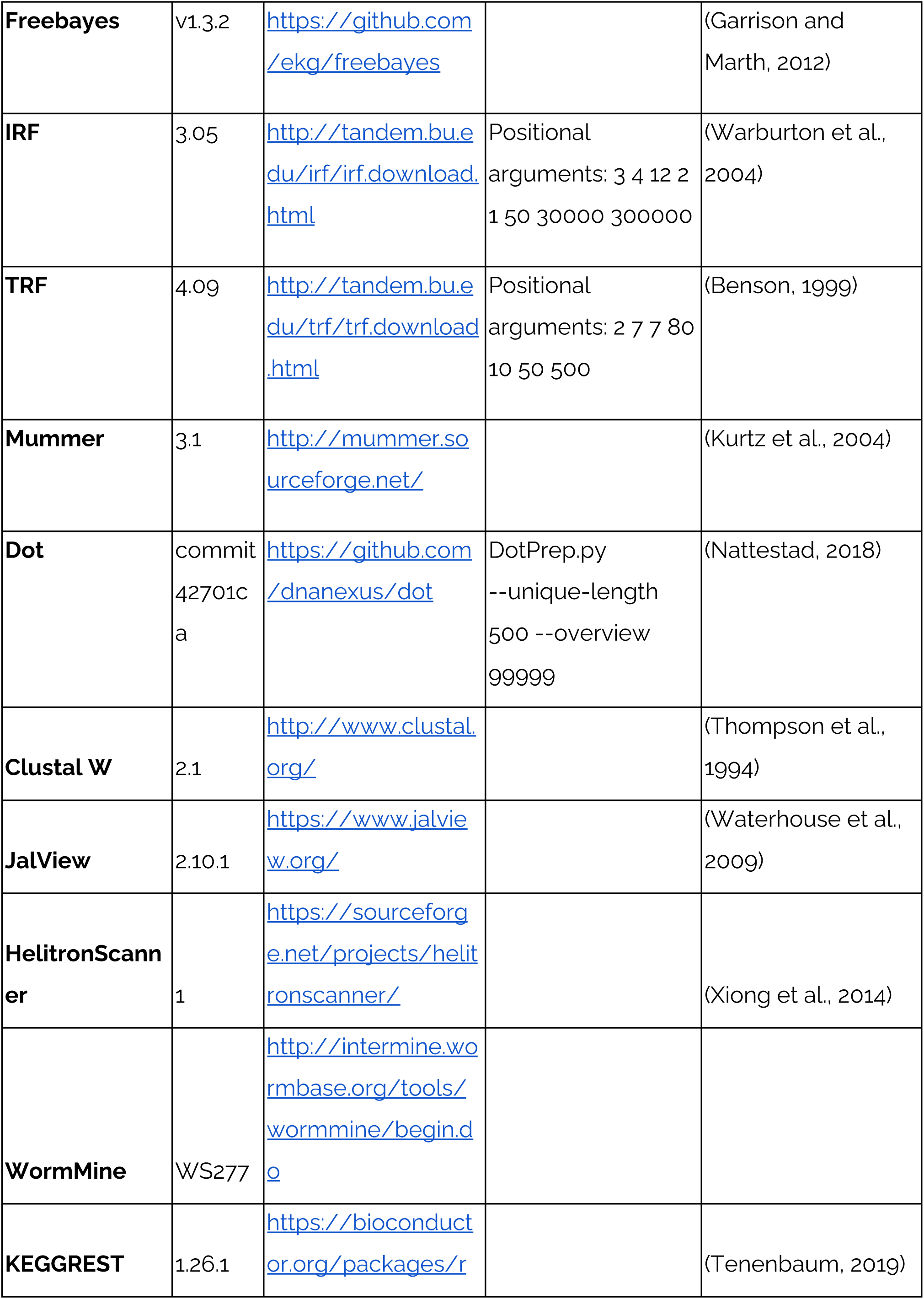
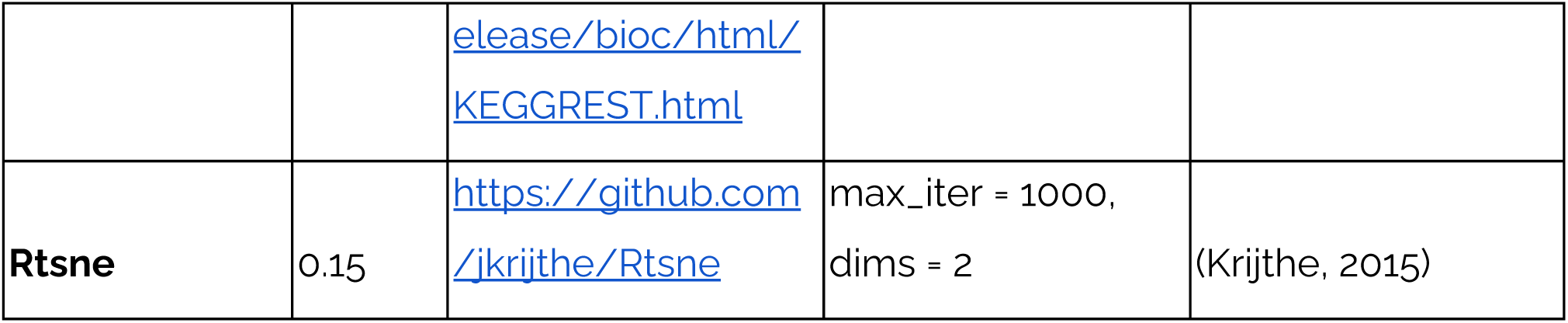
Software used.

**Table S3:** Raw data for genomic analysis of *Oscheius tipulae*.

| **Accession** | **Type** | **Platform** | **Read metrics** | **Use** |
| --- | --- | --- | --- | --- |
| **SRR12179520** | Genomic DNA | Oxford Nanopore PromethION | 108.4 Gb in 8.83 M reads with an N50 of 19.1 kb and the longest read measuring 166.0 kb | Primary genome assembly |
| **ERX1726786** | Genomic DNA | Illumina | 300 bp, paired end from 400 bp insert library* | Polishing |
| **ERX1726787** | Genomic DNA | Illumina | 100 bp, paired end from 3 kb mate pair library* | Polishing |
| **ERX1941409** | RNA | 454 GS FLX Titanium | 288.4 Mb in 610.0 K reads | Gene finding |

\* raw data from Besnard, Fabrice, et al. (2017).

**Table S4.**
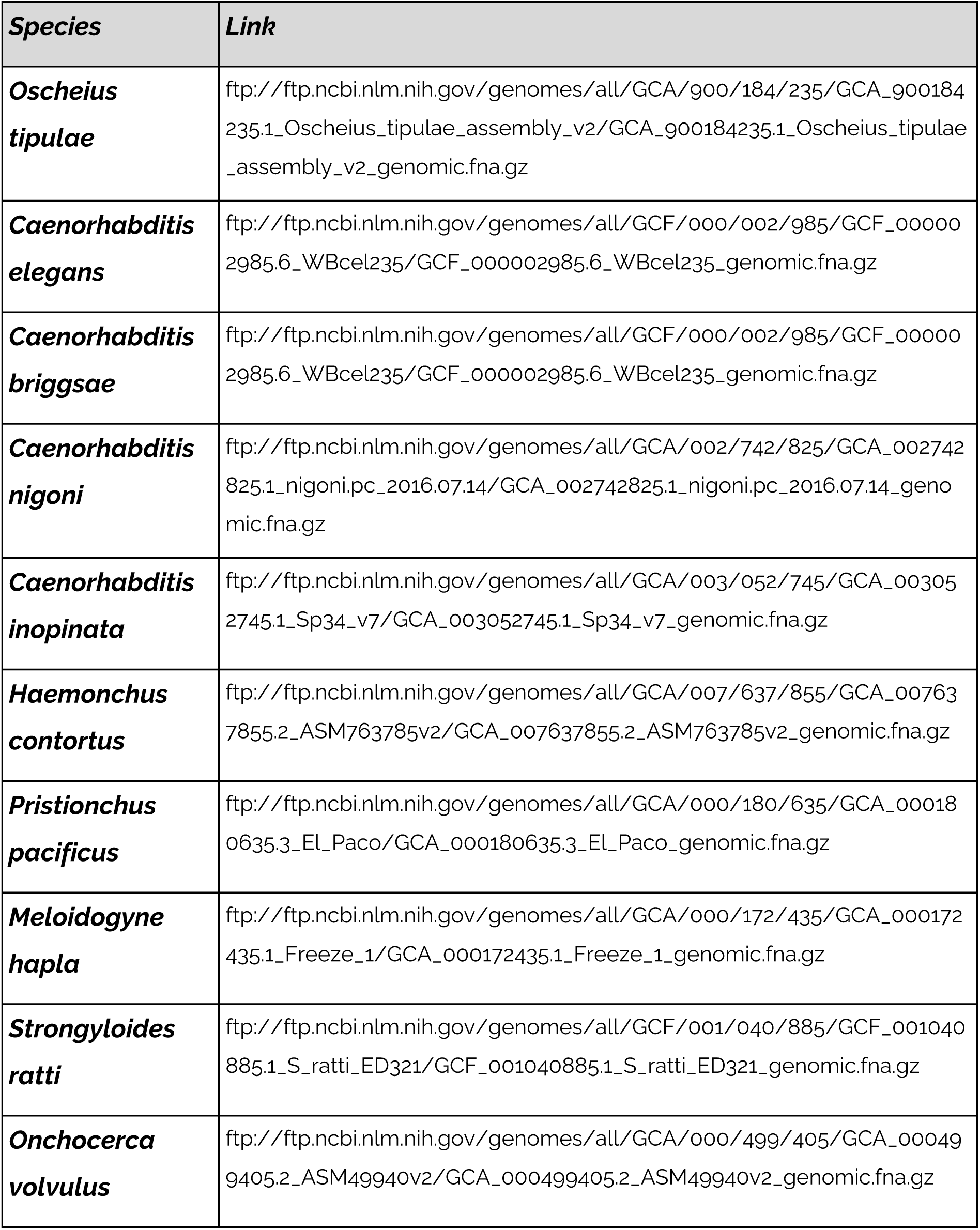
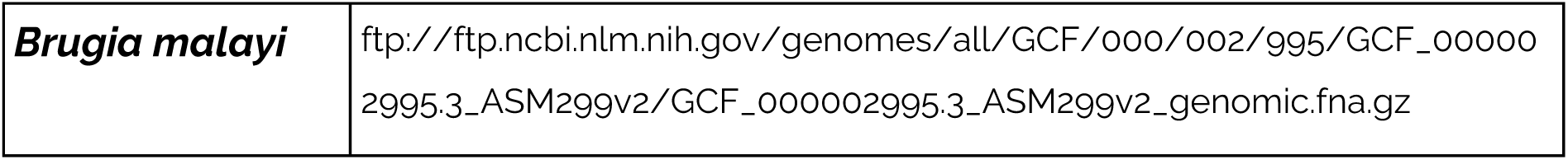
Sources of nematode assemblies included in the Kraken2 database for contamination assessment.

**Table S5.**
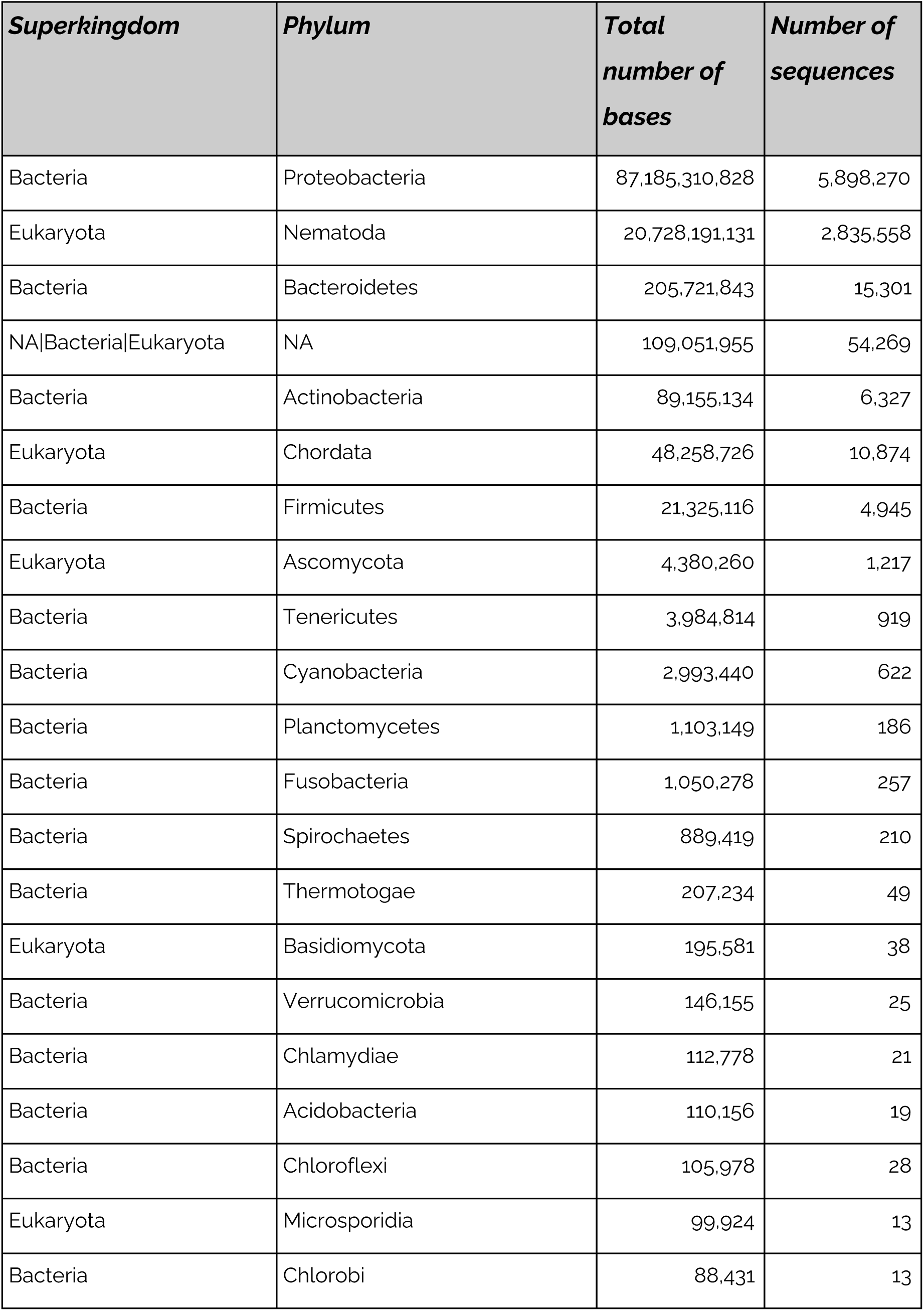
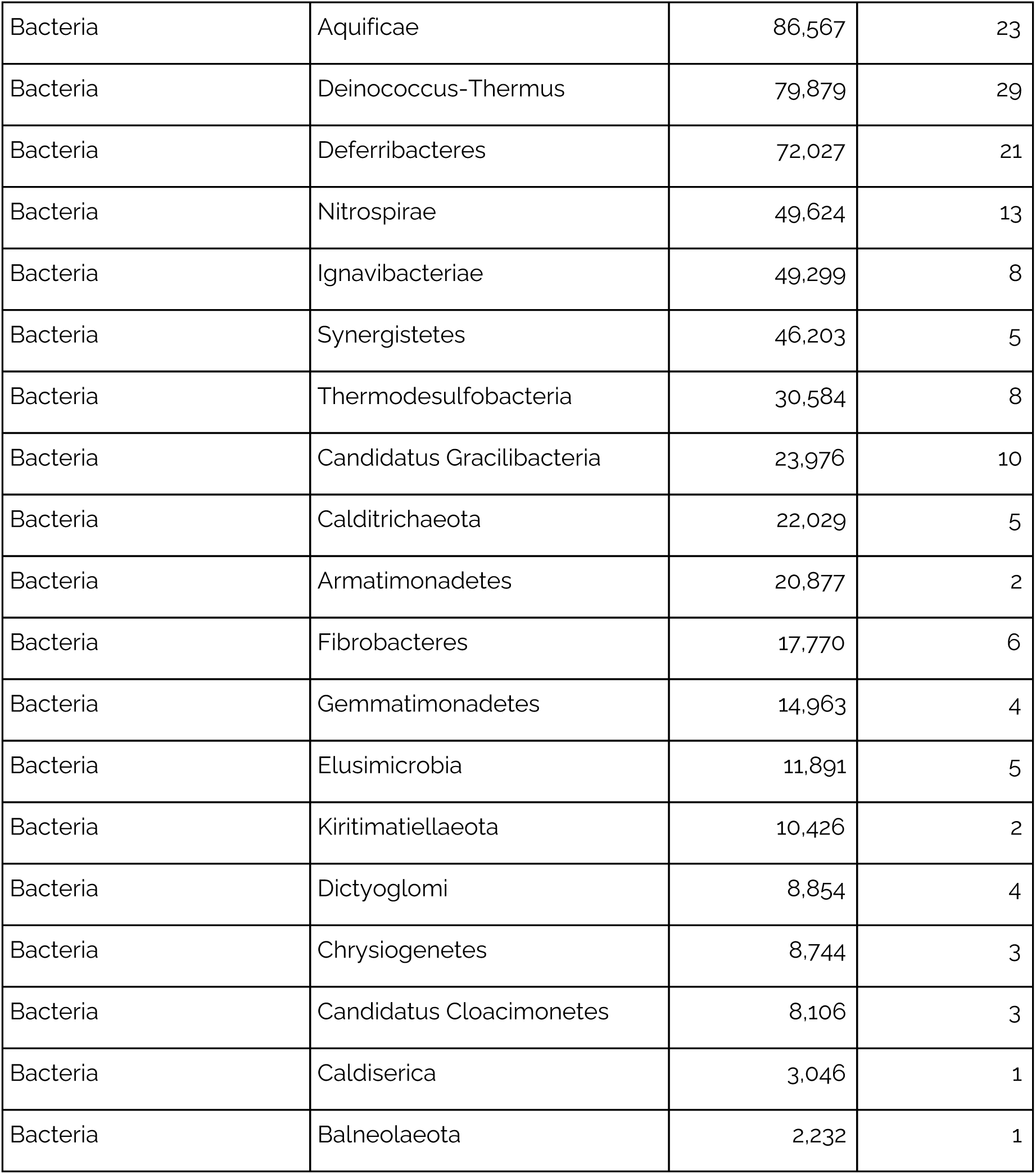
Raw sequence allocation by Kraken grouped by Phylum.

| **Superkingdom** | **Phylum** | **Total number of bases** | **Number of sequences** |
| --- | --- | --- | --- |
| Bacteria | Proteobacteria | 87,185,310,828 | 5,898,270 |
| Eukaryota | Nematoda | 20,728,191,131 | 2,835,558 |
| Bacteria | Bacteroidetes | 205,721,843 | 15,301 |
| NA\|Bacteria\|Eukaryota | NA | 109,051,955 | 54,269 |
| Bacteria | Actinobacteria | 89,155,134 | 6,327 |
| Eukaryota | Chordata | 48,258,726 | 10,874 |
| Bacteria | Firmicutes | 21,325,116 | 4,945 |
| Eukaryota | Ascomycota | 4,380,260 | 1,217 |
| Bacteria | Tenericutes | 3,984,814 | 919 |
| Bacteria | Cyanobacteria | 2,993,440 | 622 |
| Bacteria | Planctomycetes | 1,103,149 | 186 |
| Bacteria | Fusobacteria | 1,050,278 | 257 |
| Bacteria | Spirochaetes | 889,419 | 210 |
| Bacteria | Thermotogae | 207,234 | 49 |
| Eukaryota | Basidiomycota | 195,581 | 38 |
| Bacteria | Verrucomicrobia | 146,155 | 25 |
| Bacteria | Chlamydiae | 112,778 | 21 |
| Bacteria | Acidobacteria | 110,156 | 19 |
| Bacteria | Chloroflexi | 105,978 | 28 |
| Eukaryota | Microsporidia | 99,924 | 13 |
| Bacteria | Chlorobi | 88,431 | 13 |
| Bacteria | Aquificae | 86,567 | 23 |
| Bacteria | Deinococcus-Thermus | 79,879 | 29 |
| Bacteria | Deferribacteres | 72,027 | 21 |
| Bacteria | Nitrospirae | 49,624 | 13 |
| Bacteria | Ignavibacteriae | 49,299 | 8 |
| Bacteria | Synergistetes | 46,203 | 5 |
| Bacteria | Thermodesulfobacteria | 30,584 | 8 |
| Bacteria | Candidatus Gracilibacteria | 23,976 | 10 |
| Bacteria | Calditrichaeota | 22,029 | 5 |
| Bacteria | Armatimonadetes | 20,877 | 2 |
| Bacteria | Fibrobacteres | 17,770 | 6 |
| Bacteria | Gemmatimonadetes | 14,963 | 4 |
| Bacteria | Elusimicrobia | 11,891 | 5 |
| Bacteria | Kiritimatiellaeota | 10,426 | 2 |
| Bacteria | Dictyoglomi | 8,854 | 4 |
| Bacteria | Chrysiogenetes | 8,744 | 3 |
| Bacteria | Candidatus Cloacimonetes | 8,106 | 3 |
| Bacteria | Caldiserica | 3,046 | 1 |
| Bacteria | Balneolaeota | 2,232 | 1 |

**Table S6.** Contiguity and BUSCO metrics of Oscheius tipulae assemblies.

| **Read set** | **Assembler** | **Genome contiguity metrics\*** |  |  |  |  | **BUSCO\*\* scores** |  |  |  |
| --- | --- | --- | --- | --- | --- | --- | --- | --- | --- | --- |
|  |  | **#**<br>**contigs** | **Largest**<br>**contig** | **Total**<br>**length** | **N75** | **L75** | **C\*** | **D\*** | **F\*** | **M\*** |
| raw | flyemeta<sup>§</sup> | 7 | 11.30 | 60.11 | 9.02 | 5 | 88.4 | 2 | 2.1 | 9.5 |
|  | redbean | 945 | 0.59 | 47.46 | 0.04 | 323 | 62.3 | 1.9 | 1.9 | 35.8 |
|  | shasta | 55 | 5.66 | 59.83 | 1.74 | 12 | 89 | 2.3 | 2.1 | 8.8 |
| de-contaminated | flyemeta | 11 | 11.30 | 60.04 | 9.01 | 5 | 88.7 | 2 | 2.2 | 9.1 |
|  | flye | 7 | 11.24 | 60.00 | 9.01 | 5 | 88.9 | 2 | 2.1 | 9 |
|  | redbean | 19 | 11.17 | 59.63 | 8.82 | 5 | 88.7 | 2 | 2.2 | 9.1 |
|  | shasta | 34 | 9.99 | 59.47 | 2.26 | 9 | 89.2 | 2.5 | 2 | 8.8 |
|  | canu | 13 | 11.28 | 59.94 | 8.83 | 5 | 88.6 | 2.3 | 2.1 | 9.3 |
| 40x error corrected | flyemeta<sup>†</sup> | 13 | 11.18 | 59.87 | 5.13 | 6 | 89.1 | 2.1 | 2.1 | 8.8 |
|  | flye | 7 | 11.29 | 60.00 | 9.03 | 5 | 88.4 | 1.9 | 2.2 | 9.4 |
|  | redbean | 29 | 11.17 | 59.09 | 2.58 | 9 | 88.5 | 2.1 | 2.2 | 9.3 |
|  | shasta | 964 | 0.26 | 52.46 | 0.05 | 509 | 75.2 | 11.7 | 2 | 22.7 |

\* Largest contig, Total length and N75 in megabases.
\*\* BUSCO v4.0.2 with AUGUSTUS v3.3.3 scores of Nematoda\_odb10 (3131 loci); C = complete, S = single copy; D = duplicated; F = fragmented; M = missing.
<sup>§</sup> Assembly used to generate nOti 3.1.
<sup>†</sup> Assembly used to generate nOti 3.2 through a consensus with nOti 3.1.

**Table S7.**
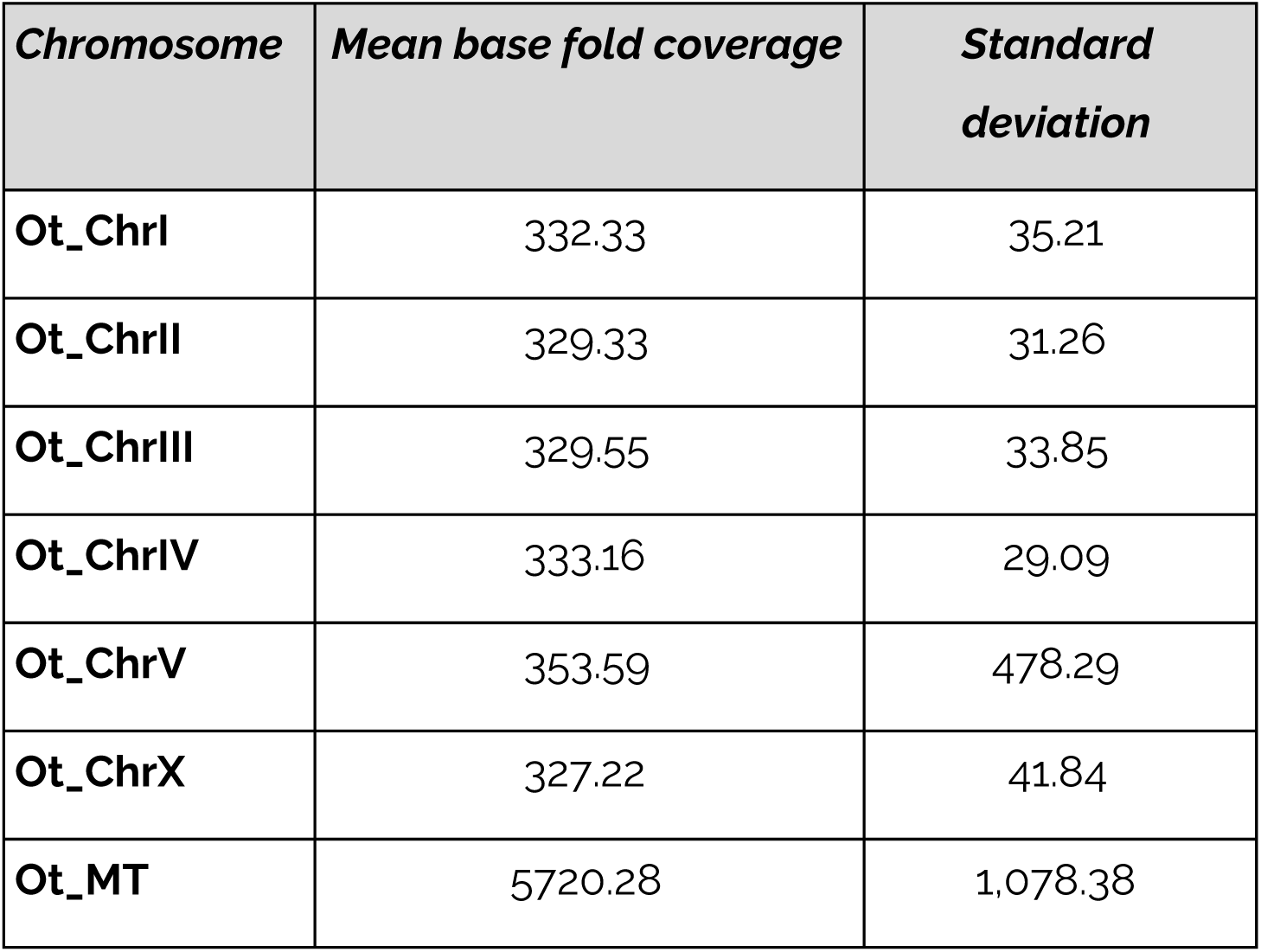
Base fold coverage of the *Oscheius tipulae* assembly.

**Table S8:** Sources and metrics of rhabditid nematode genome data.

| **Species** | **Clade** | **Reference** | **Genome size (Mb)** | **Karyotype** |
| --- | --- | --- | --- | --- |
| *Oscheius tipulae* | V | This work | 60.9 | 6 |
| *Auanema rhodensis* | V | (Tandonnet et al., 2019) | 60.6 | 7 |
| *Caenorhabditis elegans* | V | (C. elegans Sequencing Consortium, 1998) | 100.3 | 6 |
| *Caenorhabditis briggsae* | V | (Stein et al., 2003) | 108.4 | 6 |
| *Caenorhabditis nigoni* | V | (Yin et al., 2018) | 129.5 | 6 |
| *Caenorhabditis remanei* | V | (Teterina et al., 2020) | 124.8 | 6 |
| *Caenorhabditis inopinata* | V | (Kanzaki et al., 2018) | 123.0 | 6 |
| *Haemonchus contortus* | V | (Laing et al., 2016) | 283.4 | 6 |
| *Pristionchus pacificus* | V | (Rödelsperger et al., 2017) | 158.5 | 6 |
| *Steinernema carpocapsae* | IV | (Serra et al., 2019) | 84.5 | 5 |
| *Meloidogyne hapla* | IV | (Opperman et al., 2008) | 53.0 | 17 |
| *Strongyloides ratti* | IV | (Nemetschke et al., 2010) | 43.2 | 3 |
| *Ascaris suum* | III | (Wang et al., 2020) | 278.6 | 24 |
| *Onchocerca volvulus* | III | (Cotton et al., 2016) | 96.4 | 5 |
| *Brugia malayi* | III | (Foster et al., 2020) | 88.2 | 6 |

**Table S9.** BUSCO scores of assemblies and proteomes of rhabditid nematodes.

| Species | BUSCO assembly*<sup>§</sup> |  |  |  |  | BUSCO* proteome |  |  |  |  | # of protein coding genes |
| --- | --- | --- | --- | --- | --- | --- | --- | --- | --- | --- | --- |
|  | C | S | D | F | M | C | S | D | F | M |  |
| *Oscheius tipulae* (v3.0) | 90.7 | 88.7 | 2 | 1.8 | 7.5 | 90.7 | 87.9 | 2.8 | 2.4 | 6.9 | 16138 |
| *Oscheius tipulae* (nOt2.0) | 91.6 | 89 | 2.6 | 1.6 | 6.8 | 89.1 | 86.2 | 2.9 | 2.0 | 8.9 | 14626 |
| *Ascaris suum* | 94.7 | 91.7 | 3 | 1.2 | 4.1 | 79.9 | 77.4 | 2.5 | 3.4 | 16.7 | 17969 |
| *Auanema rhodensis* | 89.9 | 89.3 | 0.5 | 1.4 | 8.7 | 85.4 | 84.6 | 0.8 | 1.2 | 13.4 | 10861 |
| *Caenorhabditis elegans* | 99.5 | 98.9 | 0.5 | 0.1 | 0.5 | 100.0 | 99.6 | 0.4 | 0.0 | 0.0 | 20184 |
| *Caenorhabditis briggsae* | 99 | 98.4 | 0.6 | 0.4 | 0.6 | 99.2 | 98.5 | 0.7 | 0.4 | 0.4 | 20829 |
| *Caenorhabditis nigoni* | 99.4 | 98.6 | 0.8 | 0.1 | 0.5 | 96.8 | 95.7 | 1.1 | 1.4 | 1.8 | 29167 |
| *Caenorhabditis remanei* | 99.3 | 98.1 | 1.1 | 0.3 | 0.5 | 93.1 | 91.9 | 1.2 | 0.9 | 6.0 | 26308 |
| *Caenorhabditis inopinata* | 98.8 | 97.9 | 0.9 | 0.4 | 0.8 | 98.7 | 97.7 | 1.0 | 0.2 | 1.1 | 21443 |
| *Haemonchus contortus* | 94.4 | 93.3 | 1.1 | 0.9 | 4.7 | 93.1 | 91.6 | 1.5 | 1.1 | 5.8 | 19473 |
| *Pristionchus pacificus* | 82.6 | 80.9 | 1.7 | 1.5 | 15.9 | 77.4 | 75.2 | 2.2 | 1.0 | 21.6 | 25992 |
| *Steinernema carpocapsae* | 78.2 | 74.8 | 3.5 | 3 | 18.7 | 74.7 | 70.9 | 3.8 | 3.8 | 21.5 | 30825 |
| *Meloidogyne hapla* | 59.4 | 58.3 | 1.1 | 2.3 | 38.3 | 52.6 | 51.1 | 1.4 | 2.6 | 44.8 | 14419 |
| *Strongyloides ratti* | 68 | 66.4 | 1.6 | 1.5 | 30.5 | 70.6 | 67.8 | 2.8 | 0.9 | 28.5 | 12464 |
| *Onchocerca volvulus* | 98 | 97.4 | 0.6 | 0.4 | 1.6 | 98.3 | 97.2 | 1.1 | 0.5 | 1.2 | 12110 |
| *Brugia malayi* | 99.1 | 98.4 | 0.7 | 0.2 | 0.8 | 98.9 | 98.3 | 0.6 | 0.2 | 0.9 | 10940 |

\* BUSCO v4.0.4 with AUGUSTUS v3.2.3 scores of Nematoda\_odb10 (3131 loci); C = complete, S = single copy; D = duplicated; F = fragmented; M = missing. For genome mode
<sup>§</sup>Using flag for Augustus self optimization

**Table S10:** Nigon element-defining loci in fifteen nematode genomes.

| **Nigon** | **A** | **B** | **C** | **D** | **E** | **N** | **X** | **total** | **Proportion** |
| --- | --- | --- | --- | --- | --- | --- | --- | --- | --- |
| *Ascaris suum* | 477 | 304 | 462 | 254 | 224 | 162 | 114 | 1997 | 0.91 |
| *Auanema rhodensis* | 521 | 315 | 495 | 272 | 227 | 169 | 100 | 2099 | 0.96 |
| *Brugia malayi* | 530 | 324 | 516 | 271 | 239 | 171 | 119 | 2170 | 0.99 |
| *Caenorhabditis briggsae* | 518 | 317 | 504 | 269 | 235 | 173 | 119 | 2135 | 0.97 |
| *Caenorhabditis elegans* | 529 | 326 | 517 | 272 | 240 | 172 | 119 | 2175 | 0.99 |
| *Caenorhabditis nigoni* | 505 | 315 | 491 | 247 | 227 | 166 | 109 | 2060 | 0.94 |
| *Caenorhabditis remanei* | 491 | 306 | 482 | 242 | 220 | 166 | 115 | 2022 | 0.92 |
| *Caenorhabditis inopinata* | 526 | 318 | 507 | 271 | 239 | 172 | 119 | 2152 | 0.98 |
| *Haemonchus contortus* | 520 | 317 | 509 | 271 | 236 | 160 | 108 | 2121 | 0.97 |
| *Meloidogyne hapla* | 429 | 257 | 419 | 210 | 170 | 122 | 81 | 1688 | 0.77 |
| *Onchocerca volvulus* | 526 | 322 | 510 | 273 | 234 | 170 | 117 | 2152 | 0.98 |
| *Osccheius tipulae* | 516 | 320 | 501 | 269 | 216 | 173 | 118 | 2113 | 0.96 |
| *Pristionchus pacificus* | 453 | 297 | 466 | 248 | 223 | 163 | 113 | 1963 | 0.9 |
| *Steinernema carpocapsae* | 497 | 291 | 474 | 258 | 231 | 156 | 112 | 2019 | 0.92 |
| *Strongyloides ratti* | 509 | 309 | 484 | 252 | 234 | 170 | 114 | 2072 | 0.95 |
| **Total loci per Nigon** | **534** | **328** | **519** | **275** | **242** | **174** | **119** | **2191** | **1.00** |

**Table S11.** KEGG categories enriched among Nigon defining loci.

| ***Nigo***<br>***n*** | ***KEGG term*** | ***adjusted_p_val***<br>***ue*** |
| --- | --- | --- |
| A | Ribosome biogenesis in eukaryotes | $1.91 \times 10^{-04}$ |
| A | RNA transport | $1.37 \times 10^{-02}$ |
| C | Spliceosome | $1.87 \times 10^{-02}$ |
| C | RNA transport | $2.42 \times 10^{-02}$ |
| D | Calcium signaling pathway | $1.14 \times 10^{-02}$ |
| D | ErbB signaling pathway | $3.66 \times 10^{-02}$ |
| N | Hippo signaling pathway - multiple species | $2.38 \times 10^{-03}$ |
| X | Axon regeneration | $1.15 \times 10^{-05}$ |
| X | Calcium signaling pathway | $1.46 \times 10^{-02}$ |
| X | Neuroactive ligand-receptor interaction | $1.56 \times 10^{-02}$ |

**Table S12:**
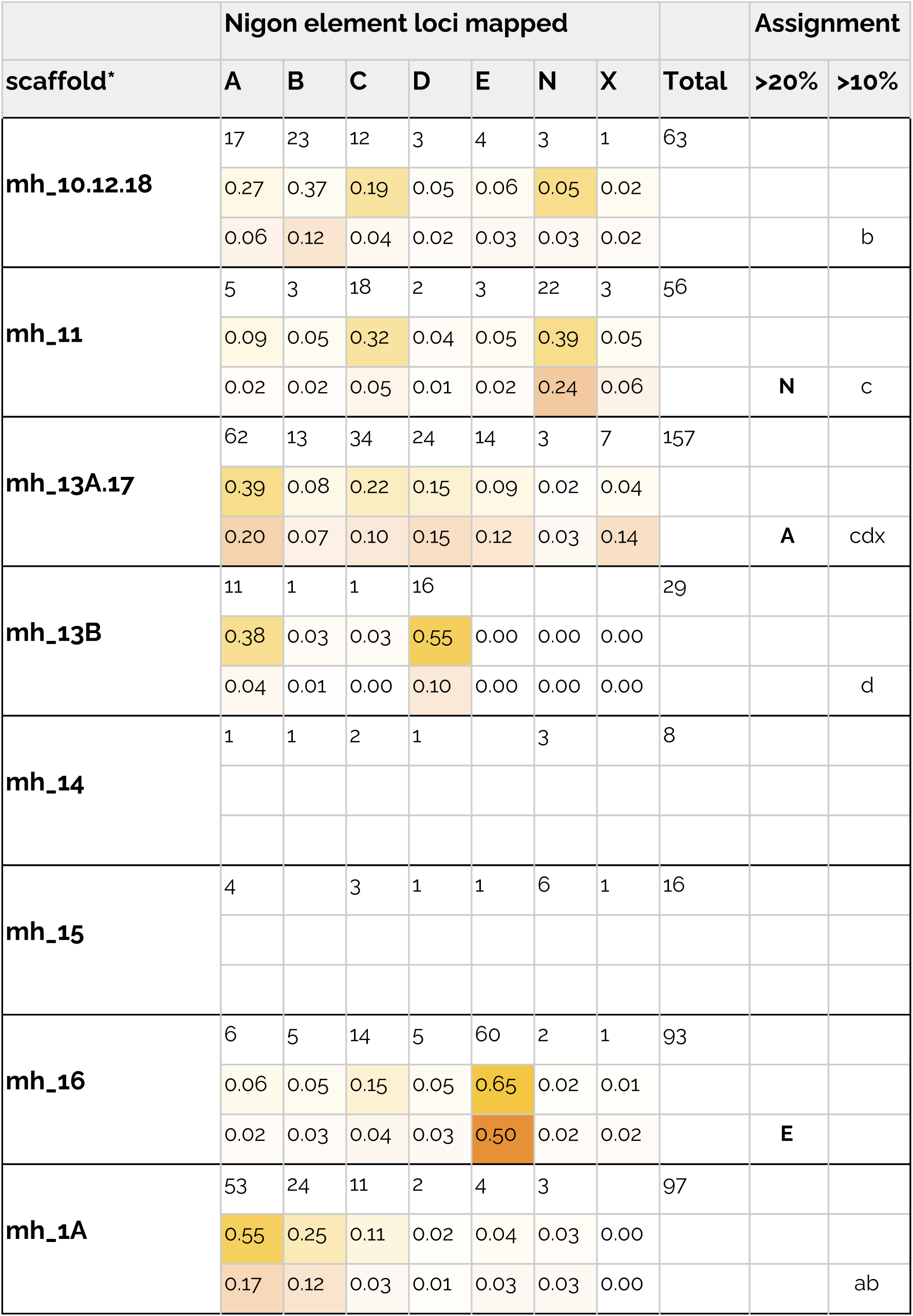
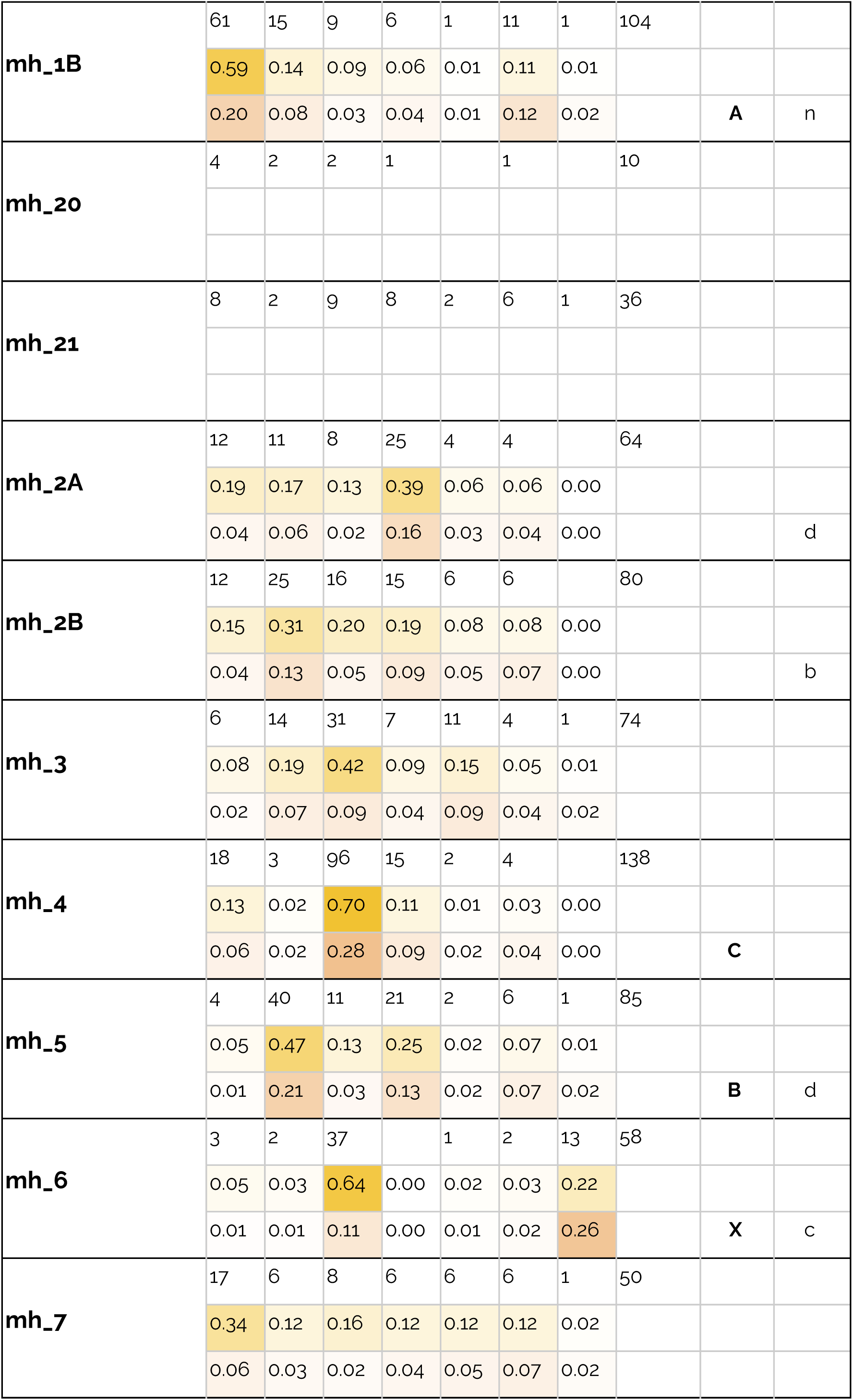
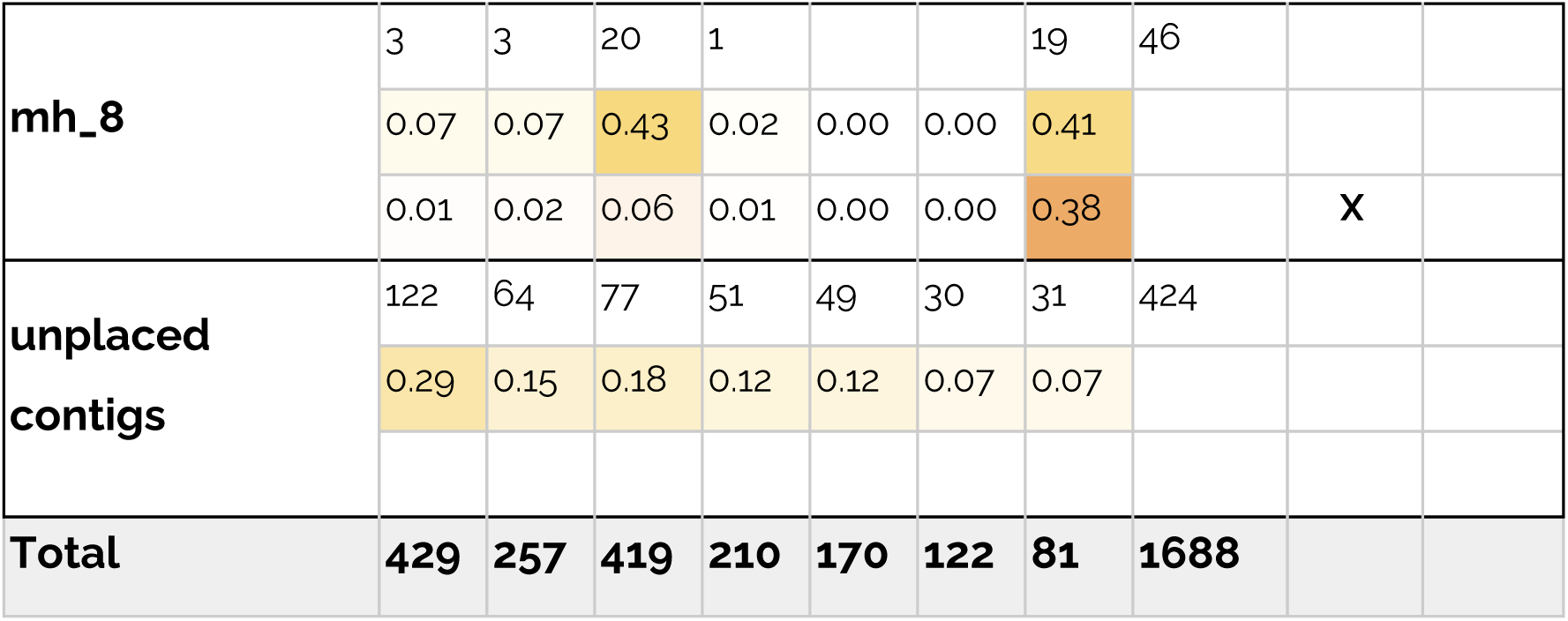
Nigon-defining loci in the Meloidogyne hapla genome assembly.

## Online data and code

A github repository containing code for the R analyses of genome features and the circos plot data and configuration files is at https://github.com/tolkit/otipu_chrom_assem.

A data sheet containing data for tables and figures is at <u>A Chromosomal assembly of Oscheius_tipulae data.</u>

A presentation file of the figures is at The Oscheius tipulae genome Figures for Paper.

